# Insertion of an invading retrovirus regulates a novel color trait in swordtail fish

**DOI:** 10.1101/2025.11.07.687308

**Authors:** Nadia B. Haghani, Tristram O. Dodge, John J. Baczenas, Theresa R. Gunn, Qinliu He, William Pangburn, Rhea Sood, Paola Fascinetto-Zago, Kang Du, Alyssa Kaatmann, Zihao Ou, Sashoya Dougan, Terrance TL Yang, Natalie S. Olmos-Santiago, Gabriel A. Preising, Cheyenne Y. Payne, Sophia K. Haase Cox, Kelsie E. Hunnicutt, Max Madrzyk, Benjamin Moran, Christian Stigloher, Daniel L. Powell, Manfred Schartl, Casey A. Gifford, Guan-Zhu Han, Molly Schumer

## Abstract

For over a century, evolutionary biologists have been motivated to understand the mechanisms through which organisms adapt to their environments. Coloration and pigmentation are remarkably variable within and between species and can serve as an important window into the mechanisms of adaptation. Here, we map the genetic basis of a newly described iridescence trait in swordtail fish to a single locus. Individuals with this trait appear to sparkle as they move through the water. We find that the trait is driven by the recent endogenization of a retrovirus that inserted near the gene *alkal2a*. This insertion is associated with changes in the chromatin landscape, upregulation of *alkal2a*, and accumulation of iridescent cells that adhere to the scales. Rather than causing diseases, our results demonstrate that invading endogenous retroviruses can also regulate novel trait variation in the host. Moreover, we find that this coloration trait may act as an important signal in interactions between fish and their predators in the natural environment.

## Introduction

The diversity of colors and patterns across the tree of life is extraordinary and can evolve rapidly within and between species. Many organisms rely on coloration traits to survive, reproduce, and communicate in their natural environments. Thus, dissecting the genetic and evolutionary mechanisms that produce variation in color can help us understand how organisms adapt to their environments. While pigment molecules are commonly used to produce color, some organisms also use nanostructures to manipulate light in ways that generate the appearance of color, a phenomenon known as structural coloration (*1*, *2*). Structural colors have convergently evolved in many lineages. In vertebrates, structural colors are often associated with specialized cell types called iridophores, which contain subcellular crystals. Yet, the genetic mechanisms and evolutionary consequences of structural colors are understudied compared to pigment-based coloration traits.

Decades of work have highlighted the importance of changes in gene regulation as a mechanism of evolutionary novelty, including in pigmentation phenotypes (*3–8*). How gene regulatory networks evolve and gain new functions has been the subject of intensive research (*9–13*). One mechanism that has emerged as an important driver of lineage and tissue specific gene regulation is co-option of transposable elements (TEs) by the host genome (*14*). TEs can alter gene expression through various mechanisms, including directly disrupting host promoters, enhancers, or repressors (*15–18*), by driving structural rearrangements (e.g. by disrupting a TAD boundary; *19*), or by triggering host repression pathways (*20*). Certain classes of TEs, namely retrotransposons, contain their own promoters and enhancers that can directly elicit changes in host gene expression (*21*, *22*). Through their impacts on gene regulation, TEs can facilitate the evolution of adaptive traits, as has been demonstrated for insecticide resistance in flies, pigmentation in mammals and pepper moths, among others (*23–25*). Notably, the number and characteristics of TE families vary widely across taxa (*26–28*). Among vertebrates, teleost fish have remarkable diversity in the types of TE families they harbor and an unusually elevated number of active or recently active TEs, making them a powerful system to study the links between gene regulation and TE activity (*27, 29, 30, 31, 32*).

Retroviruses (the *Retroviridae* family) exclusively infect vertebrates and can cause a variety of diseases. Their replication requires integration into the host genome (*33*, *34*). When the infection occurs in germline cells, integrated retroviruses can be vertically inherited, giving rise to so-called endogenous retroviruses (ERVs; *33*, *34*). At the time of insertion, ERVs encode for all the necessary machinery and regulatory sequences to promote their own transcription and retrotransposition. Therefore, retroviruses possess features characteristic of both TEs and viruses. ERVs are ubiquitously distributed in vertebrate genomes, but most of them are no longer active due to the accumulation of disruptive mutations (*35*, *36*). The long terminal repeats (LTRs) of ERVs act as enhancers and promoters for the viral genome, and as a result, ERV insertions frequently have intrinsic regulatory activity. As a consequence, LTRs are often co-opted by the host as *cis*-regulatory elements, and over evolutionary timescales, have become essential for host gene regulation and normal development (*21*, *22*). While infrequently observed, researchers have also identified a number of retroviruses that were recently active (i.e. capable of transposition in the host genome; *37*) or are actively invading their host genomes. Such cases of active invasion can provide powerful models to study the evolutionary trajectories and organismal consequences of active TEs in real-time (*38*, *39*). However, there are few known cases of active retrovirus endogenization, with the best documented case linked to disease states in Koalas (*38*, *40*), leaving many open questions about this key evolutionary transition.

Here, we describe an active or recently active ERV and show that it regulates color variation in swordtail fish. Swordtail fishes (genus *Xiphophorus*) are a tractable lab model for genetic and evolutionary studies, harboring a diverse array of heritable coloration traits (Fig. 1A). We characterize a unique structural color pattern found in a natural population of swordtail fish and show that it is generated at the cellular level by the aggregation of iridophore cells on the scales. These cells interact with light to produce the appearance of a silver color. We map this structural coloration trait to a single genomic region containing the gene *alkal2a*, which encodes a cytokine known to regulate iridophore development and maintenance (*41–43*). We find that an evolutionarily recent, full-length insertion of a novel ERV drives upregulation of the *alkal2a* gene, leading to the development of iridescence. Population genetic analysis of long-read genomes indicates that this ERV is either actively replicating or was replicating very recently in the swordtail genome. We demonstrate that fish with this iridescence trait produce distinct visual signals which could impact predator evasion in their natural populations.

**Fig. 1:**
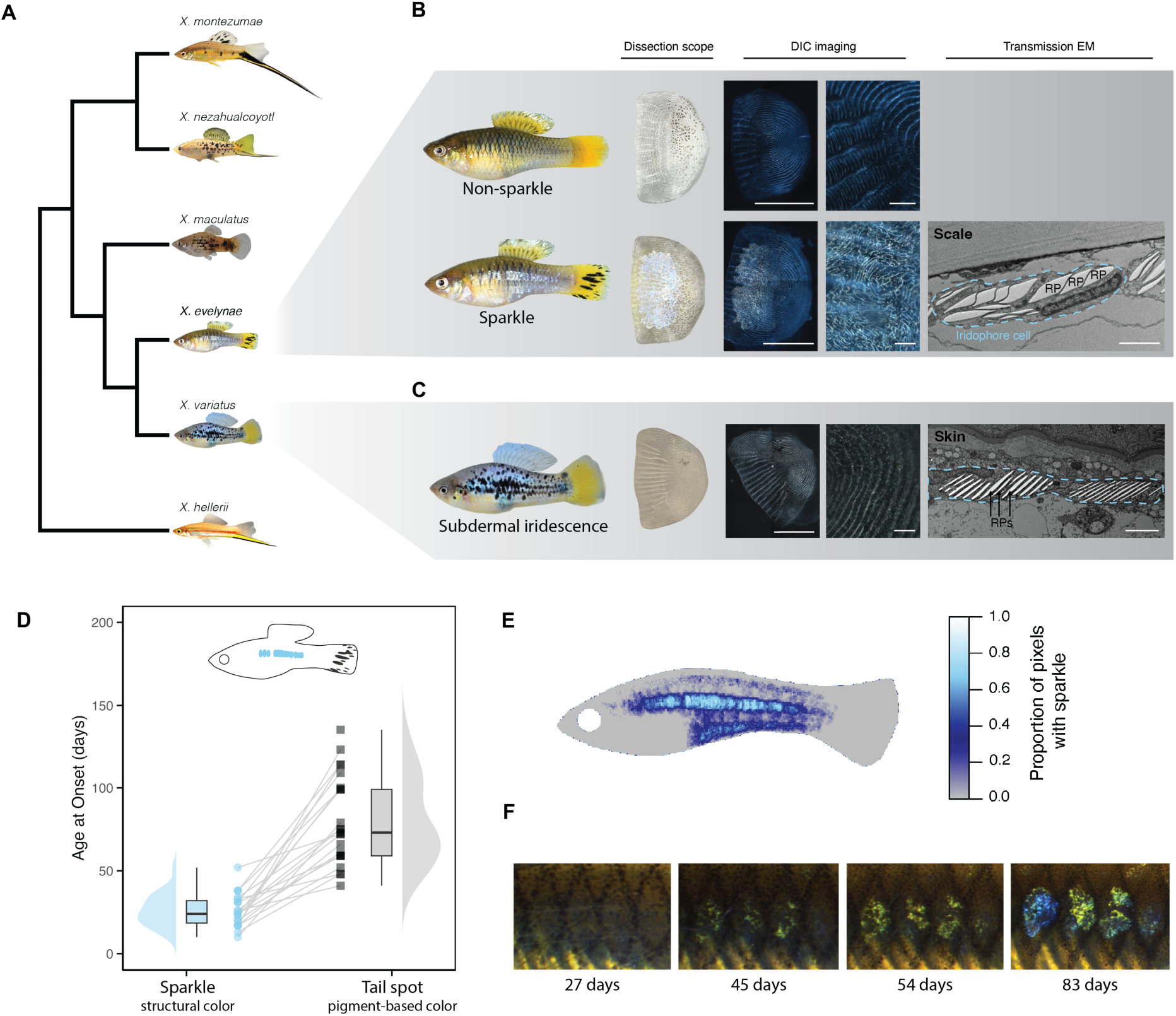
Observed pigmentation diversity and patterns in *Xiphophorus* and focal species. **A)** *Xiphophorus* fishes show immense variability in pigmentation patterns both within and between species. Here we show a simplified phylogeny (see Fig. S1 for full phylogeny), highlighting phenotypic variation in melanophore, xanthophore, and iridophore-based pigmentation patterns across different groups of *Xiphophorus* fishes. Gray triangle insets highlight focal species in this study. **B)** Natural populations of *Xiphophorus evelynae* fish segregate for a structural color polymorphism, which we refer to as the ‘sparkle’ trait. From left to right: Unlike non-sparkle individuals (top), fish with sparkle (bottom) have scales that reflect light. Dissection scope images of individually isolated scales from either non-sparkle or sparkle fish. Differential interference contrast (DIC) imaging reveals scale-adherent cells in sparkle individuals; scale bar = 1000 µm. Magnified DIC images highlight light-reflecting microstructures in scales from individuals with the sparkle trait. Note the absence of this cell population in non-sparkle scale images; scale bar = 100 µm. Transmission electron microscope (EM) images confirm the presence of iridophore cells adjacent to the scale, with typical reflective platelet (RP) microstructures present inside iridophore cells; scale bar = 2 µm. The dashed blue line highlights the outline of an iridophore cell. **C)** While the pattern of scale-associated iridescence in *X. evelynae* is unique, many species of *Xiphophorus*, including the *X. variatus* individual shown here, display subdermal iridescence. Scale images taken with a dissection microscope and DIC indicate that the cell population present on the scales of *X. evelynae* is absent in *X. variatus*. TEM image shows skin iridophores from *X. variatus*. **D)** Development of the sparkle trait (blue circles) occurs earlier than a common melanophore-based coloration trait in *X. evelynae* (i.e. tail spots; gray squares). Lines connect individual fish. **E)** Heatmap of sparkle trait location across 30 wild-caught, adult individuals fitted to a standard body outline (see Fig. S23 for landmarks). **F)** Example of a fish imaged over development (27, 45, 54, 83 days). Iridescence gradually increases over time until it fills the entire scale.

## Results

### Phenotypic variation in a structural color trait, ‘sparkle’

Sampling from rivers near the town of Juntas Chicas in Veracruz México, we identified a previously unstudied population of *Xiphophorus* closely related to the species *X. evelynae* (Fig. S1-S2; Supplementary Materials 1; hereafter *X. evelynae*). Fish sampled from this population varied in the presence of a striking coloration trait (Fig. 1A-B; Fig. S3). Approximately a quarter (25.4 ± 2.7%) of individuals collected from this population had reflective scales along the body, a trait which we refer to as ‘sparkle’ (see also Supplementary Materials 1; *44*, *45*). This color phenotype originates from the scales, in contrast to iridescence phenotypes found in other swordtail species where the signal localizes to deeper epithelial layers (Fig. 1A-C). Differential interference contrast (DIC) imaging revealed a mass of iridescent cells adhered to the scales of sparkle individuals (Fig. 1B; Supplementary Materials 2). These densely packed cells morphologically resembled iridophores (Fig. 1B-C, Fig. 2F). Iridophores commonly cause iridescence in fish via interactions between light sources and the reflective platelets embedded inside the cells (*1*, *46–48*). To evaluate whether the sparkle trait was generated by similar mechanisms, we conducted Transmission Electron Microscopy (TEM) of sparkle scales. As expected, we identified the presence of stacked, reflective platelets—microstructures inside of iridophore cells capable of producing structural coloration traits (Fig. 1B; Supplementary Materials 2). Intriguingly, the shape of the reflective platelets in the iridophores of *X. evelynae* differed from those observed in iridophores embedded in the epidermis of other swordtail species (Fig. 1B-C). The microstructures in the epidermis of other *Xiphophorus* species more closely resemble the reflective platelets within iridophores previously characterized in zebrafish (*1*, *49–51*).

**Fig. 2:**
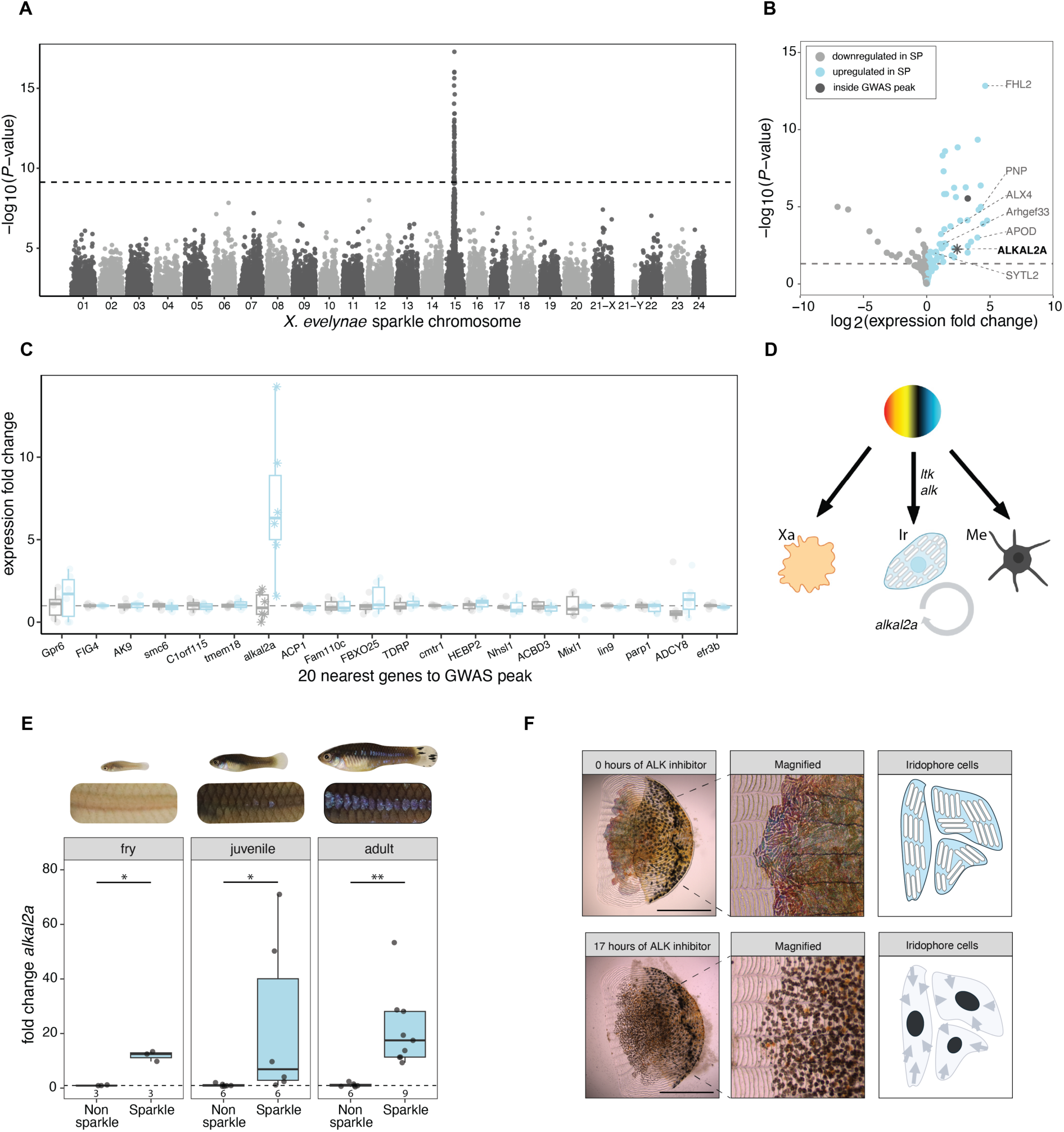
Variation in the sparkle phenotype maps to a single region containing *alkal2a*, an iridophore differentiation gene. **A)** Case-control genome-wide association study (GWAS) on wild-caught non-sparkle (n=194) and sparkle (n=66) individuals. Dashed line indicates the genome-wide significance threshold for allele frequency differences between non-sparkle and sparkle individuals (determined using permutations; p<7.4×10^−10^). **B)** Volcano plot displaying genes either upregulated (blue points) or downregulated (grey points) in sparkle scales relative to non-sparkle scales from fish lacking the trait. Dark grey points represent genes that fall inside the GWAS peak. Points above the dashed grey line have adjusted p-values <0.05. Many significantly differentially expressed genes from RNA-seq data are specific to iridophore cell types in zebrafish; these genes are labeled on the right side of the plot. **C)** Fold change in expression between sparkle and non-sparkle scales for 20 genes in the center of the GWAS region. The results here show all genes in the 400 kb region most strongly associated with the trait based on GWAS analysis. Among them, a single gene, *alkal2a*, is differentially expressed between sparkle and non-sparkle scales. **D)** Schematic of pigment cell differentiation pathway in *D. rerio* shows a pluripotent pigment progenitor cell that can differentiate into iridophores upon activation of receptor tyrosine kinase Ltk/Alk by the ALKAL2A ligand. ALKAL2A is also required to maintain iridophore cell populations once they have developed. Image adapted from Patterson & Parichy (*63*). Xa=Xanthophore; Ir = Iridophore; Me=Melanophore. **E)** Results of qPCR experiments quantifying expression of the *alkal2a* gene in non-sparkle (grey) or sparkle (blue) scales taken from the midline of fry, juvenile, and adult *X. evelynae*. Fry and juveniles were genotyped to confirm that they harbored the sparkle haplotype. These results demonstrate that *alkal2a* is differentially expressed throughout development, with differential expression preceding detection of the sparkle trait. Number of fish tested per group is noted on the x-axis of each plot. Expression differences at each stage were tested using a Kruskal-Wallis test; * *P* < 0.05, ** *P* < 0.01. **F)** Left: Individual scales dissected from sparkle fish before (top) or after (bottom) treatment with an inhibitor of the ALK receptor for 17 hours. For time course images of inhibitor treatments with paired DMSO-treated controls, see Fig. S6. Anterior surface layer of the scale contains black melanophores, while the middle/posterior layer of the scale closest to the fish skin contains reflective iridophores. Middle: Magnified color imaging allows for iridophore visualization post-ALK inhibitor treatment. Right: Cartoon depiction of changes in cell morphology in scales treated with the ALK inhibitor.

Tracking the development of *X. evelynae* fish revealed that the sparkle trait begins to appear within the first two months of life and occurs before the onset of a common pigment-based trait (Fig. 1D). While the timing of trait onset and intensity of iridescence varied between individuals (Fig. S3-S4; Supplementary Materials 3-4), we found that the trait is not sexually dimorphic and that the spatial location of this patterning is highly predictable (Fig. 1E). The trait first appeared near the midline as iridophore-dense puncta that ultimately completely fill in the scale (Fig. 1F; Fig. S4; Supplementary Materials 4-4).

### Variation in the sparkle trait maps to a single genomic region containing *alkal2a*

To identify genomic regions associated with the sparkle trait, we collected low coverage whole genome sequencing data from 260 *X. evelynae* individuals from the Juntas Chicas population (Supplementary Materials 5-6) and performed a genome-wide association study (GWAS) to test for allele frequency differences along the genome between 66 individuals with the sparkle trait and 194 controls (non-sparkle). For this analysis, we generated and annotated a high-quality genome assembly for an *X. evelynae* individual with the sparkle phenotype using PacBio HiFi technology (Supplementary Materials 5). The resulting assembly was near chromosome scale and is syntenic with the genomes of other *Xiphophorus* species (Supplementary Materials 5; Table S1; Fig. S5). Our GWAS identified a single genome-wide significant region associated with the sparkle trait (Fig. 2A). We confirmed our results using a second reference genome generated from an individual without the sparkle phenotype and show that this association is robust to corrections for population structure (Supplementary Materials 6). The most strongly associated region, defined as within two -log_10_(p-value) units of the peak SNP, spanned ∼400 kb and contained 16 genes (Table S2; Supplementary Materials 6).

To determine which genes within this region were likely to be driving the GWAS signal, we examined expression differences between phenotypes. We conducted bulk RNA sequencing from scales and their adherent cells collected from sparkle and non-sparkle individuals (4 biological replicates per phenotype; Supplementary Materials 7). We found that 91 genes were differentially expressed between the two types of scales (Fig. 2B, Supplementary Materials 7). Only two differentially expressed genes, *smyd2* and *alkal2a*, fell within the GWAS region (Supplementary Materials 7). Unlike *smyd2*, *alkal2a* is a known pigmentation gene and falls within the most strongly associated GWAS region, less than 100 kb from the peak SNP (Table S2; Supplementary Materials 7; Fig. 2C). *Alkal2a* is a ligand that activates receptor tyrosine kinases of the ALK family. In zebrafish, *alkal2a* plays dual roles in driving the differentiation of pigment progenitor cells into iridophores and in maintaining iridophore cells (Fig. 2D; Supplementary Materials 7; *41*, *43*, *52*). Using a qPCR approach, we confirmed that *alkal2a* is overexpressed in sparkle scales relative to non-sparkle scales whereas other pigmentation genes in the GWAS interval are not. (Fig. 2E; Supplementary Materials 8). Moreover, pharmacological inhibition of the *alkal2a* pathway resulted in loss of morphology of iridophore cells adhered to the scale (Fig. 2F; Fig. S6; Video S1; Supplementary Materials 9). Intriguingly, *alkal2a* is overexpressed in scales taken from the midline of 3-week-old fry that had not yet developed the sparkle trait but harbored the sparkle haplotype (Fig. 2E; Supplementary Materials 8). Thus, differential expression of *alkal2a* precedes the sparkle trait development, consistent with *alkal2a* being the driver of iridophore patterning.

Our RNA-seq analysis identified dozens of genes outside the GWAS peak that are differentially expressed between sparkle and non-sparkle scales (Fig. 2B). As expected, we found that many of the genes upregulated in sparkle individuals were associated with pigment cell-specific gene expression in zebrafish. For example, differentially expressed genes include *Fhl2*, *Pnp*, *Alx4*, *Arhgef33*, *Fmn2*, *Sytl2*, and *Apod*, which are almost exclusively expressed in zebrafish iridophore and xanthophore lineages (Supplementary Materials 7). More broadly, 17% of differentially expressed genes are known to impact pigmentation phenotypes when knocked out in zebrafish and 63% impact the phenotypes of cells derived from the neural crest lineage (Supplementary Materials 7; note that the neural crest lineage gives rise to pigment progenitor cells). Given that these differentially expressed genes do not occur in the GWAS peak, we speculate that these genes may be interacting partners of *alkal2a* (or fall in other pathways).

### Haplotype resolved long-read assemblies uncover an insertion near *alkal2a*

The region identified in our genome-wide association study had unusual patterns of linkage disequilibrium (Supplementary Materials 6). To investigate whether this is driven by a structural rearrangement in this region (*53*), we used PacBio HiFi technology to sequence and infer haplotypes for a total of 3 sparkle and 3 non-sparkle individuals (Supplementary Materials 5). This resulted in the assembly of 11 distinct haplotypes for chromosome 15 (Supplementary Materials 5; Table S3). Using these data, we aligned haplotypes from each of our assemblies to examine structural differences near the *alkal2a* gene and throughout the GWAS peak (Fig. 3A; Fig. S7; Supplementary Materials 5). We identified a 17 kb insertion (Fig. 3A, blue rectangle) approximately 4 kb downstream of the 3’ UTR of *alkal2a* (Fig. 2C). The inserted sequence occurs in an intron of the *acp1* gene in the antisense orientation but does not appear to impact *acp1* expression (Fig. 2C). This insertion was present in at least one haplotype of all sparkle individuals but was absent in both haplotypes from all non-sparkle individuals (Fig. 3A). Based on inspection of a local haplotype alignment (using MAFFT; *54*), the position of the insertion appears to be shared across all sequenced individuals (Fig. S8). We note that there are several other SNPs and smaller structural variants in this region that distinguish sparkle and non-sparkle haplotypes; we discuss these regions in detail in Supplementary Materials 5 and revisit them in experiments below. PCR-based genotyping of an additional 18 sparkle and 11 non-sparkle individuals confirmed that the insertion is diagnostic of the sparkle phenotype (Supplementary Materials 8). While all individuals with the sparkle trait shared this inserted sequence, we did not detect any difference in predicted amino acid sequence between sparkle and non-sparkle individuals in *alkal2a* (Supplementary Materials 5; Fig. S9).

**Fig. 3:**
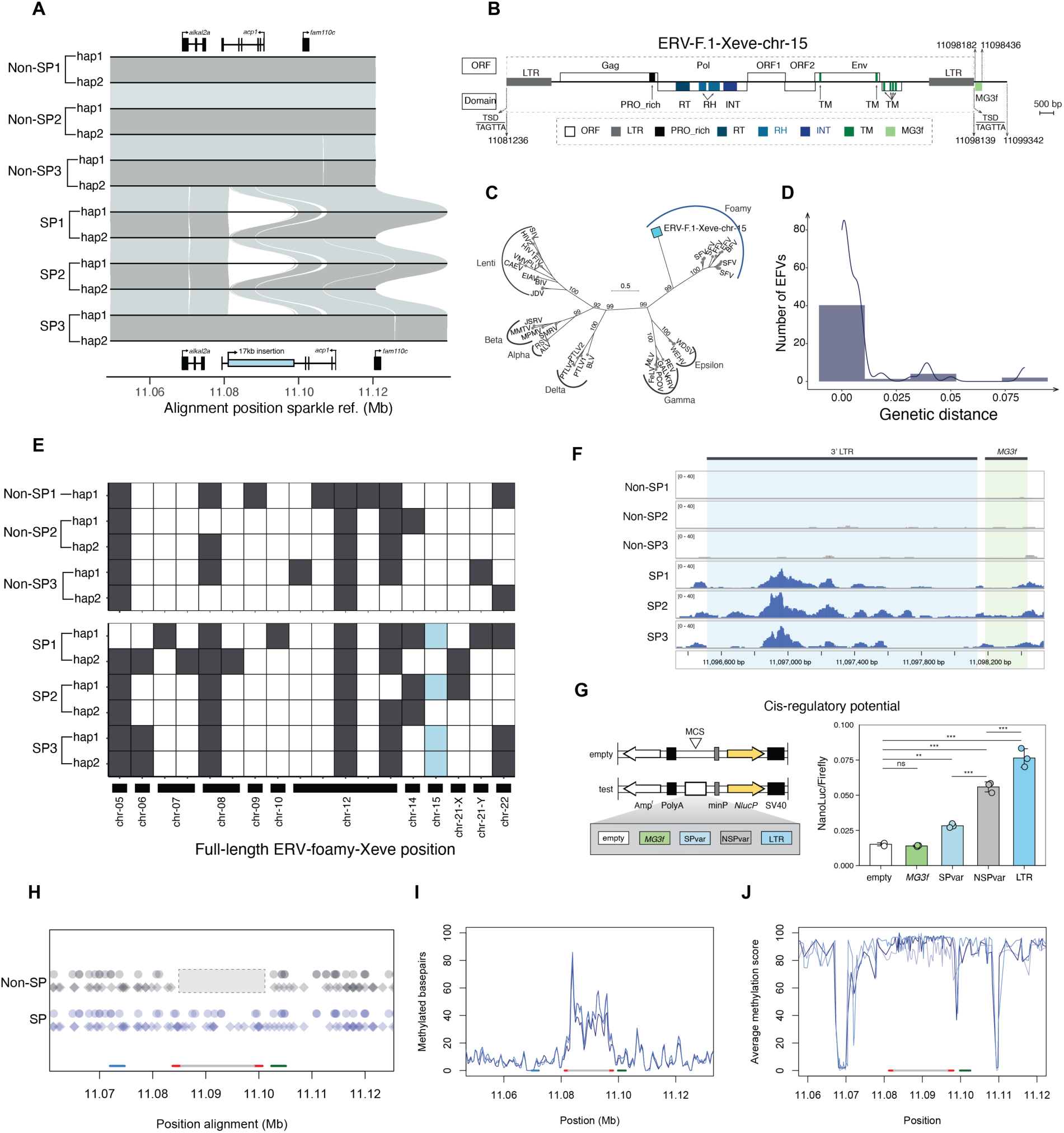
A novel endogenous foamy virus insertion is associated with changes in *alkal2a* gene expression. **A)** Chromosome 15 sequence alignments of long-read assemblies demonstrate a consistent 17 kb insertion near *alkal2a* in at least one haplotype of each sparkle (SP) assembly relative to non-sparkle (non-SP) assemblies. Haplotype 1 – hap1, haplotype 2 – hap2. Gray and blue lines connect colinear alignments across assemblies. Gene models in this region are shown by shaded boxes above and below the alignments, with arrows indicating strand. The 17 kb insertion in the intron of *acp1* is shown in blue below the alignments. **B)** This insertion is an endogenous retrovirus. The ERV-F.1-Xeve-chr-15 insertion exhibits an atypically large genome organization encoding a putative Gag, Pol, Env, and at least three additional open reading frames (ORFs). ERV-F.1-Xeve-chr-15 is flanked by two identical long terminal repeats (LTRs). Target-side duplications (TSDs), TAGTTA, were found on both ends of the insertion. The *MG3f* fragment, which is not typically part of the ERV genome and appears to have originated elsewhere in the *X. evelynae* genome (see Fig. 4), is highlighted in light green. **C)** Phylogenetic analysis of reverse transcriptase (RT) proteins from diverse retroviruses indicates that the endogenous retrovirus insertion on chromosome 15 (blue square) is a foamy retrovirus. **D)** The distribution of genetic distances between the two LTRs flanking each complete ERV-foamy element identified in the 11 *X. evelynae* long-read haplotypes. The line indicates the results of fitting a Gaussian mixture model to this distribution. **E)** Visualization of the highly polymorphic nature of ERV-foamy elements in 11 assembled haplotypes of *X. evelynae* distributed across a total of 18 loci (Supplementary Materials 10). Blue square indicates the ERV-F.1-Xeve-chr-15 insertion associated with the sparkle trait. Top panel: haplotypes assembled from non-sparkle individuals. Bottom panel: Haplotypes assembled from sparkle individuals. Shaded boxes denote the presence of complete ERV-foamy elements located on the chromosomes listed on the x-axis. **F)** ATAC-seq data from non-sparkle (n=3) and sparkle (n=3) samples mapped to the *X. evelynae* sparkle reference genome. Coordinates of the 3’LTR (light blue box) and *MG3f* (light green box) sequences are denoted with dark grey lines at the top of the plot. **G)** Left: schematic of the five plasmids tested in the reporter gene expression assay, including the control empty vector, *MG3f*, sparkle hypervariable region (SPvar), non-sparkle hypervariable region (NSPvar), and LTR sequence flanking ERV-F.1-Xeve-chr-15. Right: The LTR sequence acts as the strongest driver of reporter gene expression in HEK 293-T cells compared to an empty construct and other sequences of interest. One-way ANOVA and Tukey HSD test; ns *P* > 0.05, * *P* < 0.05, ** *P* < 0.01, *** *P* < 0.001. This result was replicated over multiple experiments (Fig. S24). **H)** Results of FIMO analysis scanning for possible transcription factor binding sites in aligned region of chromosome 15 across sparkle and non-sparkle haplotypes. Dashed grey box shows the gap in the alignment between non-sparkle (gray) and sparkle (blue) haplotypes where the ERV insertion occurs. Each diamond shows a predicted *sox10* binding site, and each circle shows a predicted *tfec* binding site. Predicted binding sites detected in sparkle haplotypes are colored in blue and binding sites detected in the non-sparkle haplotypes are colored in gray. The location of *alkal2a* is shown as a blue line on the x-axis, the ERV body is shown as a gray line with flanking LTRs highlighted in red. The green line shows the location of the sparkle and non-sparkle hypervariable region. **I)** Raw number of methylated basepairs (discretized methylation score ≥ 0.5) in all sparkle individuals for which PacBio data was collected. Each individual is shown in a different color shade. **J**) Average discretized methylation score in windows of 10 CpG bases in all sparkle individuals for which PacBio data was collected. Each individual is shown in a different color shade. In both plots, the location of *alkal2a* is shown by a blue line on the x-axis; the location of the ERV-F.1-Xeve-chr-15 insertion is shown by the gray and red lines on the x-axis (red lines indicate LTRs); the location of the sparkle and non-sparkle hypervariable region is shown as a green line on the x-axis.

### The inserted sequence is an endogenous foamy virus

An initial similarity search suggested that this inserted sequence on chromosome 15 was an endogenous retrovirus (ERV). ERV sequences within a host genome typically arise from past retroviral infections which can become vertically inherited retrotransposons by integrating into the germline (*33*). Phylogenetic analysis of reverse transcriptase proteins from representative retroviruses demonstrated that this inserted sequence clearly clusters within a specific genus of endogenous retroviruses, the foamy viruses. We refer to this endogenous foamy virus insertion on chromosome 15 as ERV-F.1-Xeve-chr-15 in our discussions below (Fig. 3B-C; Supplementary Materials 10).

We found evidence that endogenous foamy viruses are widespread in Poeciliidae, the family to which swordtails belong (Fig. S10; Supplementary Materials 10), suggesting that they may have long been associated with the group of species. The ERV-F.1-Xeve-chr-15 insertion encodes the typical Gag, Pol, and Env proteins of endogenous foamy viruses. The entire insertion is flanked by two long terminal repeats (LTRs) at the 5’- and 3’-ends (Fig. 3B). Together with the Gag and Pol proteins, these elements allow the ERV to replicate and insert in the host genome (*33*). Notably, the Env protein is also present and intact, which suggests that this ERV could still be infectious (*55*). Moreover, ERV-F.1-Xeve-chr-15 exhibits an atypically large genome organization, with at least three additional proteins, one of which encodes a transmembrane domain (Fig. 3B).

The 5’- and 3’-LTRs of ERV-F.1-Xeve-chr-15 are identical across all insertion haplotypes, and there are only 2 segregating sites between haplotypes in this entire 17 kb region (Fig. S11). This suggests that ERV-F.1-Xeve-chr-15 inserted recently, a hypothesis we evaluate further below. ERV-F.1-Xeve-chr-15 also contains a small fragment that shares nearly 100% sequence identity to the MG3-super family gene *C3* on chromosome 5 in *Xiphophorus* (hereafter *MG3f*; Fig. 3B, Fig. 4B-C; Supplementary Materials 10 and 11). Despite clearly originating from the *C3* gene on chromosome 5, no ERV elements were detected nearby on chromosome 5 in any sampled individual (closest element 1.8 Mb away; Supplementary Materials 11). Moreover, the *MG3f* element is not associated with other full or partial ERV insertions across the *X. evelynae* genome (Fig. 4C).

**Fig. 4:**
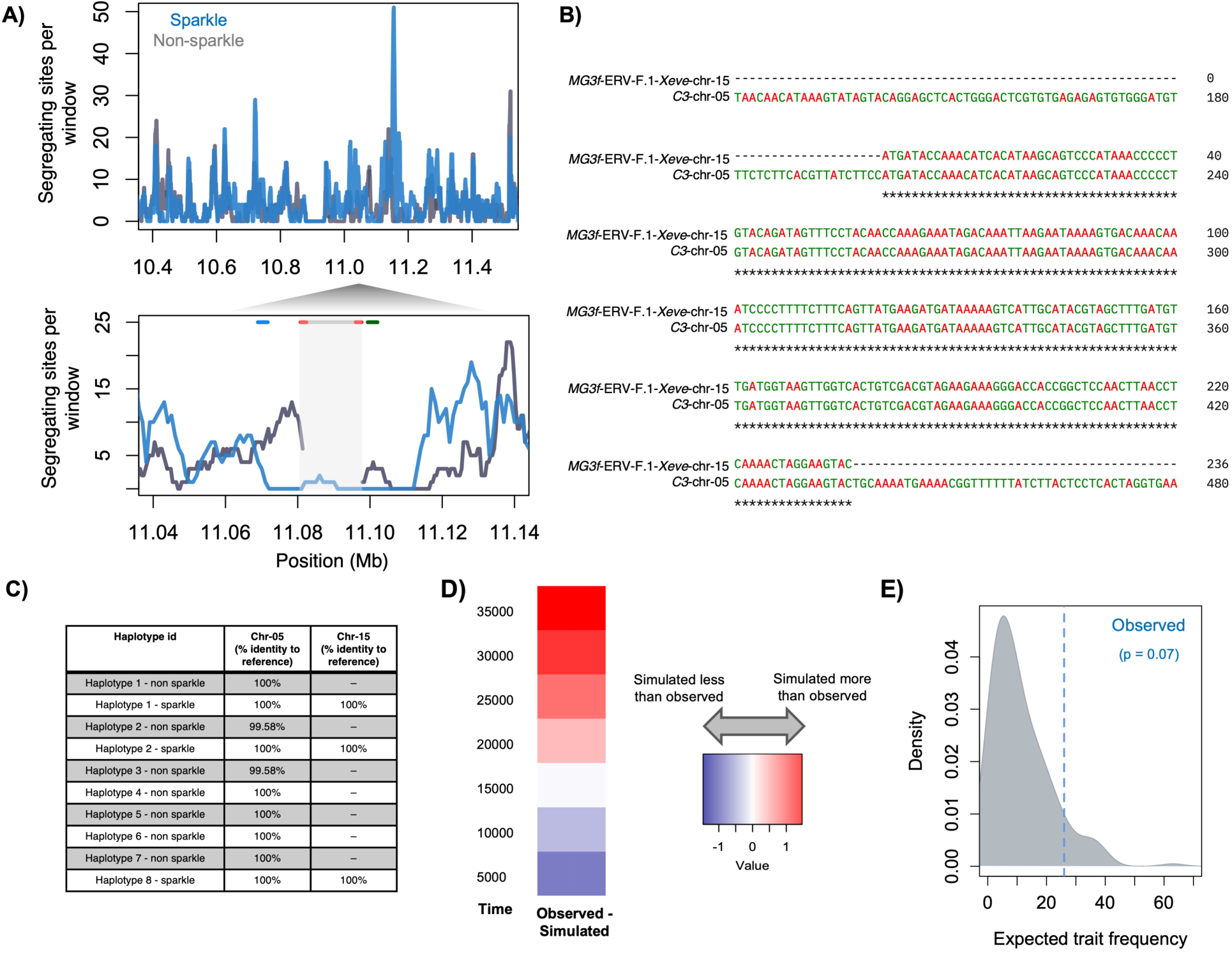
Results of population genetic analyses and simulations focusing on the ERV-F.1-Xeve-chr-15 insertion. **A)** Number of segregating sites across sparkle haplotypes (blue) and a sample-size matched number of wild-type haplotypes (gray) based on a MAFFT alignment of long-read assemblies of this region on chromosome 15. Data are summarized in sliding 5 kb windows. Bottom inset: enlarged plot focusing on the ERV-F.1-Xeve-chr-15 insertion. Note that non-sparkle individuals (gray) have no data in this region since they do not have homologous sequence at ERV-F.1-Xeve-chr-15. Line segments at the top of the plot indicate the locations of *alkal2a* (blue line), ERV-F.1-Xeve-chr-15 (red and gray lines), and the hypervariable region (green line) in this alignment. Note that reduced diversity in the sparkle haplotypes extends beyond the insertion itself. **B)** Alignment of a fragment of the *C3* gene on chromosome 5 to *MG3f* near ERV-F.1-Xeve*-*chr-15. *C3* is a large, 42 exon gene; *MG3f* spans two exons and one intron of the 42 exon *C3* gene. **C)** The *MG3f* sequence uniquely tags the ERV-F.1-Xeve-chr-15 insertion; it is not associated with other ERV elements in the 11 haplotypes we examined. Locations of sequences with high identity to *MG3f* across *X. evelynae* haplotypes compared to the sparkle reference based on blast analysis. All sampled individuals have a complete copy of the *C3* gene on chromosome 5. Only sparkle individuals have this element on chromosome 15, and this element is identical in sequence and length across sequenced sparkle haplotypes. Genome-wide blast searches did not identify any other locations beyond chromosome 5 and chromosome 15 with high sequence identity to *MG3f*. **D)** Results of grid search simulations estimating the likely time since insertion of ERV-F.1-Xeve-chr-15. Simulations modeling 16,000 generations since insertion most closely resembled the sequence variation observed in our dataset. **E)** Given this estimated age since insertion, we also used simulations to ask if the observed frequency of the sparkle trait in the population was unexpected under a scenario of genetic drift alone. Conditional on the establishment of the insertion, we find suggestive evidence that the sparkle trait is at a higher than expected frequency, though this was not significant (p=0.07 by simulation). Gray distribution shows the results of simulations where the insertion established, blue line indicates the observed trait frequency in natural populations.

### Foamy viruses may be actively invading the *X. evelynae* genome

Given evidence that the ERV-F.1-Xeve-chr-15 insertion occurred recently and has the full complement of genes required for transposition, we wanted to evaluate the possibility that foamy viruses are actively replicating in the *X. evelynae* genome. We screened our assembled *X. evelynae* genomes for complete endogenous foamy viruses (the genus of ERVs to which ERV-F.1-Xeve-chr-15 belongs). Within each of our phased assemblies, we identified 4 to 11 complete endogenous foamy viruses across 18 distinct regions of the genome (hereafter ERV-foamy-Xeve sequences; Fig. 3E). Evaluating the distribution of genetic distance between the two LTRs flanking each ERV-foamy-Xeve insertion, we find that it peaks at 0 (Fig. 3D). Since this distribution is correlated with ERV integration time into the host genome, this suggests that the class of foamy viruses to which the 17 kb sparkle-associated insertion belongs is either currently replicating in the *X. evelynae* genome or was recently active (i.e. capable of insertion in new regions of the genome). More directly, we find that the ERV-foamy-Xeve insertions are highly polymorphic across individuals, with only one ERV-foamy-Xeve insertion on chromosome 12 fixed within our sample of 11 haplotypes (Fig. 3E). This type of insertion polymorphism is a hallmark of active transposable elements (*28*).

We next sought to determine whether ERV-foamy-Xeve elements are actively transcribed in *X. evelynae*. We generated bulk RNA-seq data for *X. evelynae* testis and brain tissues and quantified expression of ERV-foamy-Xeve elements from both sparkle and non-sparkle fish (Supplementary Materials 12). We found evidence for low levels of expression of ERV-foamy-Xeve transcripts in both testis and brain (Fig. S12). Note that due to high sequence similarity, we cannot determine which ERV-foamy-Xeve copy is being expressed based on RNA-seq data (Supplementary Materials 12). Together, our results suggest that ERV-foamy-Xeve elements are transcribed in both male germline and somatic tissues in *X. evelynae*, consistent with an ongoing invasion. Despite this evidence of transcription, analysis of 5mC methylation patterns from our long-read sequencing data suggests that the host genome is recognizing and suppressing the ERV-foamy-Xeve elements (*55*, *56*; Fig. S13; Supplementary Materials 13).

Beyond these full-length elements, phylogenomic analyses show that ERV-foamy elements and their solo-LTR derivatives are ubiquitously distributed in the genomes of *Xiphophorus* and its relatives (Supplementary Materials 10; Fig. S10). Moreover, ERV-foamy elements detected within *Xiphophorus* genomes exhibit complex phylogenetic mixing (Fig. S14), which could be attributed to viral host switching, co-diversification between viruses and hosts, or hybridization between host species after virus integration (hybridization is common in *Xiphophorus*; *57*–*59*).

### The endogenous foamy virus insertion has *cis-*regulatory potential

The flanking LTR sequences derived from ERVs are commonly co-opted as enhancers over evolutionary timescales (*61*), but little is known about the impacts of invading or recently active ERVs on gene regulation. Intriguingly, although we find high levels of methylation across ERV-F.1-Xeve-chr-15, we observe a dip in methylation signal in the flanking LTR sequences (Fig. 3I-J). To investigate whether the LTRs of ERV-F.1-Xeve-chr-15 may be acting as a *cis*-regulatory element of *alkal2a*, we evaluated chromatin accessibility along this region of chromosome 15. We collected ATAC-seq data from scale adherent cells from sparkle and non-sparkle individuals (three individuals of each phenotype). We found that the LTRs flanking ERV-F.1-Xeve-chr-15 have increased chromatin accessibility in sparkle individuals, consistent with this region acting as a *cis*-regulatory element (Fig. 3F; Supplementary Materials 14). Notably, other regions in the GWAS peak, including the *alkal2a* promoter sequence, several small structural variants, and a region with a high-density SNPs within the GWAS peak (hereafter the “hypervariable region”), did not differ in chromatin accessibility between sparkle and non-sparkle individuals (Fig. S15-S19; Supplementary Materials 14).

To test the *cis*-regulatory potential of the LTR element inserted near *alkal2a*, we cloned the LTR sequence upstream of a minimal promoter into a luciferase expression vector and compared its effect on reporter gene expression to a vector lacking the LTR, but containing the minimal promoter (empty vector) in HEK 293-T cells (Supplementary Materials 15). We also evaluated other sequences that were associated with the sparkle haplotype but did not show increased chromatin accessibility based on our ATAC-seq dataset, including the *MG3f* fragment and the hypervariable region downstream of the ERV-F.1-Xeve-chr-15 insertion (Fig. 3; Supplementary Materials 15). Results of the luciferase reporter assay demonstrated that the LTR element strongly drives reporter gene activity relative to controls, suggesting that the LTR element inserted near *alkal2a* does indeed have *cis*-regulatory potential (Fig. 3G).

ERVs often harbor a diverse array of transcription factor binding sites that facilitate their activity in the host genome. We were particularly interested in determining whether the flanking LTR regions of ERV-F.1-Xeve-chr-15 contained *sox10* or *tfec* transcription factor binding sites, since these transcription factors play a key role in the differentiation of neural-crest derived progenitor cells into iridophores (*62–65*). We found evidence for multiple *sox10* and *tfec* binding sites within both flanking LTR regions (Fig. 3H). Notably, we compared predicted transcription factor binding motifs between sparkle and non-sparkle haplotypes in the region spanning from 2 kb upstream of *alkal2a* through 6 kb downstream of the ERV-F.1-Xeve-chr-15 insertion. We did not find predicted differences between sparkle and non-sparkle haplotypes in *sox10* or *tfec* binding sites outside of the ERV-F.1-Xeve-chr-15 insertion itself (Supplementary Materials 16; *66*). While *in vivo* editing is not yet possible in *Xiphophorus*, manipulation of these predicted binding sites within ERV-F.1-Xeve-chr-15 will be an exciting future direction once such technologies are established in this system.

### Population genetics of the endogenous foamy virus insertion on chromosome 15

Our data indicate that an active or recently active endogenous foamy virus controls variation in a structural color phenotype in *X. evelynae*. We were interested in whether the patterns of genetic variation in our data were indicative of directional or balancing selection on sparkle haplotypes containing the ERV-F.1-Xeve-chr-15 insertion. While genetic diversity was ∼10X lower in the insertion compared to other regions of the genome (Fig. 4A), this pattern alone is not diagnostic, since it could be consistent with either a recent insertion of the ERV-F.1-Xeve-chr-15 element, selection on this haplotype within this *X. evelynae* population, or both.

We implemented population genetic tests to investigate possible evidence for selection on this region (Supplementary Materials 17). Since ERV-F.1-Xeve-chr-15 is not present in all individuals, we focused our analysis on regions flanking the insertion in our 11 assembled chromosome 15 haplotypes. We observed reduced genetic diversity and high between-haplotype divergence in the flanking sequence of sparkle haplotypes compared to non-sparkle haplotypes (Fig. 4A; Fig. S20), differences in Tajima’s D (Fig. S20), and modest signatures of extended haplotype homozygosity in SNPs linked to the sparkle haplotype (Fig. S20; Supplementary Materials 17). Tajima’s D exceeded 2 in flanking windows on both sides of the insertion, potentially consistent with balancing selection, but we note that these values were only modest outliers compared to other aligned regions on chromosome 15 (95% quantile; mean = 0.22; Fig. S20).

We also used population genetic approaches to estimate the age of the ERV-F.1-Xeve-chr-15 insertion event. We modeled the demographic history of *X. evelynae* (Fig. S21) and used a grid search approach implemented in SLiM (*67*) to infer the likely insertion time of ERV-F.1-Xeve-chr-15 (Supplementary Materials 18). Simulations modeling an insertion time of 16,000 generations before the present minimized the difference between the observed and simulated data (Fig. 4D). Given this estimated insertion time, the sparkle phenotype is at higher-than-expected frequency in the population, though not significantly so (Fig. 4E; p=0.07). In variations of these demographic analyses, the maximum insertion time consistent with our data was 20,000 generations (Supplementary Materials 18). Together with other results, these findings suggest that this ERV is active in the *X. evelynae* genome, providing a rare opportunity to study an invading endogenous retrovirus.

### Sparkle generates unique visual signals that may impact predator responses

Reflective traits like the sparkle phenotype often impact signaling to conspecifics and heterospecifics (*68*). We measured the reflectance spectra of male and female *X. evelynae* and found that sparkle individuals maximally reflect light at a 90° incident angle, with no differences in UV reflectance across individuals (Fig. 5B-C; Fig. S22; Supplementary Materials 19). We analyzed whether *X. evelynae* individuals with and without the sparkle trait differ in boldness or in attractiveness to mates but did not find significant effects in these behavioral assays (Fig. 5D-E; Supplementary Materials 20).

**Fig. 5:**
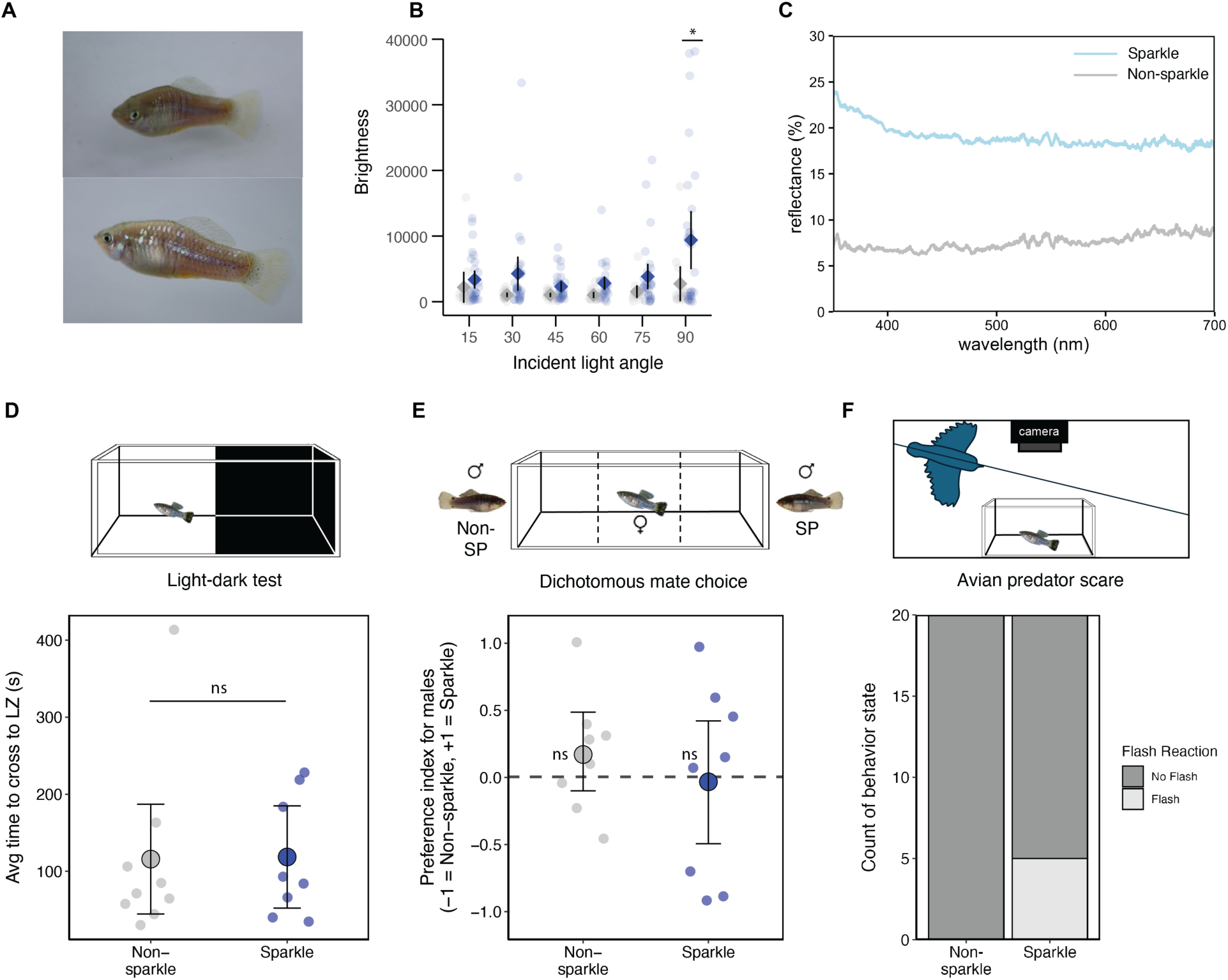
Fish with sparkle produce distinct visual signals, which may impact their appearance in the natural environment. **A)** Two photographs of the same *X. evelynae* fish at distinct angles relative to a light source, demonstrating how fish with the sparkle trait can change in appearance as they move. **B)** Brightness measured at 15°, 30°, 45°, 60°, 75°, and 90° incident light angles in non-sparkle (grey) and sparkle (blue) individuals. Data were analyzed using a linear model. The brightness of the sparkle trait significantly differed from that of the non-sparkle trait on brightness, but only at a 90° angle (p-value = 0.028). **C)** Light reflectance measurements at a 90° angle in sparkle and non-sparkle fish using a spectrophotometer. Percent reflectance at a 90° angle was greater in sparkle fish at all wavelengths measured. **D)** Results of scototaxis assay that tests boldness of individual fish by measuring average time to transition (seconds) from dark to light zone of tank. No significant differences were found between phenotypes (t-test; p-value = 0.95). LZ = light zone. **E)** Results of female dichotomous mate-preference assay measuring time spent associating with non-sparkle or sparkle male animations presented on either side of the female trial tank. Fish did not significantly differ from the null hypothesis of no preference for either phenotype (two-sided Wilcoxon signed rank test; non-SP p-value = 0.38; SP p-value = 1). **F)** Avian predator response assay involves simulating a bird predator using a realistic Kingfisher model and recording fish response using an overhead GoPro camera. Fisher’s exact test for count data shows significant differences between signals produced by non-sparkle and sparkle fish in this assay (p-value = 0.047).

Underwater videos taken of natural swordtail populations provided a different clue for the potential function of this trait. Individuals with the sparkle phenotype appeared to “flash” and as they changed directions in the water (Video S2). Past studies have suggested that such signals can make prey harder to localize (*68–72*). Consistent with our observations in the field, previous work indicates that birds are common predators of poeciliid fish like *Xiphophorus* (*73–75*). We implemented trials in the lab where a model bird appears to swoop over the tank (Video S3; Supplementary Materials 20). In response to this perceived predator, both sparkle and non-sparkle individuals display a stereotyped escape response. However, the signal produced by sparkle individuals during this escape response differed from that of non-sparkle individuals (Fig. 5F). Sparkle fish produced a flash that was visible from an overhead camera in 25% of the escape response trials (Fig. 5F). This suggests that increased reflectivity could impact the signal produced during pre-existing predator avoidance behaviors. Thus, this novel iridescence trait may layer onto existing behavioral strategies to startle or confuse predators.

### Implications

We describe the genetic, cellular, and developmental mechanisms underlying the recent evolution of a novel iridescence trait in swordtail fish. This trait is visually striking and detectable in the wild, hinting that sparkle may impact survival of these fish in their natural populations. Unlike some iridescence traits that have been studied in other fish, the sparkle trait does not appear to impact female mate preference (Fig. 5; *72*–*74*). We hypothesize that the sparkle trait may startle potential predators or interfere with their ability to localize fish during stereotyped escape responses (*79*). Previous studies have shown that iridescent prey are more difficult for avian predators to localize (*68–72*). In this regard, we note that phenotypically similar iridescence patterns have been identified across at least three distinct swordtail species in this region of Mexico (Supplementary Materials 1), raising the exciting possibility that the trait has evolved recurrently due to shared ecological pressures.

Our results indicate that the evolution of the sparkle trait was driven by the recent insertion of an endogenous retrovirus near the gene *alkal2a*, where it appears to act *cis*-regulatory element. The recent origin of the insertion, presence of a complete ERV genome, and evidence of many low frequency insertion polymorphisms, all indicate that this ERV foamy element may be actively invading the genome. Indeed, the full-length ERV-foamy-Xeve elements we detect not only maintain the Gag and Pol proteins required for transposition but also the Env protein, hinting that these retroviruses could still be infectious. While LTR sequences from ERVs are frequently co-opted by the host genome in gene regulation over evolutionary timescales, this is among the first cases where an invading retrovirus regulates a functionally important trait in natural populations. The best studied case of active ERVs comes from natural populations of Koalas. In this species, active ERVs that began invading the genome ∼10,000 generations ago are associated with neoplasia (with some populations showing evidence of evolved resistance; reviewed in *73*). In other cases, active ERVs have not been linked to phenotypic effects (*81*, *82*) or are associated with inherited abnormalities (e.g. *83*, *84*).

There are several possible fates for integrated ERVs over evolutionary timescales. The majority of these ERVs will be degraded through the accumulation of disruptive mutations, rendering them transpositionally inactive. Despite this inactivation, it is common for these sequences to be used by the host to serve as *cis*-regulatory sequences (e.g. *22*) or produce retroviral gene products that are co-opted by the host cell (e.g. *32*). Intriguingly, these elements often become indispensable to the host such that they are required for normal development (reviewed in *74*). By contrast, the sparkle insertion is not essential for development and likely represents one of the earliest stages of ERV co-option and coevolution with the host. Tracking the dynamics of evolutionary changes in such active elements will yield novel insights into the mechanistic causes and evolutionary consequences of ERV co-option.

## Acknowledgements

We are grateful to the Federal Government of Mexico and to Benemérita Universidad Autónoma de Puebla for permission to collect fish. We thank members of the Schumer lab, Jessica Feldman, Dominique Bergmann, Adam Reeves, Landen Gozashti, Peter Sudmant, James Gagnon, Claire Tobin, Raquel Fueyo, Emilia Santos, and Santos lab members for helpful comments on experiments and/or earlier versions of this manuscript. Stanford University and the Stanford Research Computing Center provided computational support for this project. We thank the NMS core at Stanford for assistance with DIC imaging and the laboratories of Eric Peterman, Emília Santos, Nick Altemose, and David Parichy for helpful input. We thank Claudia Gehrig-Höhn for excellent technical support at the Imaging Core Facility of the Biocenter. A subset of data visualization scripts were developed with assistance from OpenAI’s ChatGPT (GPT-4 and -5 models). We thank the CICHAZ field station for their support of this project. **Funding:** TOD and NBH were supported by NSF GRFP awards DGE-2146755. NBH and WP were supported by the Cell and Molecular Biology training grant T32-GM007276. BMM was supported by NSF GRFP award DGE-1656518. This research was funded by the Searle Scholars Program, Pew Biomedical Scholars Program, and Freeman Hrabowski Scholars award to MS. The JEOL JEM1400Flash transmission electron microscope is funded by the Deutsche Forschungsgemeinschaft (DFG, German Research Foundation) – 426173797 (INST 93/1003-1 FUGG). CAG is the Akiko Yamazaki and Jerry Yang Faculty Scholar in Pediatric Translational Medicine, supported by the Stanford Maternal Child Health Research Institute. ZO was supported by the Wu Tsai Neuroscience Interdisciplinary Scholar program.

## Author contributions

N.B.H., M.Schu. designed the project; N.B.H., T.O.D., J.J.B., T.R.G., W.P., R.S., P.F.Z., A.K., S.D., N.S.O., T.T.Y., Z.O., C.Y.P., S.K.H., D.L.P., M.S. collected data; N.B.H., T.O.D., Q.H., K.D., P.F.Z., G.A.P., B.M., performed analyses; G.Z.H., D.L.P., M.S., C.G., Z.O., K.E.H. provided expertise and technical support.

## Data and materials availability

Raw data are available through NCBI Sequence Read Archive and processed data are available through Dryad (Accessions Pending). Code is available on github (https://github.com/Schumerlab).

