## Supplementary Materials for "Insertion of an invading retrovirus regulates a novel color trait in swordtail fish"

**Affiliations:**
[1]Department of Biology, Stanford University
[2]Centro de Investigaciones Científicas de las Huastecas "Aguazarca" A.C.
[3]College of Life Sciences, Nanjing Normal University, Nanjing, Jiangsu 210023, China
[4]Xiphophorus Genetic Stock Center, Texas State University San Marcos
[5]State Key Laboratory of Mariculture Biobreeding and Sustainable Goods, Yellow Sea Fisheries Research Institute, Chinese Academy of Fishery Sciences, Qingdao, Shandong
[6]Laboratorio de Ecología de la Conducta, Instituto de Fisiología, Benemérita Universidad Autónoma de Puebla
[7]Department of Materials Science & Engineering, Stanford University
[8]Current address: National Oceanic & Atmospheric Administration and UC Santa Cruz
[9]Imaging Core Facility, Biocenter of the University of Würzburg
[10]Department of Biology, Louisiana State University
[11]Institute of Pathology, University of Würzburg, Germany
[12]Research Department for Limnology, University of Innsbruck, Mondsee, Austria
[13]Institute for Stem Cell Biology and Regenerative Medicine, Stanford University
[14]Department of Pediatrics, Stanford University
[15]Department of Genetics, Stanford University
[16]Howard Hughes Medical Institute

### Supplementary Materials 1. Sample collection and phylogenetic analysis

*Sample collection*

Samples for the genome-wide association study were collected using baited minnow traps from the Juntas Chicas population in Veracruz, Mexico (20°34'12.49"N 98°1'58.83"W). Each sexually mature individual was anesthetized in MS-222 diluted to 100 mg/mL in river water and photographed using a copy stand set up with a Nikon d90 DSLR digital camera. We included a standardized grid background, ruler, and a ColorChecker Passport Photo 2 as a color standard in each image. We next took a small fin clip from the caudal fin of each individual and returned them to their sampling site after allowing them to recover in a holding tank (following Stanford APLAC 33071). A subset of individuals was transported back to the CICHAZ field station and later to Stanford to establish a lab breeding colony.

*Phylogenetic analysis to determine species of origin for the Juntas Chicas population*

*Xiphophorus* has three major phylogenetic groups: Platyfish, Northern Swordtails, and Southern Swordtails (*1*). The entire group is often referred to as "swordtails" or "swordtails and platyfish". Samples formerly collected from the geographical region where the Juntas Chicas population occurs have previously been reported to belong to the species *X. variatus* (*2*), but the phylogenetic placement of this geographically distinct population has not been evaluated with genomic data. To identify the most closely related lineage to the Juntas Chicas fish, we performed phylogenetic analyses including the sample used to generate our reference genome (see below) and whole genome sequences from all available *Xiphophorus* species.

Briefly, we inferred the phylogenetic tree based on whole genome sequence alignment (WGA). The approach used was based on a previous study (*1*). First, the genome of each species was aligned to that of *X. maculatus* using minimap2 (*3*). Second, the pairwise alignments were then refined using Genome alignment Tools from the Hiller lab (*4*). Third, we merged these pairwise genome alignments into the multiple WGA using MULTIZ (*5*). Finally, based on the WGA, a maximum likelihood phylogeny was inferred using RAxML under the GTR+Gamma model with 1,000 rapid bootstraps (*6*). Before implementing phylogenetic inference with RAxML, alignment blocks shorter than one kilobase were removed. The left alignment blocks were also trimmed using trimAl (*7*).

The resulting phylogeny had 100% bootstrap support at every node and placed the Juntas Chicas population as most closely related to *X. evelynae* among described *Xiphophorus* species (Fig. S1). We note however, that the sequenced lineage is somewhat divergent from extant *X. evelynae* lineages and may ultimately be considered for designation as its own species by experts in this area.

*Note on previous studies of a similar trait iridescence trait and nomenclature*

When we identified the sparkle trait in the Juntas Chicas population, we noted that it appeared phenotypically similar to a previously described iridescence trait referred to as "argent" in the popular aquarium literature (*8*). To our knowledge, "argent" has been reported a single time in the scientific literature, in a species identified as *X. variatus* (*2*). Species identification in Chen and Borowsky (2004) relied on phylogenetic inference based on RAPD data. Moreover, the *X. variatus* populations sampled in this earlier study are found to the south of the Juntas Chicas *X. evelynae* population identified in our study (>25 km).

In describing the iridescent phenotype that we observed in the Juntas Chicas *X. evelynae* population, we considered whether it should be treated as synonymous with the “argent” phenotype identified in the aquarium literature and discussed in Chen and Borowsky (2004). However, we opted to treat this as a potentially distinct phenotype for the purposes of this manuscript for several reasons. First, it was unclear to us whether the phenotype Chen and Borowsky (2004) discuss is limited to *X. variatus*. Second, we were unsure about the accuracy of species identification in this previous study. The molecular data collected by Chen and Borowsky did not readily distinguish between *X. evelynae* and *X. variatus*, raising the possibility of species misidentification or that the trait is shared across species. Since Chen and Borowsky (2004) refer to the species they collected with argent as *X. variatus*, we continue with this assumption here, while noting that species identification may shift as genomic data becomes available for these populations.

A final reason we treat the “argent” trait discussed in Chen and Borowsky (2004) as distinct from “sparkle” is due to a lack of clarity on the characteristics of the phenotype in the available literature. In photographs of the fish collected by Borowsky and in the aquarist literature, we see evidence of two distinct patterns. One pattern occurs along the horizontal midline of the fish (as we see in Juntas Chicas fish; Fig. 1), while the other appears to be a distinct speckled phenotype with iridescent scales distributed across the body. Strikingly, a third species found in this region, *X. maculatus*, also has a visually similar iridescence pattern. However, our examinations of this population of *X. maculatus* in lab have indicated that iridophores are not concentrated in the scales as they are in the *X. evelynae* Juntas Chicas population and are instead found in deeper skin layers. Thus, we conclude that iridescence is a common trait in this region of Mexico, but iridescence traits are seemingly found in varying forms across multiple species (see Discussion). While uncertainties in species delimitation and evidence of convergent evolution of similar traits among *Xiphophorus* in this region of Mexico result in us treating sparkle as a distinct trait from the previously described argent phenotype (*2*), future studies may warrant revisiting this naming convention.

**Supplementary Materials 2. Imaging of sparkle scales with staining, DIC, and TEM**

*Differential interference contrast (DIC) microscopy of sparkle and non-sparkle scales*

Non-sparkle (n=2) and sparkle (n=2) scales were removed from the left and right side of 3 sparkle and 3 non-sparkle *X. evelynae* individuals. Scales were sampled from the midline region. To mount scales while preserving the 3-dimensional structure of the sample, we placed 3 binder reinforcing stickers on a microscope slide, deposited the isolated scale inside the well created by the stickers, surrounded the scale with fish water, and gently added on a cover slip. Each scale was imaged with differential interference contrast microscopy (DIC) on a Zeiss AxioImager Motorized Widefield Fluorescence Microscope at a constant exposure (10ms), allowing for the optimal visualization of light-reflecting sub-cellular structures that illuminated the iridophore cells attached to each sparkle scale.

*Transmission electron microscopy of fish scales*

In preparation for transmission electron microscopy (TEM), fish were euthanized and samples were washed in PBS and fixed in 2.5% glutaraldehyde overnight at 4°C. Further processing is described in (*9*) for EM preparation of cells. Embedded samples were cut in ultrathin slices of about 80 nm thickness and contrasted for 15 minutes with 2.5% uranyl acetate in dissolved ethanol and 8 minutes in lead citrate after Reynolds (*10*) diluted 50% in water. Electron microscopy analysis was performed at a JEOL JEM1400Flash transmission electron microscope at 120kV with a Matataki Flash camera.

**Supplementary Materials 3. Developmental timing of sparkle in *X. evelynae***

To characterize the sparkle trait over development, we individually tracked, imaged, and phenotyped 30 *X. evelynae* fry born in the lab from 1 week of age until they reached sexual maturity. These fish were derived from 5 broods and were imaged weekly for at least 17 weeks. Fish were individually housed in zebrafish tanks to track fish identity, since they were too small to label with elastomer tags at the beginning of the experiment. Every week, fish were anesthetized in MS-222 mixed with fish water, inspected under a dissecting microscope for iridophore emergence to the scale tissue, and imaged with a color standard on a Canon EOS 77D camera with flash using a Meike MK-14EXT TTL Macro Ring Flash. Confirmation of sparkle trait development under a dissecting microscope was necessary to accurately determine the age of sparkle onset because iridophores are not visible by eye until later in sparkle development when fish are larger. Out of the 30 tracked fish, 7 individuals were non-sparkle and 23 individuals ultimately developed the sparkle trait, with the onset age ranging from 10-52 days after birth (swordtails and their relatives are livebearing species). Note that the typical frequency of the trait in the wild is only ~25% (see main text), but crosses in the lab were designed to enrich for the sparkle phenotype.

**Supplementary Materials 4. Image analysis highlights consistent spatial patterns in the sparkle phenotype**

We found that *X. evelynae* adults collected from the wild and born in lab developed the sparkle trait primarily on the midline (Fig. S3). To formally quantify the spatial patterns of this trait, we took 30 images of adult fish with sparkle from our GWAS population (9 male, 21 female), white-balanced each photo using ColorChecker Passport Photo 2, and assigned landmarks along the fish body for each image (see Fig. S23). We did not observe sexual dimorphism in the presence or absence of the sparkle trait or in the trait patterning. Some differences were observed in reflectance in our spectral analysis, but we speculate that this may be due to other male coloration traits that develop during maturation (Supplementary Materials 19). We used these landmarks to align all fish to a reference body using the alignLan function in the StevenVB12/patternize package in RStudio (https://github.com/StevenVB12/patternize). We then converted raster objects to image arrays and plotted these as PNGs. We manually colored the sparkle trait in Adobe Photoshop using the pen tool. To calculate and visualize the incidence of sparkle across aligned adult fish flanks (i.e. how often a pixel is part of a sparkle mask across all 37 adult fish images), we summed all the sparkle mask binary matrices, where a pixel with sparkle is coded as 1 and without sparkle is coded as 0, and normalized each pixel count by the number of total masks to generate a matrix representing the proportion of images in which each pixel had iridescence. These data were plotted as a heatmap using geom_raster to identify consistent sparkle patterns in adult fish.

**Supplementary Materials 5. Long read assemblies with PacBio HiFi**

*Genome assembly*

Six chromosome-level *X. evelynae* genomes were generated using PacBio HiFi long-read sequencing. DNA was extracted using the NEB Monarch HMW extraction kit according to manufacturer's directions. All steps used wide-bore pipette tips to avoid shearing the DNA. Briefly, fish were euthanized via ice (for the *X. evelynae* sparkle and non-sparkle reference genomes) and via tricaine (for the 4 additional *X. evelynae* genomes sequenced). Brain tissue was dissected and frozen at -80°C for less than 1 week. We weighed tissue mass to determine whether to follow the standard input protocol (10-25 mg) or a low input protocol (5-15 mg). Tissue was mechanically homogenized and transferred to lysis buffer with proteinase K. Samples were re-homogenized with provided pestles, lysis master mix was added, and samples were incubated at 56°C for 1 hour with agitation. RNase A was added, samples were mixed by inversion 7 times, incubated for 10 minutes at 56°C, and lysate was chilled on ice for 3 minutes before adding Protein Separation Solution to ensure efficient phase separation. Finally, samples were centrifuged for 30 minutes at 16,000xg and transferred to a provided 2 mL tube. DNA capture beads were added to each tube along with isopropanol, and tubes were mixed for 5 minutes by hand to allow genomic DNA to attach to beads. Liquid was discarded and genomic DNA was washed twice with Wash Buffer. Beads were placed into a new tube and genomic DNA was eluted. Extracted DNA for the *X. evelynae* sparkle reference fish was sent to University of Washington PacBio sequencing core (Seattle, Washington) to be sequenced on one Revio SMRT cell, and DNA from the *X. evelynae* non-sparkle reference fish was sent to Cantanta Bio, LCC (Scotts Valley, California) to be sequenced on two Sequel SMRT cells. DNA for the four additional *X. evelynae* genomes was extracted as described above and sent for sequencing at the University of Washington PacBio sequencing core on one Revio SMRT cell.

Adapters were filtered with HiFiAdapterFilt.sh and NanoPlot was used to assess read quality and ensure CCS Q>20 (*11*). To assemble reads into a phased diploid genome, hifiasm was run with default parameters and Bandage was used to confirm untig.gfa plots were well resolved, especially near the *alkal2* gene and ERV-F.1-Xeve-chr-15. For the reference *X. evelynae* genomes, RagTag (https://github.com/malonge/RagTag; (*12*) was used to scaffold each haplotype to *X. variatus*.

*Xiphophorus* species have young, weakly differentiated sex chromosomes (*13*). In most species studied to date, the sex chromosome appears to be chromosome 21 (*13*). To identify haplotigs that likely correspond to the sex chromosomes, we aligned the scaffolded assemblies to PacBio HiFi genome assemblies for *X. malinche* (*1*) and *X. nezahualcoyotl* (Given et al., *in prep*) with minimap2 (*3*). These assemblies contain the X but not the Y chromosome. For the non-sparkle *X. evelynae* reference, there were exactly two haplotigs that aligned to chromosome 21. These alternative haplotigs aligned across the length of chromosome 21 and we considered these likely be the X and Y chromosomes (*13*). The chromosome with more inversions and repetitive content relative to the reference X-chromosomes was treated as the putative Y-chromosome for the purposes of our analyses. For the sparkle reference, there was one haplotig that aligned and was syntenic across the entire length of the known X-chromosomes from *X. malinche* and *X. nezahualcoyotl*. Therefore, we considered this haplotig to be the likely X-chromosome. In addition, there were two other haplotigs in the sparkle reference assembly that aligned to chromosome 21 in other species (*X. malinche* and *X. nezahualcoyotl*), which we joined with ragtag.py agp2fa. We aligned these joined haplotigs to the putative X-chromosomes of the sparkle and non-sparkle references and the known X chromosomes of *X. malinche* and *X. nezahualcoyotl.* We found that these joined haplotigs in the sparkle assembly had an inversion relative to each of these known or putative X chromosome assemblies. We thus treated this joined haplotig as the putative Y-chromosome in the sparkle reference. To represent both the putative X and Y chromosomes in the final *X. evelynae* assemblies, we aligned the putative X and Y haplotypes and used the inversion breakpoints to mask the Y-chromosome sequence that is homologous with the X (following *14*).

The same procedure was used to identify the alternate haplotypes for chromosome 15 for the sparkle *X. evelynae* reference. However, instead of masking homology between the two haplotypes like we did for chromosome 21, we instead generated two versions of the primary assembly, one with each of the two haplotigs from chromosome 15.

We also repeated this procedure to generate assemblies and haplotigs for an additional two sparkle and two non-sparkle individuals. This allowed us to formally compare genome structure on chromosome 15 across haplotypes belonging to sparkle and non-sparkle individuals (see below). The average coverage for these additional individuals was 57X and ranged from 27-131X. For assembly statistics for these additional individuals see Table S4.

*Genome annotation*

We annotated the genome assemblies developed for this research using a custom pipeline adapted from previous approaches used by our groups (*1*, *15*). This workflow integrates homology-based, transcriptome-based, and *ab initio* gene prediction approaches to generate a comprehensive set of gene annotations. Briefly, protein sequences and RNA-seq reads were aligned to the assemblies to provide homologous and transcript-derived evidence, respectively. In parallel, *ab initio* gene predictions were carried out using AUGUSTUS (*16*). All evidence was subsequently integrated into a unified gene set.

For homology-based annotation, we curated a dataset comprising 505,310 protein sequences, sourced from the vertebrate entries in Swiss-Prot (https://www.uniprot.org/statistics/Swiss-Prot), the RefSeq "vertebrate_other" subset (filtered for entries with IDs starting with "NP"), and genome annotations from NCBI for human (GCF_000001405.39_GRCh38), zebrafish (GCF_000002035.6), platyfish (GCF_002775205.1), medaka (GCF_002234675.1), mummichog (GCF_011125445.2), turquoise killifish (GCF_001465895.1), guppy (GCF_000633615.1), and shortfin molly (GCF_001443325.1). These proteins were aligned to the assembled genome using GeneWise and Exonerate (*17*). To improve computational efficiency, GenblastA was first employed to identify candidate regions for GeneWise alignment (*18*). In cases where multiple gene models overlapped within the same genomic region, only the highest-scoring model was retained.

To incorporate transcriptome evidence, RNA-seq datasets derived from *X. maculatus*, a closely related species to *X. evelynae* (Fig. S1), were aligned to the genome using HISAT (*19*). Gene models were reconstructed and refined from these alignments using StringTie (*20*). In parallel, Trinity was used for *de novo* transcriptome assembly, and the resulting transcripts were aligned to the genome with Splign (*21*) to infer exon-intron structures.

*Ab initio* gene prediction was conducted using AUGUSTUS (*16*), which was trained in two stages. The initial training utilized BUSCO (*22*) with the "--long" parameter to generate an initial model set. A second round of training was performed using high-confidence gene models that were consistently predicted across multiple tools (GeneWise, Exonerate, StringTie, and

Splign). We then executed AUGUSTUS using both the homology and transcriptome evidence for external hints.

To finalize the annotation given these multiple methods, each locus was systematically evaluated by comparing *ab initio* and homology-derived predictions. For loci with multiple competing gene models, preference was given to the gene model best supported by transcriptomic data. When the selected gene model contained poorly supported exons (i.e. with low sequence identity or lacking transcriptome evidence), alternative exon models were re-evaluated for better-supported exon structures. Finally, gene predictions that lacked both transcriptome support and homology evidence were excluded from the final annotation.

*Comparison of synteny to closely related species*

To evaluate how our newly generated genome assemblies compared to two existing high quality reference genomes of *Xiphophorus* species, we compared the *X. evelynae* reference to both the *X. maculatus* (*13*); NCBI GCA_002775205.2) and *X. birchmanni* (*14*) reference assemblies. We performed whole chromosome alignments using MUMmer4 (*23*) and plotted the alignment results to identify any large structural variants that differed between *X. evelynae* and related species. Overall, we found high synteny between the *X. evelynae* reference genome and genomes from these species. Notably, karyotype was highly conserved between *X. birchmanni* and *X. evelynae*, and where inversions did occur, many were restricted to the sub-telomeric regions. Since *X. birchmanni* and *X. evelynae* belong to distinct evolutionary clades within *Xiphophorus* (Northern Swordtails and Platyfish, respectively), this is suggestive of high levels of conservation in chromosome structure. Example alignments with both *X. maculatus* and *X. birchmanni* for chromosome 1 are shown in Fig. S25. A whole genome alignment is shown in Fig. S26.

*Analysis of structural variation within the sparkle GWAS peak*

Initially, we visualized PacBio reads mapped to the *X. evelynae* reference genome in IGV and discovered evidence of a ~17 kb insertion near the *alkal2a* gene (see below). We performed pairwise alignments between the *X. evelynae* sparkle and non-sparkle haplotypes using MUMmer4 (*23*) and visualized non-syntenic regions within the GWAS peak region (Fig. 3).

We also performed a formal analysis to identify variants and rearrangements in this region that differed between sparkle and non-sparkle long-read assemblies using the phased assembly caller PAV (https://github.com/EichlerLab/pav). Structural variant callers for long read data are still developing, but PAV has good performance detecting insertions, deletions, inversions, duplications, as well as single nucleotide variants (*24*). Within the significantly associated region identified from our genome-wide association study, we found 96 total segregating structural rearrangements greater than 50 bp, 24 of which were within 300 kb of the gene *alkal2a* (Fig. S27). The vast majority of these variants are rare and do not segregate with pigmentation phenotype (Fig. S28). However, four structural variants and a total of 42 single-basepair variants segregate with the sparkle phenotype in this interval. Focusing on the highly significant interval in our GWAS from 10.75-11.16 Mb (10.68-11.15 in the non-sparkle reference assembly), there are 35 single-basepair variants, 8 indels, and four larger structural variants (>50 bp) that segregate with the phenotype. This included a large, ~17 kb insertion in the intron of the gene *acp1* that was present in all sparkle individuals and absent from all non-sparkle individuals (Fig. S27; Fig. S28). We note that the *acp1* gene was not differentially expressed in our scale

RNA-seq dataset (see Fig. 2C). We focused on these regions for downstream analysis of chromatin accessibility and in reporter assays.

*Analysis of protein evolution*

We performed several analyses of predicted protein sequences. First, we evaluated whether nonsynonymous changes existed that differentiate *X. evelynae* individuals with and without the sparkle trait. We used the annotated *alkal2a* cDNA sequence from the Juntas Chicas sparkle reference assembly as a guide sequence for AUGUSTUS *de novo* annotation of *alkal2a* on chromosome 15 in our long-read assembly pseudohaplotypes. We found no synonymous or nonsynonymous substitutions segregating within the eleven *alkal2a* sequences from sparkle and non-sparkle individuals from Juntas Chicas.

In addition, we extracted *alkal2a* sequences from available annotated *Xiphophorus* genome assemblies and used the program codeml to infer the rate of nonsynonymous substitutions per nonsynonymous sites versus synonymous substitutions per synonymous sites (dN/dS). This resulted in alignment of *alkal2a* sequences from 10 species (Fig. S29). The estimated dN/dS based on this alignment was 0.055. We used a likelihood ratio test to compare the observed data to a model of neutral evolution by setting omega to 1. Based on this test we found that we could reject a model of neutral evolution ($\chi$=11.4, $p<0.001$), suggesting that *alkal2a* is under purifying selection in *Xiphophorus*.

**Supplementary Materials 6. Genetic basis of the sparkle trait in the Juntas Chicas population**

*GWAS analysis*

The sparkle trait is polymorphic in the *X. evelynae* population in Juntas Chicas, Veracruz, Mexico, allowing us to perform a genome-wide association study (GWAS). We photographed and fin-clipped 262 individuals from this population (66 cases with sparkle and 196 controls without sparkle) in February 2023. DNA was extracted from fin clips using the Agencourt DNAdvance kit (Beckman Coulter A48705) according to the manufacturer's protocol, but half reactions were used. Briefly, tissue was incubated in lysis LBH buffer, DTT, and proteinase K overnight. Pre-Bind PBBA buffer was added the following day and Bind BBE beads were mixed and allowed time to separate onto a magnet. Supernatant was aspirated, beads were washed with ethanol twice, and DNA was eluted. DNA was quantified and diluted in preparation for tagmentation with a Tn5 enzyme (Illumina #20034197). A unique i5 and i7 primer were added for PCR, and a final bead-based library cleanup was performed. Low coverage (~1x) whole genome sequencing was performed on the Illumina HiSeq at Admera to generate paired-end 150 basepair reads on a pooled library.

Because *Xiphophorus* are known to frequently hybridize and we also found *X. malinche* at the Juntas Chicas site, we first performed ancestry analysis to identify any putative hybrids (*25*). This resulted in the exclusion of two individuals that are likely hybrids from our GWAS dataset. For the remaining individuals, reads were mapped to the sparkle *X. evelynae* reference genome with bwa-mem (*26*) and sorted using SAMtools (*27*, *28*). Reads with mapping quality < 30 were removed. Since our data was low-coverage, we performed a case-control GWAS implemented using samtools-legacy to estimate allele frequency differences between sparkle and non-sparkle individuals as we have previously (*14*, *15*). We filtered the output data to remove sites with minor allele frequency less than 5% in both cases and controls, as well as sites with greater than 2X or less than 0.5X average coverage. Results were plotted in R using the qqMan package.

To determine a genome-wide significance threshold for allele frequency differences between the sparkle and non-sparkle individuals, we used a simulation-based approach. For each simulation, we permuted the observed phenotypes such that each individual was randomly assigned as sparkle or non-sparkle. Next, we ran the GWAS with the permuted phenotypes as we had for the real data and recorded the minimum p-value observed. We repeated this procedure 500 times to generate a distribution of minimum p-values expected by chance, since the relationship between genotype and phenotype was determined randomly for each simulation. We then took the lower 5% quantile of this distribution and used this value ($7.4 \times 10^{-10}$) as our genome-wide significance threshold (Fig. 2A).

A region of approximately 2 Mb on chromosome 15 passed our genome wide significance threshold. The peak associated SNP in this region of the sparkle reference was at basepair 11157622 on chromosome 15 with an association p-value of $5.3 \times 10^{-18}$ or $-\log_{10}$(p-value) of 17.3 (Fig. 2). Although we evaluate the entire associated region in our analyses, we considered SNPs with p-values within two orders of magnitude (i.e. two units of the $-\log_{10}$(p-value)) of the peak SNP as the most likely to be associated with the trait. This spanned a ~400 kb region from 10.75 to 11.16 Mb on chromosome 15 in the sparkle reference and from 10.68 to 11.15 Mb on chromosome 15 in the non-sparkle reference. The genes that occur in the larger 2

Mb region and the narrower ~400 kb region are listed in Table S2. We repeated this analysis for the non-sparkle reference and found that our results were qualitatively unchanged (Fig. S30).

*Accounting for population structure in GWAS*

The simple case-control approach that we use for our genome-wide association study allows us to collect and analyze sequence data from hundreds of individuals in a cost-effective way. However, this approach can result in an excess of false positives in the presence of population substructure. While there are standard corrections to control for this inflated false discovery rate, our low coverage data are not well-suited to these methods. Instead, we used a number of analyses to investigate the presence of population structure and evaluate whether it could be impacting the genome-wide significant signal we observe on chromosome 15. To do so, we first generated pseudo-haploid calls for each individual by randomly sampling a read from each position in a pileup file generated with BCFtools (*27*). We assigned the randomly sampled allele as the genotype of that individual at that position and converted sites that were not covered in the individual to NAs. We filtered sites with a minor allele frequency of <5% or where data was missing in >75% of individuals using the recode function in PLINK (http://pngu.mgh.harvard.edu/purcell/plink/; *29*). We next performed PCA using PLINK and plotted our results. We observed no separation between sparkle and non-sparkle individuals in PCA space (Fig. S31) and no significant correlations between phenotype and PCs 1-20.

These results suggest that population structure is unlikely to be impacting our results. However, to directly test this, we used the pseudohaploid calls we had generated to repeat the GWAS using PLINK, incorporating the first 5 PCs as covariates. While we expect this analysis to have reduced power, we still identify a single peak on chromosome 15 strongly associated with sparkle presence or absence (Fig. S30).

*Linkage disequilibrium analysis of SNPs within the GWAS peak*

In plotting results of the GWAS, we noticed that the associated region was larger than we typically identify in *Xiphophorus* species, where linkage disequilibrium tends to decay over ~50 kb in most species studied to date. This prompted us to ask whether there might be unusual patterns of linkage disequilibrium in this region. We first used the pseudohaploid dataset described above and PLINK to compare the decay of linkage disequilibrium over physical distance from the center of our GWAS peak to other regions on chromosome 15. We found that while other regions showed the expected decay of linkage disequilibrium over physical distance, this was not observed for SNPs at the center of the GWAS peak. An example of this pattern is shown in Fig. S32. However, we also did not observe strongly elevated linkage disequilibrium, as we might expect if the region were inverted.

Together with the observation that there were several regions on chromosome 15 where few reads from non-sparkle individuals mapped, this pattern prompted us to collect long-read data from individuals with and without the sparkle phenotype. From these long-read assemblies, we found evidence of an insertion (see Supplementary Materials 5). Using this smaller long-read dataset, we see evidence of high linkage disequilibrium surrounding the insertion (Fig. S33). We expect that the mapping dynamics of short reads to this complex insertion region likely explain the patterns observed in our GWAS dataset.

### Supplementary Materials 7. Gene expression RNA-seq using scale tissue

*Gene expression analysis*

We extracted bulk RNA from lateral line scales of 8 adult *X. evelynae* (4 non-sparkle, 4 sparkle) using the Qiagen RNeasy Mini Kit (Catalog #74106, Qiagen, Valencia, CA). Briefly, fish were euthanized in ice-cold water, spinal cord was severed, and the entire fish was submerged in a petri dish filled with RNAlater. Fine forceps were used to individually pluck all scales from the midline region on both left and right sides of the fish. Scales were placed in 1.5mL tubes filled with RNAlater, transferred to Buffer RLT, and homogenized thoroughly with Monarch kit pestles (NEB #T3002). We followed the manufacturer's instructions for RNA extraction from whole tissue. RNA was quantified using a Qubit and sent to Admera Health Services (South Plainfield, New Jersey) for RNA-seq library preparation and sequencing using the Illumina NextSeq 4000 to collect paired-end 75 bp reads.

To process our samples, we first trimmed adapters and filtered low-quality reads with TrimGalore! (--phred33 --quality 30 -e 0.001 --stringency 1 --length 32 --paired --retain_unpaired; (*30*). To analyze transcript abundance, we used kallisto (*31*), a tool that uses pseudoalignment to estimate transcript abundance. We used kallisto's index command along with the genome annotation file to generate a 'pseudotranscriptome' from the reference genome sequence. We reformatted the *X. evelynae* sparkle reference genome annotation file for compatibility with the DESeq2 software (*32*). We loaded the kallisto abundance files into R to compare expression patterns between samples with DESeq2.

We first created a tx2gene table to match transcript names to gene ID names, and extracted gene-level count information from kallisto counts. Next, we built a model to identify expression differences as a function of phenotype (sparkle or non-sparkle) while controlling for possible batch-effects from RNA extraction and library prep (design = ~ treatment + batch). We used a local dispersion model and generated a PCA of our non-sparkle and sparkle scale samples to identify any outliers. Based on this analysis, we retained all samples in our final analysis. We performed differential gene expression analysis with the lfcShrink() command using an adaptive shrinkage (ashr) method. Results for differentially expressed genes at an adjusted p-value of 0.05 are available in Table S5. We observed 92 genes differentially regulated by phenotype in scale tissue (Table S5).

We then asked whether any genes that were differentially expressed between RNA extracted from scales with and without sparkle fell within our GWAS peak using the program BEDTools (*33*). Out of the 91 total differentially expressed genes, we identified only two genes that fell within the GWAS region: *smyd2* and a gene annotated as *alkal2,* which we confirm is the gene *alkal2a* in the next section. Note that many genes in teleost fish are found in two copies, typically referred to as *a* and *b* copies, due to the teleost whole genome duplication; see next section for more details. Given their colocalization with the GWAS peak, we *considered alkal2a* and *smyd2* as candidates for driving the sparkle phenotype. We note that *alkal2a* falls within the most strongly associated GWAS region from 10.75-11.16 Mb, while *smyd2* does not.

To investigate the proportion of differentially expressed genes with pigmentation related phenotypes, we initially used a gene ontology-based analysis. However, in practice we found that many genes known to be involved in pigmentation in zebrafish, including *alkal2a*, were not annotated as such in PANTHER (https://pantherdb.org/). Instead, we manually searched the zebrafish knockout database to identify phenotypes associated with gene perturbations (https://zfin.org/search). We did this search for 88 (out of 91) differentially expressed genes that had gene symbols associated with their annotations in the *X. evelynae* sparkle reference genome. Based on this search, we identified 27 genes with associated knockout phenotypes in zebrafish (Table S6) and found that 15% of the differentially expressed genes were related to pigmentation phenotypes and that 63% impacted phenotypes derived from the neural crest lineage.

*Confirming the identity of alkal2a with reciprocal best blast, gene tree, and microsynteny analysis*

The gene annotated in the *X. evelynae* reference genome on chromosome 15 as *alkal2* is an excellent candidate for the causal gene driving the sparkle trait. Zebrafish have three ALKAL homologs, also referred to as "FAM150" or "augmentor" (AUG) proteins: ALKAL1 (AUG-β), ALKAL2A (AUG-α1)*,* and ALKAL2B (AUG-α2; (*34*)*.* Previous work demonstrates that *alkal2a* and *b* mediate embryonic iridophore development and adult pigment patterns, while *alkal2b* and *alkal1* are important for eye iridophore formation (*34*). Therefore, we were interested in determining which one is likely homologous to the gene we had identified in *Xiphophorus.*

Searching across the genome, we confirmed that *X. evelynae* has two genes annotated as *alkal2,* one on chromosome 15 and one on chromosome 24. We also identified an additional gene annotated as *alkal-like* on chromosome 9. We extracted the ALKAL2A, ALKAL2B, and ALKAL1 proteins from the zebrafish genome (GRCz11, GCF_000002035.6) and performed blastp to the *X. evelynae* predicted protein sequences. We identified the best blast hit in terms of e-value, performed a reciprocal blastp to the zebrafish protein database, and again identified the best blast hit in terms of e-value. For ALKAL1, we were able to exclude homology to the gene of interest on chromosome 15 in *X. evelynae*. Interestingly, however, the gene on chromosome 15 was the best blast hit for both ALKAL2A and ALKAL2B. This remained the case with the reciprocal blast analysis of the gene on chromosome 15 to the zebrafish genome. In both cases, the gene on chromosome 24 was the second best hit to zebrafish ALKAL2A and ALKAL2B. Aligning the predicted peptide sequences of the *alkal2* copies on chromosome 15 and 24 to zebrafish ALKAL2A and ALKAL2B, we observe some evidence of better alignment of the gene on chromosome 15 to zebrafish ALKAL2A and the gene on chromosome 24 to zebrafish ALKAL2B (in terms of contiguity; Fig. S34).

To confirm that the *alkal2* gene on chromosome 15 was indeed homologous to the zebrafish *alkal2a* copy, we performed a gene tree and microsynteny analysis. First, we downloaded known *alkal2a* and *alkal2b* protein sequences from NCBI for *Danio rerio*, *Takifugu rubripes*, and *Poecilia formosa*, and known *alkal2* protein sequences from *Homo sapiens* and *Mus musculus*. We aligned these protein sequences and the predicted protein sequences for chromosome 15 and 24 genes from *X. evelynae* using MAFFT version 5, building a neighbor joining tree using the 86 conserved sites in this alignment (*35*). We rooted the tree by the branch leading to the *alkal2* copies in *H. sapiens* and *M. musculus*, which formed a monophyletic group. We found that the *alkal2a* and *alkal2b* sequences formed monophyletic clades, excluding the *D. rerio alkal2b* sequence which clustered basal to the group (Fig. S35). This pattern has been previously observed for ancient gene duplicates (*36*). Based on this analysis, the chromosome 15 gene from *X. evelynae* clusters with *alkal2a* and the chromosome 24 gene clusters with *alkal2b*.

As orthogonal evidence, we evaluated patterns of microsynteny (i.e. the order of nearby genes). We extracted genes within 150 kb of *alkal2* paralogs from our *X. evelynae* reference assemblies and in *D. rerio* from the ZFIN database. We found that gene order was conserved for the genes directly upstream and downstream of *alkal2a*, even across >200 million years of divergence (Fig. S35). Moreover, for *alkal2b*, *X. evelynae* and *D. rerio* also shared the closest gene: *sh3yl1* (Fig. S35).

*Evaluating likely cell-type origin of differentially expressed genes based on zebrafish databases*

The most closely related species with extensive single-cell datasets to *Xiphophorus* is zebrafish. Thus, we used the ZebraHub tool from CZ Biohub to investigate the cell-type specific expression of differentially expressed genes in zebrafish (https://zebrahub.sf.czbiohub.org/; *37*). First, we took the 91 differentially expressed genes between sparkle and non-sparkle scales in our RNA-seq dataset and searched for them in the zebrafish Neural Crest lineage transcriptomics dataset. We then examined whether each gene with a corresponding zebrafish annotation (explained below) was expressed primarily in iridophore cells. As previously discussed, teleosts experienced a whole genome duplication in their common ancestor, and paralogs that originated from this event are typically referred to as *a* and *b* copies if they are both retained in the genome. Given that zebrafish and swordtails have some differences in the retained orthologs from the teleost whole genome duplication, we considered any gene with a shared annotation in this analysis, regardless of whether it was annotated as an *a* or *b* copy in either species.

We found that of our 91 differentially expressed genes, *fhl2a/fhl2b*, *pnp4a*, *alx4a/alx4b*, *apoda.1/apodb*, and *sytl2a* are all specifically expressed in iridophore cells in zebrafish (Fig. S36). We also observed that *pnp5a*, *arhgef33*, *fmn2a*, and *sytl2b* are expressed in iridophores and a subset of other cells including xanthophores, retinal pigmented epithelium, and a group of unassigned cells. This suggests that many of the differentially expressed genes are iridophore associated or originate from a similar developmental lineage (Fig. S36).

**Supplementary Materials 8. Real time qPCR to determine gene expression**

To confirm the association between *alkal2a* upregulation and sparkle trait formation, we conducted qPCR on scale tissue extracted from both sparkle and non-sparkle fish at three developmental stages: 1) fry stage: before trait onset at 3 weeks old, 2) juvenile stage: during trait development, and 3) adult stage: post trait development and post-sexual maturation. Although our RNA-seq dataset revealed *alkal2a* as the only differentially expressed gene within the most significant GWAS region, we investigated whether any other genes within the GWAS peak were known to interact with *alkal2a* using STRINGdb (*38*). We looked to see which genes are known to interact with the homolog of ALKAL2A in humans, ALKAL2, since this database was the most complete. We found that FAM110C and ACP1, which both occur in the most significant GWAS region, are inferred to interact with or be coexpressed with ALKAL2 (Fig. S37). FAM110C is inferred to have a direct interaction and ACP1 is inferred to have a second order interaction with ALKAL2, potentially hinting at a functional link. As a result, we evaluated whether either of these three genes are differentially regulated by phenotype during development.

Non-sparkle and sparkle fish were euthanized and examined under a dissection microscope to confirm sparkle trait presence or absence when applicable. We used forceps to remove all lateral line scales from the left and right flanks in both sparkle and non-sparkle fish. Scales were immediately stored in either RNAlater or flash frozen. For fry that had yet to develop the sparkle trait (e.g. individuals 3 weeks of age), we similarly removed scales from the lateral line where sparkle typically develops first. We collected a matched fin clip for DNA extraction and genotyping (see below section: *PCR and qPCR to determine insertion copy number*).

RNA was extracted and reverse transcribed to cDNA using the GoScript reverse transcription kit (Promega) following the manufacturer's instructions. Primers for qPCR were designed using the predicted cDNA sequence of the gene of interest and the PrimerBlast tool from NCBI. Primers were designed for *alkal2a*, *acp1*, *fam110c*, and *efa1-alpha* (housekeeping gene). We confirmed that primers amplified a single product matching the expected target size using the Phusion High-Fidelity PCR kit before proceeding with efficiency testing for qPCR. Primer sequences are available in Table S7.

For qPCR amplification, each reaction contained 5 µl of iQ SYBR mix, 0.3 µl of the forward and reverse primers, 3.4 µl of water, and 1 µl of cDNA. All samples were run in triplicate. Reactions were run in a 384-well plate format on a Bio-Rad CFX384 Touch Real-Time PCR System. We determined qPCR efficiency for each primer set using a 6-set serial dilution approach with each replicate run in triplicate. We estimated that primers for e*fa1-alpha* had an efficiency of 101.1%, *acp1* at 109.7%, *fam110c* at 101.7%, and *alkal2a* at 99.8%. Following efficiency testing, RNA was extracted from adults (6 non-sparkle, 9 sparkle), juveniles (6 non-sparkle, 6 sparkle), and fry (3 non-sparkle, 3 sparkle) using the QIAquick RNA extraction kit. As described above, samples were reverse transcribed and qPCR was conducted using the iQtm SYBR Green Supermix kit. Average Ct for each biological replicate was calculated from 3 technical replicates per fish. Fold change was calculated by the $2^{-\Delta\Delta Ct}$ method (with the focal gene normalized to *efa1-alpha* expression).

Given that our qPCR data did not appear to be normally distributed, we performed a Kruskal-Wallis test between non-sparkle and sparkle phenotypes for each life stage to test for significant differences in expression. We found that *alkal2a* was differentially expressed between sparkle and non-sparkle fish at all life stages (see main text; fry: p-value=0.0495, juvenile: p-value=0.0104, adult: p-value=0.00146). By contrast, *acp1* and *fam110c* were not differentially expressed between sparkle and non-sparkle individuals at any developmental stage evaluated (Fig. S38; *acp1* fry: p=0.827, juvenile: p=0.513, adult: p=0.439; *fam110c* fry: p=0.513, juvenile: p=0.513, adult: p=0.302; based on a Kruskal-Wallis test).

*PCR and qPCR to determine insertion copy number*

For individuals too young to express the sparkle trait, we were nonetheless interested in determining their genotype so that we could investigate gene expression patterns that preceded the onset of the trait. We were able to generate highly specific primers for the sparkle haplotype (Table S7) by amplification of a sequence that spans the edge of the 3' LTR of ERV-F.1-Xeve-chr-15 and the *MG3f* sequence. Thus, these primers are expected to be specific to the insertion on chromosome 15 and not amplify other ERV-foamy-Xeve elements in the genome since our analyses suggest that other ERV-foamy-Xeve elements lack the *MG3f* insertion (see Fig. 4).

We demonstrated that non-sparkle individuals consistently lacked amplification whereas all sparkle individuals amplified this sequence, verifying the primer performance on 18 sparkle and 11 non-sparkle adults. The complete absence of amplification in non-sparkle individuals is expected given our inference from sequencing data that the sparkle haplotype is dominant. Thus, amplification with these primers distinguishes between non-sparkle and sparkle individuals and confirms that the insertion of the TE is present across a larger sample of sparkle individuals than was tractable to sequence using PacBio technology. Importantly, these primers also allow us to identify fry that are genetically predicted to be sparkle or non-sparkle as adults before they develop the trait. This allowed us to quantify expression of *alkal2a* and other genes in fry before the trait develops, as described in the previous section.

We were further interested in whether we could use qPCR on genomic DNA to distinguish between individuals that are heterozygous versus homozygous for the sparkle haplotype. We reasoned that we could make this distinction by comparing qPCR amplification profiles of genomic DNA relative to a conserved single copy control gene. We chose Nup43, which is a highly conserved BUSCO single copy gene that we have previously used as a single copy control gene (*39*). If this approach was successful, we expected to see three classes of expression in our delta Ct results. Non-sparkle individuals should have normal amplification of Nup43 but fail to amplify primers for the sparkle haplotype. Heterozygous individuals should have a lower Ct for Nup43 than for the sparkle haplotype, with an expected delta Ct of 1 given that these individuals harbor two copies of Nup43 in their diploid genome and one copy of the sparkle haplotype. Homozygous individuals show an equivalent Ct for Nup43 and the sparkle haplotype given that they should have two copies of both Nup43 and the sparkle haplotype in their diploid genomes.

Examining the distributions of delta Ct for this experiment across sparkle and non-sparkle individuals, we clearly saw that non-sparkle individuals had high Ct values (e.g. >30) for the primer set diagnostic of sparkle but had amplification at Nup43, resulting in large delta Cts between the primer sets, as expected. Among sparkle individuals, delta Ct generally fell into two classes consistent with our expectations: one with delta Ct <0.5, which we infer to be homozygous, and one with delta Ct between 0.5 and 1.5, which we infer to be likely heterozygous individuals (Fig. S39). However, since these distributions were somewhat overlapping, we did not consider these results precise enough to test for downstream associations between TE copy number and gene expression or phenotypic associations (e.g. dosage effects or sparkle area).

### Supplementary Materials 9. Pharmacological inhibition of *ALK* in sparkle scales

ALK inhibitor (NVP-TAE 684, MedChem Express) was diluted in DMSO to make a stock solution at 160 µM. A 2% ALK inhibitor solution was made in cell culture media (L15+20% FBS +15mM HEPES). Control samples were cultured in a 2% DMSO solution in cell culture media. We added either 100 µL of the control media or ALK inhibitor media to individual wells of a 96-well plate (clear flat bottom). Surrounding wells were filled with water to prevent evaporation. Fish were anesthetized in MS-222 and scales were plucked from the left and right side of the lateral line of a sparkle fish and individually placed in the wells. Scales were immediately imaged on the Cytation5 and incubated for 20 hours at room temperature, then imaged again to validate any reduction in iridophore cell populations.

To track changes in iridophore morphology in response to the inhibitor, we repeated this procedure of imaging sparkle scales approximately every hour for 35 hours after the scales were submerged in the 2% ALK inhibitor solution. During this time, we saw iridophore cells lose their typical morphology and shrink into puncta, while we observed no changes in iridophore morphology in our DMSO control (Fig. S6).

**Supplementary Materials 10. Analyses of ERV-F.1-Xeve-chr-15 insertion**

*Classification of the virus insertion and genome structure annotation*

As described in the main text, we identified an inserted sequence on chromosome 15 with sequence similarity to endogenous retroviruses. To classify this sequence, phylogenetic analysis was performed based on the reverse transcriptase (RT) proteins of the virus insertion and representative exogenous and endogenous retroviruses as described previously (*40*, *41*). Sequences used in this search can be found on Dryad (doi: pending). RT proteins were aligned using MAFFT 7.505 (*42*). Phylogenetic analysis was performed using the maximum likelihood method implemented in IQ-tree 2 (*43*). ModelFinder was used to select the best amino acid model (*44*). The ultrafast bootstrap approximation (UFBoot2) was used to calculate statistical support values for nodes with 1000 replicates (*45*, *46*).

The genome organization of the insertion of interest, ERV-F.1-Xeve-chr-15, was annotated using ORFfinder, HHpred (*47*, *48*), TMHMM (*49*, *50*), BLASTp, and CD-search (*40*).

*Identification of ERV foamy elements in Poeciliidae*

We were also interested in whether there were other full length or partial endogenous foamy viruses in the *X. evelynae* genome or in closely related species. The program LTRharvest (*51*) was used to mine sequences of >5,000 bp flanked by LTRs in the 11 haplotypes of *X. evelynae*. The tBLASTn algorithm was used to search homologs of RT proteins. Endogenous foamy viruses (EFVs) are the genus of ERVs to which the chromosome 15 element belongs. Additional EFVs were then identified based on phylogenetic analysis of RT hits and RT proteins of representative retroviruses (*40*, *41*). RT proteins were aligned using MAFFT 7.505 (*42*). Phylogenetic analysis of the resulting alignments was performed using IQ-tree 2 (*43*).

The tBLASTn algorithm was used to search against the genomes of Poeciliidae species with the RT protein of ERV-F.1-Xeve-chr-15 as the query and an *e*-value cut-off of $10^{-5}$. For a list of genomes searched, see Table S8. EFVs were then identified based on phylogenetic analysis of RT hits and RT proteins of representative retroviruses as described previously (*40*, *41*). The RT nucleotide sequences of EFVs identified in the genomes of Poeciliidae species were aligned using MAFFT 7.505 (*42*) with L-INS-i strategy. The alignment was refined using trimAl v1.4 (*7*). Phylogenetic analysis was performed using IQ-tree 2 as described above (*43*).

Over evolutionary timescales, it is common for recombination to occur between the LTRs of ERV sequences, resulting in the emergence of "solo-LTRS". To search for these sequences in Poeciliidae species, we used BLASTn to identify solo-LTRs with the LTRs of EFVs of *X. evelynae* used as the query sequence and an *e*-value cut-off of $10^{-5}$.

*Evolutionary dynamics of ERV foamy elements in X. evelynae*

The LTRs of complete EFVs were aligned using the L-INS-i strategy in MAFFT (*42*). The genetic distance of 5'- and 3'-LTR flanking a complete EFV was calculated using a Kimura two-parameter substitution model (*52*). To establish whether there were orthologous relationships between EFV sequences represented in *X. evelynae*, the 1000 bp sequences flanking EFV LTRs were retrieved and subject to all-to-all BLASTn with an *e*-value cut-off of $10^{-5}$. Orthologous EFVs share highly similar flanking sequences outside of LTRs.

### Supplementary Materials 11. Analysis of genetic variants associated with the sparkle phenotype

*Analysis of MG3f element across X. evelynae genome assemblies*

Analysis of the insertion on chromosome 15 revealed that it contains a complete ERV sequence and a fragment of an MG3-domain-containing gene, which we refer to as *MG3f* (Fig. 3B). We first evaluated all other sparkle haplotypes and confirmed that the 255 bp *MG3f* sequence was always inherited with the ERV insertion on chromosome 15—this was also confirmed by qPCR with our sparkle haplotype primers, since the reverse primer was designed to the *MG3f* sequence. Next, we used blastn with an e-value threshold of $10^{-20}$ to identify homologs of *MG3f* in the *X. evelynae* haplotypes we had assembled. We determined that all *X. evelynae* individuals have a sequence with >99.5% identity to *MG3f* on chromosome 5, but only sparkle individuals have this sequence on chromosome 15 (directly downstream of ERV-F.1-Xeve-chr-15). The sequence on chromosome 5 falls within the MG3-super family gene C3 and overlaps with an entire exon, entire intron, and part of a second exon (Fig. 4). C3 is a large gene with 42 annotated exons on chromosome 5 in the *X. evelynae* reference genome. We also used the same approach to investigate its presence in other *Xiphophorus* species and found that it is present as a single copy gene on chromosome 5 in all other species with available genome assemblies. These results suggest that *MG3f* might be originally derived from the *X. evelynae* C3 sequence on chromosome 5. However, our analysis of available *X. evelynae* genomes indicates that there are not ERV foamy element insertions near this gene on chromosome 5, nor do other ERV foamy element insertions in the genome contain *MG3f*-like sequences. Therefore, the sequence of events that led to the insertion of ERV and *MG3f* on chromosome 15 are, as of yet, unclear. Nevertheless, we evaluated whether the *MG3f* sequence may act as a *cis* regulatory element in downstream analyses and experiments.

*Analysis of variable region downstream of the ERV insertion*

Analysis of genetic variation within the GWAS interval revealed a ~3 kb intronic region with high SNP-density located 1.7 kb downstream of the ERV-F.1-Xeve-chr-15 insertion. We refer to this region as the "hypervariable" region here and in the manuscript. This region included 23 single nucleotide polymorphisms (SNP) within a 3.2 kb interval that distinguished sparkle and non-sparkle haplotypes in our dataset. We refer to this variable region as "SPvar" in the sparkle haplotype and "NSPvar" in the non-sparkle haplotype (Fig. 3G). This region has unexpectedly high divergence between the sparkle and non-sparkle haplotypes (0.7% compared to ~0.1% genome-wide). We treat this region as a candidate *cis* regulatory element in downstream analyses and experiments (see below).

The high level of differentiation in this region suggests that it is either a rapidly evolving intronic sequence or that it is derived from hybridization or incomplete lineage sorting. To investigate these possibilities, we performed a number of analyses. First, we extracted this region and flanking regions from all long-read based assemblies available to us, resulting in 31 sequences from 14 species. We used Clustal Omega (*53*) to align these sequences, resulting in an alignment of 10,316 basepairs. Next, we calculated pairwise divergence between all haplotypes and used this alignment as input into RAxML to infer phylogenetic relationships under a GTR+GAMMA model of sequence evolution with 100 rapid bootstraps. We rooted the phylogeny using the branch that separated the platyfish clade, including *X. evelynae*, and the Northern swordtail clade. The resulting phylogeny highlights phylogenetic discordance within the platyfish clade. While the Northern swordtail clade generally follows patterns consistent with the species tree (*1*), branching patterns in the platyfish clade do not. The *X. evelynae* sparkle and non-sparkle haplotypes each form monophyletic clades, but the non-sparkle haplotypes cluster with *X. xiphidium* haplotypes with high confidence (Fig. S40). We speculate that this is likely to be the result of incomplete lineage sorting or ancient gene flow for two reasons. First, the region that shows these patterns is very short (~3 kb) and is thus inconsistent with recent gene flow. Second, the pairwise sequence divergence between *X. xiphidium* and non-sparkle *X. evelynae* haplotypes in this region is quite high (~2%), again inconsistent with recent gene flow. Together, this makes incomplete lineage sorting (or ancient gene flow) a likely cause of this region of elevated divergence.

We next wanted to evaluate whether this region could be the true causal driver of the sparkle phenotype. In other words, we wanted to consider the possibility that this variable region was the causal driver of the sparkle trait but became associated with the ERV-F.1-Xeve-chr-15 via by its chance insertion in the same genomic region. Based on our data, we consider this less likely than the scenario we outline in the main text for several regions. First, given that this is an intronic sequence, if it were associated with differential *alkal2* expression, we expect this would be via a regulatory mechanism. However, we see no evidence of differential chromatin accessibility between sparkle and non-sparkle samples in this hypervariable region (Fig. S15). Second, we do not see differences in the number or locations of predicted *sox10* or *tfec* binding sites between the two versions of the hypervariable region (Fig. 3H). Third, we describe below experiments testing the *cis* regulatory potential of these two versions of the hypervariable region and the LTR sequence using a luciferase assay (see Supplementary Materials 15). Surprisingly, we find that the non-sparkle hypervariable sequence drives higher reporter activity *in vitro* than the sparkle hypervariable sequence, which is the opposite of what we would expect if this region was important in regulating the sparkle trait (Fig. 3G). Moreover, the LTR sequence derived from the ERV-F.1-Xeve-chr-15 insertion drove reporter activity more strongly *in vitro* than either the sparkle or non-sparkle hypervariable regions, consistent with our hypothesis that the ERV insertion is the true driver of the difference in *alkal2a* expression associated with the sparkle trait (Fig. 3).

### Supplementary Materials 12. Collection of RNA-seq data from brain and testes for analysis of ERV-foamy expression

While the host genome often uses mechanisms such as 5mC methylation to suppress TE activity (see next section), past work has indicated that active endogenous retroviruses can be expressed in both somatic and germline tissues (*54*, *55*). To evaluate evidence of expression of the ERV-foamy-Xeve elements, we collected bulk RNA-seq data from brain and testis tissues. As described above, we extracted bulk RNA from testes and brain of *X. evelynae* individuals using the Qiagen RNeasy Mini Kit (Catalog #74106, Qiagen, Valencia, CA). Briefly, fish were euthanized with an ice-bath, photographed, and their spinal cord was severed. Fish brain and testes were immediately dissected and flash-frozen in 1.5 mL tubes. We homogenized tissue thoroughly with Monarch kit pestles (NEB #T3002) and followed manufacturer's instructions for the RNA extraction from whole tissue. We measured RNA concentrations by Qubit and sent samples for library preparation and sequencing at Admera Health Services (South Plainfield, New Jersey). We generated data from 8 brain and 8 testis samples (from four non-sparkle and four sparkle fish) on the Illumina NextSeq 4000 to collect paired end 75 bp reads.

We used trim_galore (--phred33 --quality 30 -e 0.001 --stringency 1 --length 32 --paired --retain_unpaired) to remove Illumina adapters and filter low-quality reads (*30*). Given the repetitive nature of ERV-foamy-Xeve elements in a genome, we required a unique approach to analyze evidence of TE expression. Annotated EFVs in our assemblies vary in sequence similarity to each other from moderate (~90%) to high (>95%). Moreover, all but one of the full-length ERV-foamy-Xeve sequence detected in our assemblies are polymorphic (Fig. 3E), and many of these elements occur at low frequencies. This suggests that there are likely ERV-foamy-Xeve sequences present in the individuals from which we collected RNA-seq data that are not annotated in our long-read haplotype assemblies. Therefore, evidence of RNA-seq reads that map to the ERV-foamy-Xeve sequences suggests that EFV family is transcribed and active but does not demonstrate of expression of individual elements.

To generally test whether ERV-foamy-Xeve elements are actively transcribed in the genome, we performed an analysis where we sequentially masked all ERV-foamy-Xeve sequences annotated in our sparkle reference genome except for a single focal sequence. We then mapped all RNA-seq reads to each version of the genome using STAR (--outFilterMultimapNmax 150, --winAnchorMultimapNmax 150; *56*), which retains multi-mapped reads. We used BEDTools (*33*) to intersect our BAM files to regions of interest for analysis, which included annotated ERV-foamy-Xeve sequences and control regions. To generate matched control regions, we first excluded the coordinates of all protein coding genes as well as regions within 5 kb of these sequences. We then excluded regions with annotated TE sequences since we do not have a catalog of which TEs are expected to be active in *Xiphophorus* genomes. Finally, we randomly selected one hundred regions as control regions to compare with expression levels of ERV-foamy-Xeve elements in the brain and testes. Note that we expect *a priori* that the control regions should be less repetitive than ERV-foamy-Xeve sequences, and thus more likely to map uniquely, but that this should be conservative for the purposes of our analysis.

We calculated RNA-seq coverage in 100 bp windows in each of our focal and control regions for brain and testis samples respectively using Sambamba (*57*). We normalized coverage to the total number of mapped reads. To compare expression of the ERV-foamy-Xeve sequences to other regions of the genome which we do not expect to be expressed, we compared normalized expression in the ERV-foamy-Xeve sequences to control regions across samples.

Finally, we plotted coverage at each ERV-foamy-Xeve and example null region from the BAM files in both brain and testes samples. We present these results, focusing on the analysis where ERV-F.1-Xeve-chr-15 is unmasked in Fig. S12. We found evidence for low levels of expression of the ERV-foamy-Xeve sequences in both brain and testes tissue, but expression in these regions exceeded levels observed across most of the control regions (Fig. S12). For analyses in which ERV-F.1-Xeve-chr-15 was masked and other ERV-foamy-Xeve sequences were unmasked, we generally observed similar results, although several elements had few mapped reads (see examples in Fig. S12).

**Supplementary Materials 13. Analysis of methylation signals at the sparkle insertion and elsewhere in the genome**

Since one of the primary mechanisms for repression of TE activity is CpG methylation (*58*), we were interested in whether ERV-foamy-Xeve sequences showed signals of 5mC methylation at the chromosome 15 insertion site and elsewhere in the genome. Information on methylation at CpG sites can be extracted from PacBio data. We first mapped individual PacBio reads either to 1) the sparkle reference genome (containing ERV-F.1-Xeve-chr-15) and/or to 2) the genome assembled from their corresponding PacBio reads. We removed reads with a mapping quality less than 30. Using these bam files and the software pb-CpG-tools (https://github.com/PacificBiosciences/pb-CpG-tools), we used the --pileup-mode to generate modification scores at each site where 5mC methylation is possible. We calculated average discretized modification scores at CpG sites from our data in windows of 10 CpG sites.

After we had generated a modification score output for each individual with both mapping approaches, we analyzed the data relative to the locations of annotated ERV-foamy-Xeve insertions. Near ERV-F.1-Xeve-chr-15, we found that most non-sparkle individuals had highly divergent, and likely mismapped, reads in this region of the genome. These presumably originated from ERV-foamy-Xeve elements not annotated in our sparkle reference genome. Accordingly, for analysis of sequences mapped to the sparkle reference genome, we limited our analysis to the sparkle individuals. In doing so, we found evidence for elevated 5mC methylation across ERV-F.1-Xeve-chr-15 in sparkle individuals (Fig. 3I-J).

We were also interested in whether other complete ERV-foamy-Xeve insertions had similarly high levels of 5mC methylation. To analyze this for each of our long-read assembly haplotypes, we took the coordinates of complete ERV-foamy-Xeve insertions in their genome and analyzed 5mC methylation in self-mapped PacBio reads in these regions. We followed the analysis approach described above to quantify 5mC methylation and generally found evidence for locally elevated levels of 5mC methylation along the distinct ERV-foamy-Xeve insertions (Fig. S13).

### Supplementary Materials 14. Cell dissociation and ATAC-seq on scale tissue

*Sample preparation*

Fish were euthanized using an ice-bath followed by spinal cord severing, in accordance with Stanford APLAC protocol 33071. Fish were imaged, and their entire flanks were immediately descaled while soaking in 1XPBS/1%BSA on ice. Scales were placed into 35 mm petri dishes filled with ice-cold 1XPBS/1%BSA. All liquid was removed, 1 mL of 100U/mL Type 2 collagenase (Worthington Biochemical Corporation) was added to the petri dish, and the scales were incubated at 35°C for 30 minutes. Collagenase was removed and pre-warmed 1X TrypLE Express Enzyme (Thermo Fisher Scientific) was added to each dish, incubating at 35°C for 20 minutes with a 10x trituration step every 5 minutes. Then, all liquid was filtered with a 40-micron filter (Scienceware® Flowmi™ Cell Strainers for 1000µl Pipet Tips, Bel-Art, Porosity:40 µm, Sigma-Aldrich) into a 15 mL conical tube. This sample was centrifuged at 300xg for 5 minutes at 4°C and a cell pellet was visually confirmed. The supernatant was aspirated, and the pellet was vigorously resuspended in ~100 µL ice-cold 1XPBS/1%BSA. Live cell counts were determined by staining with Trypan blue and counting unstained cells with Invitrogen's Countess 3 Automated Cell Counter instrument.

Next, we proceeded to nuclei isolation and library preparation. 50 µL of this cell solution was added into a small, glass douncer (KONTES Dounce Tissue Grinders, Kimble Chase – 7 mL), and 100 µL of EZ lysis buffer was added before douncing 10x with pestle A and 10x with pestle B. Liquid was transferred into a 5 mL tube, incubated on ice for 2 minutes, and strained through a 40-micron filter. Nuclei were centrifuged at 500xg for 5 minutes at 4°C, the supernatant was aspirated, and nuclei were resuspended in 50 µL cold 1x PBS. We evaluated the number and quality of isolated nuclei with a cell counter and with images taken on the Agilent Cytation5 machine. The sample was diluted to ~25,000 nuclei per sample using PBS. Samples were tagmented with the Tn5 enzyme and cleaned either with a bead-based or column-based approach. Tagmented samples were combined with indexed primers and PCR mastermix for PCR amplification for 5 cycles. Additional amplification steps were performed by running a qPCR on a light cycler to calculate the number of additional cycles needed to generate half the total amplitude of the qPCR amplification plateau. The required number of additional PCR cycles were performed for each sample. Finally, a bead- or column-based cleanup was performed on each library, which was then analyzed on an Agilent Tapestation to confirm that the size and distribution matches that of a typical ATAC-seq library. Samples were sent to Admera Health Services for sequencing at a total depth of 150-200 million paired-end 150 basepair reads on a NextSeq1000 P1. A subset of samples were resequenced on a NovaSeq 10B to collect higher coverage.

*Pre-processing and quality control analysis of ATAC-seq data*

To prepare ATAC-seq libraries for analysis, we first used the program Cutadapt to trim possible adapter sequences and reduce read length to 75 bp (*59*). We then mapped reads to a modified version of the sparkle reference genome (see below) using bwa-mem (*26*). We marked duplicates and realigned indels with PicardTools (https://broadinstitute.github.io/picard/) and removed reads mapping to the mitochondrial genome with ngsutilsj (https://github.com/compgen-io/ngsutilsj; *60*). We visualized mapped reads in IGV and calculated average coverage in sliding windows using Sambamba (*57*).

To evaluate the quality of our ATAC-seq data collection at a genome-wide scale, we used several approaches. First, we analyzed BAM files generated as described above with PicardTools CollectInsertSizeMetrics to generate a plot of the length distribution of each sample. All libraries showed the expected insert size profile for ATAC-seq libraries (Fig. S41). We next used the ATACseqQC package in R to evaluate library complexity, calculate promoter/transcript body scores, and the transcription site enrichment score (Fig. S41). Based on these analyses, we determined that our libraries were successful and useful for analyzing chromatin accessibility differences as a function of phenotype.

In addition to these genome-wide signals, we were interested in examining local patterns of accessibility in promoter regions known to be highly active across tissues. Past RNA-seq experiments have identified genes that are constitutively active across tissues (*61*), and a subset of the 1 kb upstream regions have been functionally verified by our group to drive GFP expression in *Xiphophorus* cells. We examined patterns of chromatin accessibility in these regions as a positive control and found that as expected they had consistently high levels of accessibility across samples (Fig. S17).

*Analysis of ATAC-seq data*

We were interested in investigating the hypothesis that sparkle individuals with the ERV-F.1-Xeve-chr-15 insertion near *alkal2a* have increased chromatin accessibility in this region. We were specifically interested in the LTR regions, which have been frequently shown to act as enhancers. Specifically, the 5' and 3' LTR sequences of the ERV-F.1-Xeve-chr-15 insertion are identical, such that if both are included, all reads mapping to these LTRs will receive a mapping quality score of 0. We thus performed two separate analyses in which we masked either the 5' LTR or the 3' LTR of ERV-F.1-Xeve-chr-15. We present the 3' LTR analysis in the main text, but our results are qualitatively unchanged with the 5' LTR analysis.

In addition to this issue, there are several genetically similar ERV-foamy copies throughout the *X. evelynae* genome (which we refer to as ERV-foamy-Xeve sequences) as well as over 100 genetically similar full or partial LTR sequences throughout the *X. evelynae* genome. This meant that we required a unique approach to examine chromatin accessibility in this region. We reasoned that even if short-read sequences originating from other regions of the genome mismapped to ERV-F.1-Xeve-chr-15, we should not expect them to correlate with the sparkle phenotype and thus they should not erroneously drive ATAC-seq peaks associated with the sparkle phenotype. While there are some SNP differences between the LTRs in ERV-F.1-Xeve-chr-15 and the genetically similar LTRs spread throughout the genome, we expected that many reads would not map uniquely. When reads map with similar probability to multiple regions of the genome they receive a low mapping quality score. Given these features of our data, this necessitated a distinct analysis strategy where we included reads with poor mapping quality (<20) in our analysis. We then calculated average coverage using Sambamba (*57*) with a mapping quality cutoff of >0 in sliding 100 basepair windows along the genome (step size 50 basepairs). We visualized results for each sample at our focal region in IGV and in R (Fig. 3F) and calculated enrichment of mapped reads in this region relative to the genome wide background. We note, however, that low mappability also raises issues with interpreting enrichment relative to the genome-wide background.

Because of the unique challenges of analyzing this region, in addition to comparing coverage in our focal region to median coverage genome-wide, we also performed a targeted analysis of genetically similar LTR regions as a control. We identified 80 regions across the genome that were ≥95% identical to the LTRs from ERV-F.1-Xeve-chr-15 and were ≥60% of its length (i.e. at least 1 kb in length). These regions had the added benefit of being similar in GC content to the focal LTRs which fall in the 75% quantile of local GC content genome-wide (all 80 comparison regions were within 5% of the GC content of the LTRs from ERV-F.1-Xeve-chr-15). For each of these 80 regions, we quantified coverage in sliding 100 bp windows as described above, calculated average coverage and mapped read enrichment across the entire region. We also visualized each region in IGV and used a t.test to compare normalized coverage in sparkle and non-sparkle samples.

The vast majority of these control LTRs had very low average coverage in our ATAC-seq data, with 50% of regions ≤0.5X per basepair coverage and 75% of regions ≤0.75X. Only a handful of control LTRs had a substantial number of mapped reads in any sample. All control regions with normalized LTR coverage ≥3X in any sample are shown in Fig. S18. Ninety percent of regions had average normalized coverage less than 2X and examples of these regions are shown in Fig. S19. By contrast, in the focal LTR region on chromosome 15, sparkle samples had a coverage enrichment of 1.7-6X in the LTR regions of ERV-F.1-Xeve-chr-15 (average 3X). Non-sparkle samples ranged from 0.25-0.6X coverage in this region relative to the genome-wide background (average 0.45X). Based on the comparison to the control LTRs, enrichment at ERV-F.1-Xeve-chr-15 in sparkle individuals ranged from 17-60X. By contrast, enrichment in coverage signal at ERV-F.1-Xeve-chr-15 in non-sparkle individuals compared to other LTRs ranged from 1-4X.

We also evaluated possible differences in chromatin accessibility at other nearby regions that distinguish sparkle and non-sparkle individuals, including the MG3 fragment (*MG3f*) and the hypervariable region flanking the insertion sites of ERV-F.1-Xeve-chr-15. We did not find evidence for increased accessibility between sparkle and non-sparkle individuals in *MG3f* or the hypervariable region (Fig. S15), in the promoter, coding, or intronic regions of *alkal2a* (Fig. S15), or in other structurally variable regions within the most significantly associated GWAS peak (Fig. S16). These data suggest that the LTR sequence of ERV-F.1-Xeve-chr-15, but not other nearby sequences, was likely to have *cis* regulatory function, a hypothesis we consider further in reporter assays below.

**Supplementary Materials 15. Luciferase cell reporter assay**

To test whether the LTR sequence or other sequences associated with the sparkle phenotypes had the capability to act as a *cis*-regulatory element, we designed a luciferase cell reporter assay. Because the LTR sequence is present in multiple copies in the *X. evelynae* genome, we synthesized the sequence rather than preparing the insert with a PCR-based approach. We included XhoI and KpnI restriction enzyme cut sites in the synthesized sequence from IDT and used the dual-reporter luciferase assay kit from Promega Biosciences. We digested the pNL3.2 NlucP vector from Promega Biosciences and the LTR insert plasmid from IDT with XhoI and KpnI-HF enzymes at 37°C overnight. We then ran the LTR insert plasmid digested product on a 1% agarose gel and extracted the 1.6 kb LTR insert band under a UV imager and performed a gel extraction with the QIAquick Gel Extraction Kit, following the manufacturer's instructions. Using the Zymo Clean and Concentrator™-5 kit, we purified the pNL3.2 vector and the 1.6 kb LTR insert. We treated the pNL3.2 vector with Antarctic Phosphatase to prevent spontaneous re-ligation of the vector backbone and repeated another Zymo Clean and Concentrator™-5 kit on the pNL3.2 vector. We performed an overnight ligation at 16°C using the T4 DNA ligase at the recommended 3:1 ratio (insert:vector). The next day, we heat-inactivated the ligation reaction at 65°C for 10 minutes and transformed our ligated product into One Shot Stbl3 Chemically Competent cells (Invitrogen), using Puc19 as a positive control and nuclease-free water as a negative control. We plated our transformed cells on LB agar plates with 100ug/mL concentration carbenicillin and grew them in LB at 37°C overnight. We pelleted the resulting culture at 3,400xg for 10 minutes at 4°C and used a ZymoPURE™ Plasmid Miniprep kit following the manufacturer's instructions. We confirmed that the plasmid sequence was correct with sequencing by Plasmidsauraus. We then cultured more bacteria for a maxiprep to have sufficient amount of plasmid for the luciferase reporter assays detailed below.

We repeated this procedure to generate versions of the pNL3.2 vector containing *MG3f*, sparkle variable, and non-sparkle variable inserts, except that we generated these inserts by PCR with XhoI and KpnI cut site overhangs added to the primers. We included these additional regions in the luciferase assay to test the hypothesis that these other regions, rather than the ERV-foamy insertion, may be important for driving *alkal2a* upregulation in sparkle individuals.

To perform the luciferase reporter assay, we first thawed, passaged, and trypsinized HEK 293T cells, plating ~6.25 x $10^5$ cells per well in a 6-well plate with 2.5 mL of Dulbecco's Modified Eagle Medium (DMEM) growth medium (Gibco™, DMEM, high glucose, GlutaMAX™ Supplement, pyruvate) the night before transfection, aiming for >60% confluence in each well on the day of transfection. We followed the *Trans*IT-LT1 Transfection Reagent (Mirus Bio) protocol to transfect the empty NanoLuc plasmid pNL3.2 or the experimental constructs with a control firefly pGL4.54 luc2/TK into wells. We transfected 3 wells of the empty pNL3.2 plasmid and 3 wells for each experimental plasmid to control for transfection repeatability. Briefly, we made a master mix with 250uL Opti-MEM Reduced Serum Medium, 2.5ug total DNA, and 7.5uL *Trans*IT, allowing the *Trans*IT-LT1 Reagent:DNA complexes to form during an incubation, and we distributed this mixture in a dropwise manner to respective cells. We incubated cells for 24 hours, imaged and harvested, resuspending in DMEM growth medium (Gibco™, DMEM, high glucose, GlutaMAX™ Supplement, pyruvate). We confirmed transfection was successful by first checking control cells transformed with a GFP plasmid (Custom plasmid from vectorbuilding, EGFP:T2A:Puro driven by Zebrafish Beta actin). We moved ~80 µl of cells to a BRANDplates opaque, white tissue culture 96-well plate in triplicate, aiming for a total of ~4000 cells in each well.

For the luciferase assay, we used the Nano-Glo Dual-Luciferase Reporter Assay System (Promega) with the Firefly Luciferase with TK Promoter (pGL4.54 [luc2/TK] Vector). We prepared the Stop & Glo and One-Glo Reagents fresh, allowing them to thaw to room temperature while protected from light before the start of the luciferase assay. We configured the Agilent Cytation5 to read the Firefly luminescence of each well (integration time = 0.5s, gain = 150). We added the One-Glo Reagent to each well with a multichannel pipette and incubated for 3 minutes while orbitally shaking at 425 rpm. We then added the Stop & Glo Reagent with a multichannel pipette to each well, incubated while orbitally shaking, and measured NanoLuc luminescence with the same settings. We ran the entire set of luciferase experiments described above on three separate days to confirm that our results were robust to batch effects (Fig. S24).

To visualize these results, we first subtracted each luminescence reading from an averaged background reading (untransfected HEK 293T cells) and normalized the empty or experimental NanoLuc luciferase wells to firefly luciferase (pGL4.54) activity (NanoLuc/Firefly) to account for transfection efficiency within each well. We then averaged across 3 technical replicates for each sample (N=3 empty, N=3 LTR) and for other sequences of interest in the GWAS region including the *MG3f* sequence (N=3; pNL3.2 with *MG3f* insert) and sparkle and non-sparkle hypervariable regions (N=3 pNL3.2 with sparkle variable, N=3 pNL3.2 with non-sparkle variable). To analyze how different inserts impacted reporter gene expression while controlling for possible batch effects, we ran a one-way ANOVA on the data from each day we ran the luciferase experiment (3 total), followed by a Tukey HSD test. Across all experiments, we found a statistically significant increase in reporter gene activity in all inserts compared to the empty control (Fig. 3G, Fig. S24) except for the *MG3f* sequence (*MG3f* vs control: batch 1 – $t=0.413$, $p=0.993$; batch2 – $t=-3.89$, $p=0.020$; batch3 – $t=-0.273$, $p=0.999$; One-way ANOVA and Tukey HSD).

Our results clearly point to an important role of the LTR as a possible *cis*-regulatory sequence associated with the sparkle trait. The LTR sequence drove higher levels of reporter gene expression than any other tested plasmid (LTR vs control: batch 1– $t=21.70$, $p<0.001$; batch 2 – $t=16.13$, $p<0.001$; batch 3 – $t=21.89$, $p<0.0001$; based on One-way ANOVA and Tukey HSD). The *MG3f* insert did not drive expression *in vitro* relative to the control plasmid, while the sparkle variable region seemed to demonstrate weaker *cis*-regulatory potential than the LTR (Fig. 3G, Fig. S24). Intriguingly, the hypervariable region derived from non-sparkle individuals did have elevated activity in the reporter assay (non-sparkle variable region versus control: batch 1 – $t=14.43$, $p<0.001$; batch 2 – $t=12.55$, $p< 0.001$; batch 3 – $t=13.99$, $p<0.0001$; One-way ANOVA and Tukey HSD). However, the LTR sequence was consistently a stronger driver of luciferase activity than other tested sequences (Fig. 3; Fig. S24), including the hypervariable region derived from non-sparkle individuals (LTR versus non-sparkle variable region: batch 1 – $t= 7.270$, $p<0.001$; batch 2 – $t=3.59$, $p=0.03$; batch 3 – $t=7.91$, $p<0.0001$; One-way ANOVA and Tukey HSD). Moreover, the non-sparkle hypervariable region is not found in sparkle individuals and thus cannot be the regulatory element driving the trait. Nonetheless, this observation raises interesting questions about expression modulation that may be exciting to pursue in future studies.

**Supplementary Materials 16. Analysis of transcription factor binding motifs in LTR region**

The LTRs of ERVs often contain transcription factor binding domains that facilitate their success in invading the host genome. To evaluate evidence for transcription factor binding domains in ERV-F.1-Xeve-chr-15, we used the TFBSTools package in R with the JASPAR2022 database. We searched the LTR region for the 156 binding domains included in the SELEX database of JASPAR2022. The results of this analysis are available in Table S9.

Examination of predicted transcription factor binding motifs highlighted the presence of predicted *sox10* and *tfec* binding sites, among a number of other binding motifs that are potentially of interest for understanding the sparkle trait (Table S9). *Sox10* and *tfec* transcription factors are particularly interesting in the case of the sparkle phenotype, since they play key roles in cell fate determination of iridophore cells. Specifically, in zebrafish upregulation of *sox10* and *tfec* in the presence of *ltk* (a receptor tyrosine kinase) drives differentiation of neutral-crest derived pigment progenitor cells into iridophores. The JASPAR database includes position weight matrices for the conserved vertebrate binding sites of *sox10* and the human *tfec*, and we found evidence for multiple predicted binding sites for both transcription factors within the LTR region of ERV-F.1-Xeve-chr-15.

Given evidence of *sox10* and *tfec* binding sites in the LTR region, we next formally scanned for these binding sites both in this region and the larger region surrounding *alkal2a* and the ERV-foamy insertion. To do so, we downloaded the position weight matrix for each transcription factor from JASPAR, and used FIMO (*62*) to scan the genome of an individual that was homozygous for the sparkle haplotype and an individual that was homozygous for the non-sparkle haplotype for potential *sox10* and *tfec* binding sites. We visualized the distribution of these predicted binding sites relative to *alkal2a*, the location of the ERV-foamy insertion and its flanking LTRs associated with the sparkle haplotype, as well as SNPs that distinguish the sparkle and non-sparkle individuals in the hypervariable region. We also performed this analysis on haplotypes aligned using MAFFT (*35*) to identify any variation between individuals (Fig. 3).

Our results indicate that the LTRs that occur in the sparkle haplotype add a substantial number of potential *sox10* binding sites (Fig. 3H), as well as four potential *tfec* binding sites (Fig. 3H). While the region upstream of *alkal2a*, the hypervariable region downstream of the TE insertion, and the *alkal2a* gene also contain potential *sox10* and *tfec* binding sites, they did not differ in number or location between the sparkle and non-sparkle haplotypes (Fig. 3). More directly measuring differences in transcription factor binding activity between sparkle and non-sparkle individuals will be an exciting future direction when such experiments become possible in *Xiphophorus*.

**Supplementary Materials 17. Population genetic analysis of ERV-F.1-Xeve-chr-15 insertion and surrounding region**

*PSMC analysis*

To understand the population history of the *X. evelynae* at Juntas Chicas and better parameterize simulations exploring the ERV-F.1-Xeve-chr-15 insertion (see below), we performed demographic inference using Li & Durbin's implementation of the Pairwise Sequentially Markovian Coalescent (PSMC) model (*63*).

To generate variant calls, we mapped PacBio HiFi data to the Juntas Chicas *X. evelynae* sparkle reference genome with minimap2 (*3*) and used DeepVariant (*64*) with a PacBio model type (--model_type PACBIO) to call variants using a deep learning approach. Following variant calling, we used the BCFtools (*27*) consensus command to generate a fasta sequence with variants called for each individual, masking invariant sites that were not well covered (DP<10) and variant sites that did not pass DeepVariant quality score thresholds. We used a script to convert from fasta to fastq format, arbitrarily setting the quality of all bases to 20 (https://github.com/Schumerlab/Lab_shared_scripts/blob/master/fasta_to_fastq.pl). Next, we used the psmc utils command fq2psmcfa to convert the fastq file to a PSMC formatted fasta file. We used default parameters for time interval parameters, set the generation time to 2 generations a year, and set the mutation rate to $3.5 \times 10^{-9}$ based on estimates from closely related species (*65*, *66*). To infer where we lose resolution to reliably infer population size in the recent and distant past, we performed bootstrap resampling of the data by sampling 250 kb regions with replacement and re-running PSMC.

The above analyses were performed using variant calls generated from long read data, which rely on relatively recently developed computational pipelines, and generally do not report quality scores for invariant sites. To confirm that this approach was not impacting our results, we repeated the analysis using simulated short read data. We used wgsim (https://github.com/lh3/wgsim) to generate simulated 150 basepair paired-end Illumina data from a diploid version of an individual male (JUCH-6-V-24-S178-M-01) and mapped these reads to the reference genome. We performed variant calling with GATK and analyzed the resulting data with PSMC as described previously (*15*, *67*). Based on this analysis, we found that inferred population size changes in both long-read and short-read approaches were qualitatively similar.

For further analyses, we generated an estimate of the time-averaged population size for the Juntas Chicas *X. evelynae*. Based on bootstrap results described above, we determined the likely timepoints where we lose resolution to infer population size in the recent and distant past. We restricted our analysis to this time period (between 2,000 and 100,000 generations) and estimated the harmonic mean of population size over time. We used this value (57,421) as our estimate of historical population size.

*Calculation of population genetic statistics*

To evaluate how local haplotype diversity changes as we approach ERV-F.1-Xeve-chr-15, we generated alignments of long-read haplotypes with the insertion. We selected a 1.5 Mb region that centered on the ~17 kb ERV-F.1-Xeve-chr-15 insertion from each haplotype and extracted this sequence using fastahack. We next used MAFFT to align these regions (*42*). Using Geneious, we visually inspected these alignments and generated alignment summary statistics, identifying no artifacts caused by poor alignments. We next applied a script to extract variable sites and their locations in the alignment (https://github.com/AdmiralenOla/PuppetMaster/tree/master). We summarized the locations of variable sites within sparkle and non-sparkle haplotypes as a function of distance to ERV-F.1-Xeve-chr-15 in sliding 5 kb windows (Fig. 4). We also calculated average pairwise sequence divergence between sparkle and non-sparkle haplotypes in sliding windows based on this alignment relative to the location of the ERV-F.1-Xeve-chr-15 (Fig. S20).

We also generated global estimates of $\theta_\pi$ for the *X. evelynae* in the Juntas Chicas population. Because we wanted to accurately estimate the denominator, we needed to use a variant caller that would also output information at invariant sites. We used the vcf generated for PSMC analysis by GATK described above. We counted all heterozygous sites that passed our quality thresholds (*68*) and divided this by the number of invariant sites that passed our quality thresholds. This resulted in an estimated $\theta_\pi$ of 0.09%, similar to values previously observed for other swordtail species. This value provides a useful comparison to observed population genetic statistics in windows flanking ERV-F.1-Xeve-chr-15 (Fig. 4; Fig. S20).

*Extended haplotype homozygosity and Tajima's D analysis*

We were also interested in whether we could detect evidence of selection using population genetic approaches. This is complicated by the nature of this insertion, since the region that may be under selection has no homologous sequence in the non-sparkle individuals. We thus focused on regions directly flanking the TE insertion for these analyses. We selected 1 Mb of sequence in the 5' and 3' direction of ERV-F.1-Xeve-chr-15. We used blastn to extract the coordinates of these regions from each of the *de novo* assemblies from the 11 phased haplotypes for chromosome 15. We used MAFFT to align these regions and visually inspected the alignments in Geneious to ensure that they were free of artifacts. Next, we converted the alignments into .hap format for calculation of the Extended Haplotype Homozygosity (EHH) statistic using the R package rehh (*69*). We focused on several SNPs in the hypervariable region downstream of ERV-F.1-Xeve-chr-15 and examined patterns of EHH with these SNPs (Fig. S20).

It is possible that the sparkle phenotype is not under directional selection but instead experiencing balancing selection in the population. To investigate this possibility, we used the same alignment and the program tajimas_d (https://github.com/not-a-feature/tajimas_d) to calculate Tajima's D in sliding windows. We excluded windows with high levels of gaps from this analysis (>20% gaps). We found evidence of positive Tajima's D on the edges of the alignment flanking ERV-F.1-Xeve-chr-15, potentially consistent with balancing selection, but this signature was localized to an approximately 30 kb region on the 5' and 3' edges of the insertion (Fig. S20).

### Supplementary Materials 18. Simulations to study the evolution of ERV-F.1-Xeve-chr-15

*Simulations to infer the likely age of the ERV-F.1-Xeve-chr-15 insertion*

Comparisons of sequence similarity at ERV-F.1-Xeve-chr-15 within and across individuals suggested that it likely inserted into the genome in the recent past (Fig. 4A). We identified only three nucleotide differences (two SNPs and one INDEL) in comparisons across the four assembled sparkle haplotypes containing the ERV-F.1-Xeve-chr-15 insertion, with an alignment length of 16.9 kb. The average pairwise haplotype divergence was 0.009%. We were curious whether we could infer the likely time since the element was incorporated in the genome using a grid-search approach and simulations in SLiM 4 (*70*).

To perform simulations, we designed a scenario to model the recent insertion of an ERV element. We used a mutation rate of $3.5 \times 10^{-9}$ based on empirically measured mutation rates in close relatives of *Xiphophorus* (*65*, *66*). We set the population size to 57,421 matching the time-averaged population size for *X. evelynae* from Juntas Chicas inferred from PSMC. We modeled a region 16.9 kb in length and assumed that a single copy was inserted at some time in the past. For each time in our grid, ranging from 5000 generations to 35,000 generations, we performed 100 replicate simulations. At generation 1, we introduced a mutation that we treated as a "tag" of the ERV-F.1-Xeve-chr-15 insertion. We used code from the SLiM recipe resource to check at generation 1,000 and each subsequent generation whether the simulated TE insertion had been lost. If it was lost, we restarted the simulation. Once the simulation reached the end of the specified number of generations, we selected only individuals with the tag mutation and used the outputVCF function to report their genotypes along the 16.9 kb region. We calculated pairwise sequence divergence between two randomly sampled simulated haplotypes and recorded this value as our summary statistic. For each value in our grid, we repeated this process until we had results from 100 simulations and calculated the average pairwise sequence divergence. We compared this value to that observed in our real data and repeated this for all entries in the grid (5,000-35,000 generations).

Because meiotic recombination is likely to be locally reduced around insertions and may be particularly reduced near insertions involving active ERVs, we performed these simulations with multiple recombination rates. The simulations described above modeled the average genome-wide recombination rate inferred for another *Xiphophorus* species, *X. birchmanni* (*68*). We next modeled a recombination rate $10\times$ lower than the genome-wide average, and found our results were qualitatively unchanged. In practice, we expect that the empirical recombination rate in this region will change over time as the insertion increases in frequency and is more likely to be found in the homozygous state. However, the insensitivity of our simulations to the exact recombination rate used suggests that our results will be robust to this parameter.

Initial simulation results indicated that the insertion likely occurred between 10 and 20 thousand generations before the present. We thus repeated the procedure described above for every 1,000 generations between 10-20 thousand generations. While a range of parameter sets are consistent with the results we observe, the similarity between the observed and simulated data was maximized at a simulated insertion age of 16,000 generations (Fig. 4; average difference between number of simulated and observed variants: -0.007). This indicates that the insertion occurred relatively recently, consistent with the conclusion that ERV-F.1-Xeve-chr-15 is active or was recently active in *X. evelynae*.

*Simulations incorporating more realistic population history*

In the above simulations, we used the time-averaged population size and did not model changes in population size. However, there is clear evidence of a relatively recent bottleneck in our PSMC analysis. To evaluate expectations for insertion age in this scenario, we modeled the inferred population history of the Juntas Chicas *X. evelynae* using two step functions to roughly approximate historical changes in population size (Fig. S42). We repeated the simulations described above with this more realistic demographic history. We found that simulation results were qualitatively similar but consistent with a slightly older insertion time, with simulations of an insertion 20,000 generations before the present most closely matching the observed diversity in the insertion haplotype on chromosome 15 (average difference between simulated and observed data: 0.064).

*Simulations to evaluate expected frequency of ERV-F.1-Xeve-chr-15 under neutrality*

Given our results regarding the likely age of the ERV-F.1-Xeve-chr-15 insertion, we were interested in examining its expected frequency under a scenario of neutral evolution. To do so, we implemented simple simulations in SLiM 4 (*70*). As a starting point for these simulations, we used the time-averaged Juntas Chicas population size of 57,421 from our PSMC results and the inferred insertion time of 16,000 generations before the present from our grid search results. We assumed that insertion occurred a single time, so we set the starting frequency to 1 out of 2Ne and performed simulations as described above. Given that we observe the ERV insertion in the present-day population, we restarted any simulations where the simulated insertion was lost. For all simulations where the insertion established, we continued the simulation until we reached 16,000 generations and recorded its frequency at the end of the simulation. We repeated this procedure until we had recorded frequencies from 200 simulations and compared the distribution of simulated frequencies to the observed frequency.

Conditional on the simulated insertion being preserved, we found that a broad range of frequencies were observed after 16,000 generations. Assuming that the sparkle haplotype is dominant (as indicated by our empirical data), simulated trait frequencies ranged from 0.0003 to 0.63 (95% confidence intervals: 0.007-0.36). While the observed frequency of the trait trends towards being significantly higher than expected, these simulations suggest that we cannot confidently exclude genetic drift as a cause of the high frequency of the sparkle trait in the Juntas Chicas population (p-value = 0.07 by simulation).

**Supplementary Materials 19. Maximum reflectance angle and brightness analysis**

*Reflectance measurements*

To measure reflectance of individuals with different phenotypes, we used 46 sexually mature *X. evelynae* fish from Juntas Chicas, of which 22 were males and 21 females, and 3 juvenile fish. These individuals were divided into two groups: fish with apparent iridescence (SP) and fish without iridescence (non-SP). The group of fish with sparkle included 12 females, 17 males, and 3 juveniles, while the fish without iridescence included 9 females, 5 males, and no juveniles. Fish were randomly selected from a stock population established at the Centro de Investigaciones Científicas de las Huastecas Aguazarca (CICHAZ) facilities in November 2023. The fish were maintained at the CICHAZ outdoor facilities in four 400 L mesocosm tanks and were fed twice daily.

Fish were anesthetized by immersion in water with MS-222 at a dose of 100 mg/mL. This same solution was placed in the anodized tub used to measure fish reflectance (see below) to ensure they remained sedated and in a humid environment during measurements. Once reflectance was measured, fish were placed in a recovery tank. After the experiment, each fish was returned to its home tank.

All reflectance measurements were made in a darkroom. The experimental setup consisted of an optical sensor (a 400 µm diameter optical fiber that records reflectance) attached to a clamp on a goniometer at a height that varied depending on the anatomy of each fish (the height was adjusted so that the light from the optical fiber focused on and illuminated the focal scale). The goniometer was moved until the optical fiber was adjusted to the desired angle (15°, 30°, 45°, 60°, or 90°). The optical fiber used was bifurcated: one end was connected to the light source (a DT-MINI-2 GS deuterium halogen lamp) and the other to a spectroradiometer (HR4000) with a spectral range of 300 to 800 nm. The spectroradiometer sent the signal to a computer (HP 4dq2531la), where the reflectance measurements were recorded using OceanView 2.0.8 software. Once anesthetized, the fish was placed in an anodized tub while reflectance measurements were taken at all desired angles. A Spectralon Ocean Optics white standard (WS-1-SL) was used as a reference for reflectance measurements.

*Brightness analysis*

The brightness of the scales was measured when scales reached their maximum reflectance from our above measurements (following *Bird Coloration, Volume 1*, 2006). To do this, the area under the reflectance distribution curve was measured using the equation:

$$Brightness = \int_{\lambda min}^{\lambda max} Ri = \sum_{\lambda min}^{\lambda max} Ri$$

where $\lambda_{max}$ is the maximum reflectance and $\lambda_{min}$ is the minimum reflectance for each wavelength *Ri*, from 300 to 800 nm (Fig. S43). This procedure was repeated for each of the angles considered in this study. To analyze these results, we fit a linear model to our brightness data. The model used was lm(brightness ~ as.factor(sparkle)*angle + sex. We found the brightness of the sparkle trait differed significantly from the non-sparkle trait effect at an angle of 90° (p-value = 0.0297), but not at other angles.

**Supplementary Materials 20. Behavior trials and analyses**

*Scototaxis Behavior trials*

Nineteen wild-caught *X. evelynae* (8 sparkle, 11 non-sparkle) were tested in a scototaxis assay to investigate whether the having the sparkle trait correlated with differences in boldness behavior. These fish were acclimated to the lab for several months before trials began. Scototaxis trial lanes (50cm L x 19cm W x 19cm H) were half lined with white panels and half lined with black panels at the length midline. Fish were placed into covered, black habituation buckets with a shelter for 10 minutes before trials. Each fish was gently placed into the dark side of the trial tank. The experimenter was visually occluded, and fish movement was recorded for 15 minutes using the EthoVisionXT16 software and an overhead camera. Side bias was tested by re-testing each fish twice in a tank with a flipped black and white scheme and averaging data across the two trials. After each trial, a water change was performed on both the trial tanks and habituation buckets.

Data were analyzed in EthoVision. We used the automated tracking function with manual correction of any errors in EthoVision's tracking of individuals. Total time spent in the white area and black area, latency to cross into the white zone, and the number of times a fish crossed between the two areas were determined for each video. We analyzed whether there were differences in boldness behaviors between phenotypes using a t-test. We found no significant differences in the average time to cross from the dark zone into the light zone ($t = -0.065$; $p = 0.95$), the number of crosses between zones ($t = 0.38$; $p = 0.71$), or the total time in the light zone ($t = -0.37$; $p = 0.72$). This suggests that the sparkle trait is not linked to differences in boldness in *X. evelynae*, but we note that we have limited power to detect slight differences given our sample size.

*Female preference trials with animations*

To test whether *X. evelynae* females exhibited preferences for mates with or without the sparkle trait, we conducted a dichotomous mate choice trial using twenty wild-caught females that had habituated to the lab environment over several months. In this assay, we placed a single, sparkle or non-sparkle female fish in a 75x19x20 cm trial tank containing 14 L of water and allowed her to habituate in the tank for 10 minutes. The trial tank contained a small, black shelter in the middle (neutral) zone of the tank. On either side of the tank, we introduced a courting male fish animation. To create these animations, we imaged a *X. evelynae* male with secondary sexual characteristics (tail spot, dorsal fin) common of courting males and a male with the sparkle trait. We used Adobe Photoshop to overlay the sparkle trait onto the first fish, creating a custom set of animations where females are exposed to two males that are identical in appearance and behavior apart from the presence of the sparkle trait. We projected these pairs of animations onto flanking monitors positioned on each side of the tank, with one side showing a sparkle male and the other side a non-sparkle male. After the acclimation period, female fish were shown the male courtship animations for 5 minutes, allowed 5 minutes with no stimulus between trials, and then exposed to swapped animations to test for side-bias and tested female preference for an additional 5 minutes. Water was changed following each trial, and females were anesthetized and photographed at the end of the trial.

Using BORIS v.7.13.9, a free open-source software to analyze behavior videos, we recorded female movement across three predetermined zones of their tank. These zones were defined as one neutral (middle) zone and two preference zones (25 cm from each video screen). We created two binary codes, "non-sparkle" or "sparkle", in which the trial time spent in the two preference zones could be summed from each female. We removed trials in which the female fish spent >90% of her time in the same preference zone, regardless of which animation was projected. Since animations were swapped between the two trials, we interpreted this behavior as evidence of side-bias. For the remaining trials, we averaged the time spent in the sparkle or non-sparkle preference zone across the two, 5-minute trial periods for each trialed female that did not exhibit side bias (n= 8 sparkle, n=8 non-sparkle females). Then, we calculated a preference index from 1.0 to -1.0, where 1.0 means a female spends 100% of time in the male Sparkle preference zone, and -1.0 means a female spends 100% of time in the male non-sparkle preference zone. We calculated this by taking the difference of the average time spent with Sparkle fish from the average time spent with non-sparkle fish and dividing by the sum of average time spent with Sparkle and Non-sparkle fish. We then ran a two-sided Wilcoxon signed rank test on the Non-sparkle and the Sparkle preference indices, with the null hypothesis that there is no preference (index = 0.0), finding no significant deviations from no preference in either female Non-sparkle (p-value = 0.38) or Sparkle (p-value = 1) phenotypes (see main text; Fig. 5).

*Response to perceived predator trials*

We designed a custom behavioral trial to investigate potential differences between sparkle and non-sparkle individuals in their responses to perceived predators. The dorsal side of a kingfisher model bird (BirdMobile Card Sculpture Kingfisher, Hobbyprof) was attached to a 1 mL plastic pipette tip with glue. A fishing line was threaded through the pipette tip and attached to two points above the tank at a downward slope, starting at a height of 67 inches and ending at 42 inches. The fishing line was attached to the ceiling above the trial tank. Individual *X. evelynae* fish were placed into a tank (dimensions = 18 x 18cm) filled with 4 inches of water and covered with black panels on all sides to conceal fish from early detection of avian predator. Fish acclimated in the trial tank for 5 minutes. Two soft box photography lights were mounted above the trial tank to increase lighting from all angles in order to best capture reflected light from the fish. All tested fish were food-deprived on the morning before trials. After the habituation period, the avian replica was dropped over the tank to elicit a predator-induced response from the fish (Video S3). The startle reaction was classified as a freezing, diving, or C-start response. These behaviors were recorded with an overhead mounted GoPro Hero 11 Black (5.3K, standard view, 60fps, narrow lens, auto white balance, natural color). At 10 seconds after the start of the recording period, the avian model was dropped from its peak position, appearing to "dive" over the tank (Video S3), and the fish was recorded for an additional 20 seconds.

Trial videos were downloaded, and each frame was inspected for a "flash" response visible to the overhead camera and avian viewpoint. We trialed both wild-caught fish that were habituated to the lab for X months, and lab-born fish that were born in the roof tank and brought down as juvenile fish in the main lab. We excluded trials in which the fish initiated a startle response before the avian model was released and trials where the fish did not initiate a startle response (no freezing, diving, or C-start response).

### Supplementary Figures

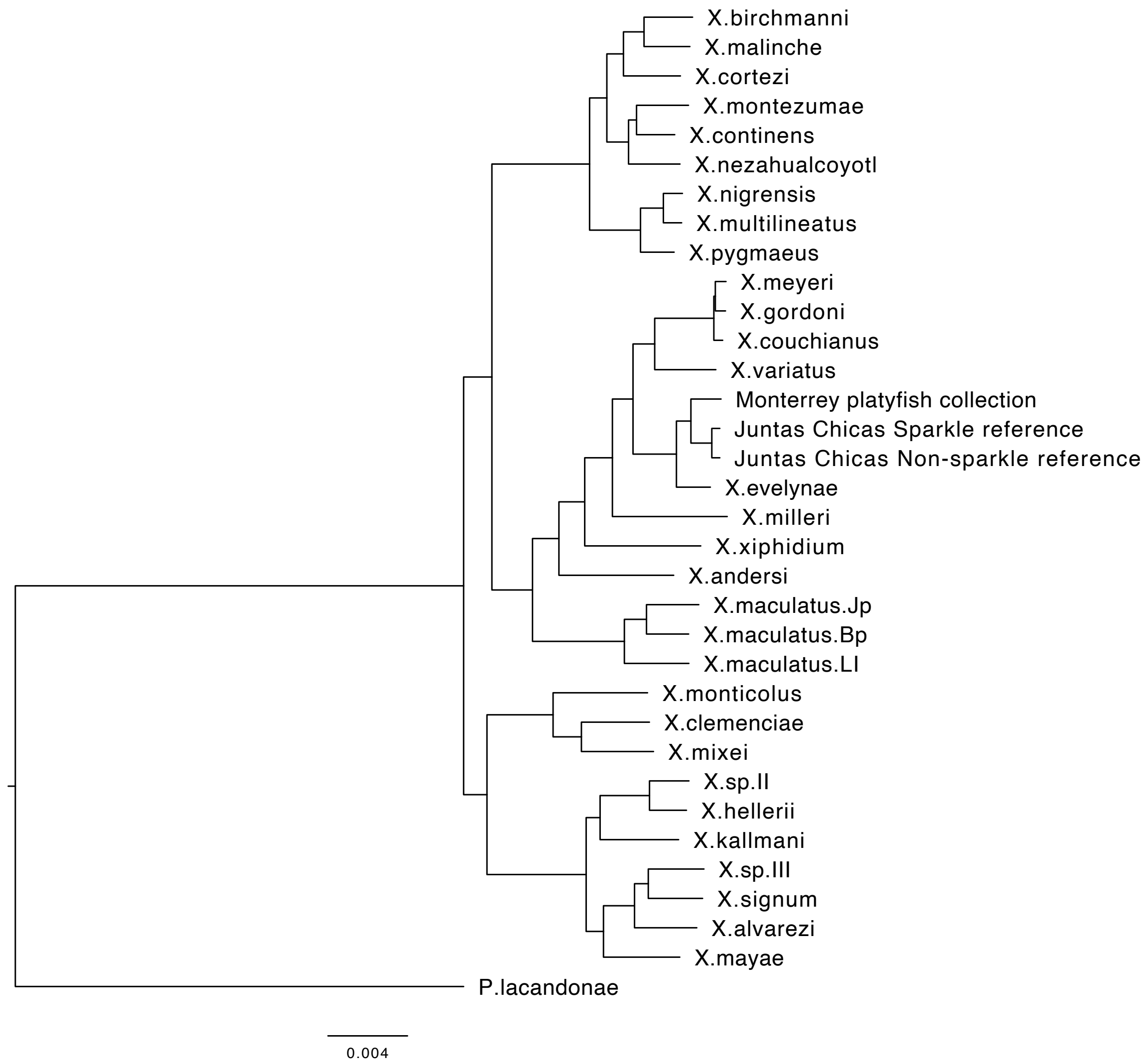


**Fig. S1.** Results of phylogenetic analysis of Juntas Chicas sample along with other previously sequenced *Xiphophorus* species. These results indicate the fish sampled from Juntas Chicas are closely related to the *X. evelynae* lineage. We note that the genetic divergence between the Juntas Chicas sample and described species may indicate that it is a distinct lineage, but for the purposes of this manuscript we treat it as a sub-species of *X. evelynae*. All nodes had 100% bootstrap support based on RAxML analysis, so they are not listed in the figure.

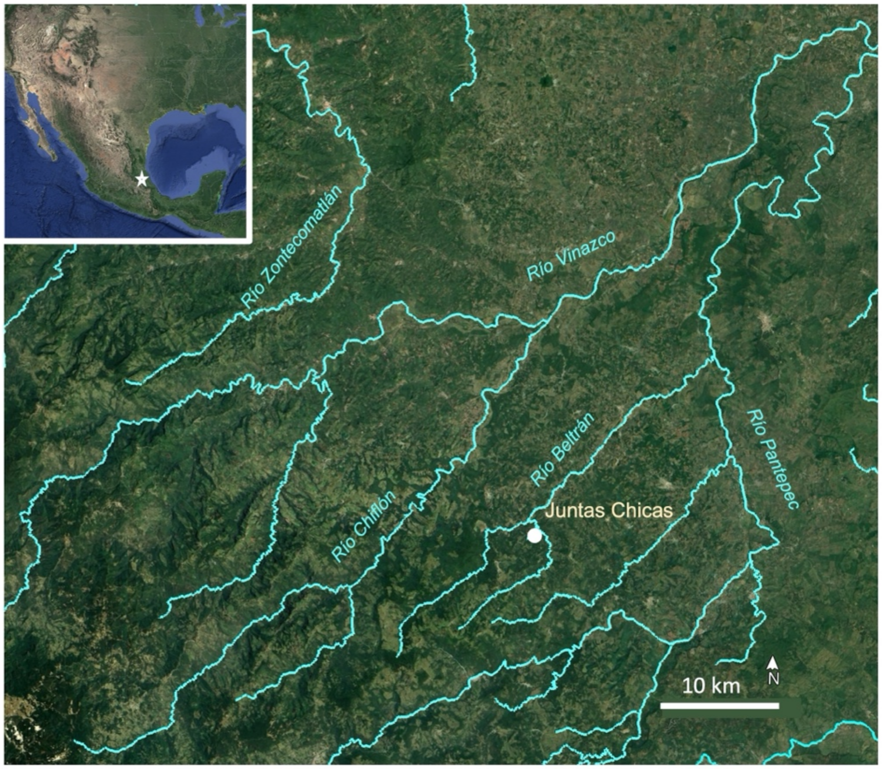

**Fig. S2.** Map of sampling location of Juntas Chicas population in Veracruz Mexico. All GWAS individuals were wild-collected fish from this site. Most fish used in laboratory experiments were lab-raised offspring of wild-caught fish.

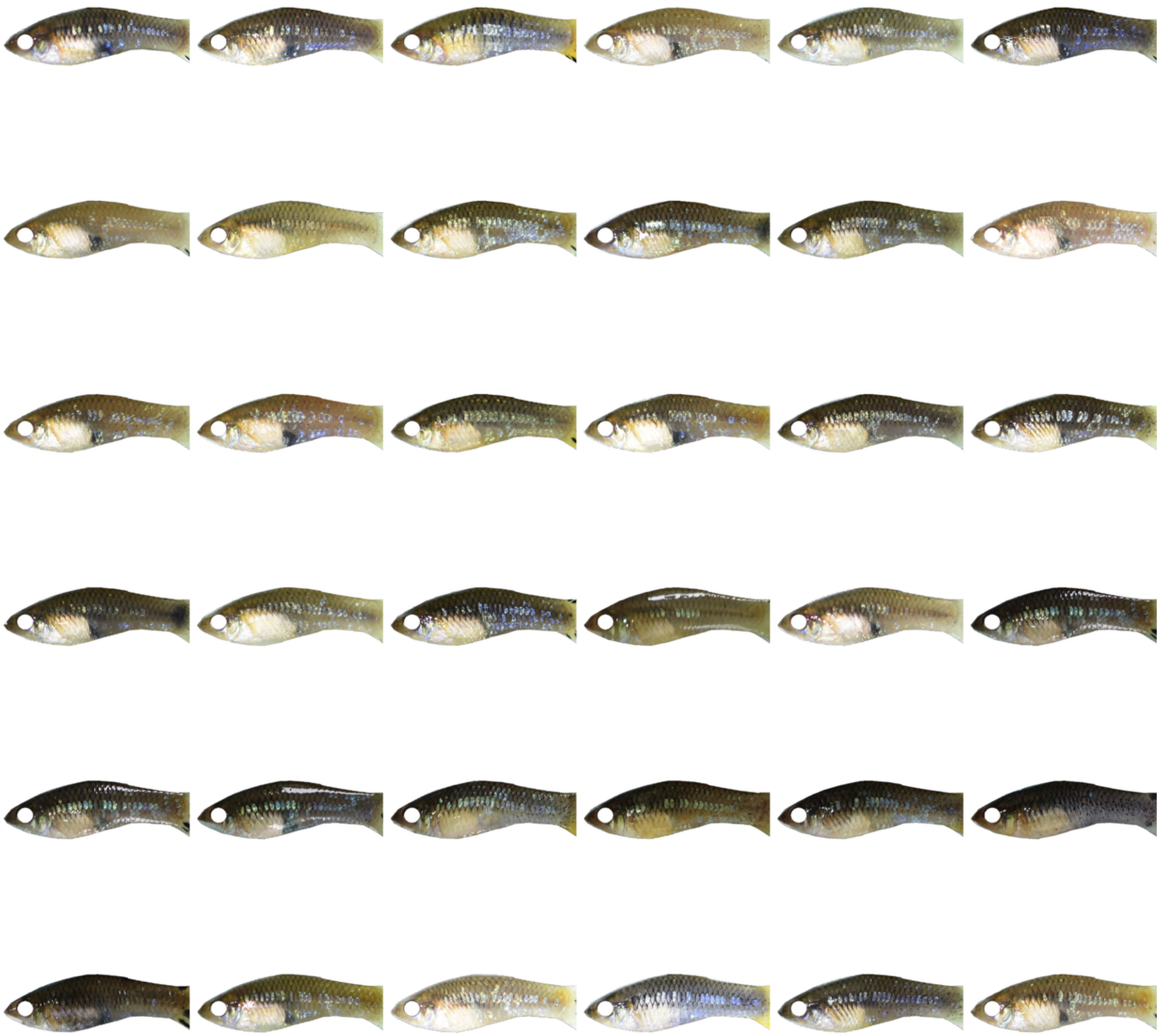

**Fig. S3.** Example of variation in adult sparkle phenotype in wild-caught *X. evelynae* included in our GWAS. These photos highlight variability in localization and intensity of the iridophore phenotypes. Masks of eyes and caudal fin added by an experimenter were used for automated quantification.

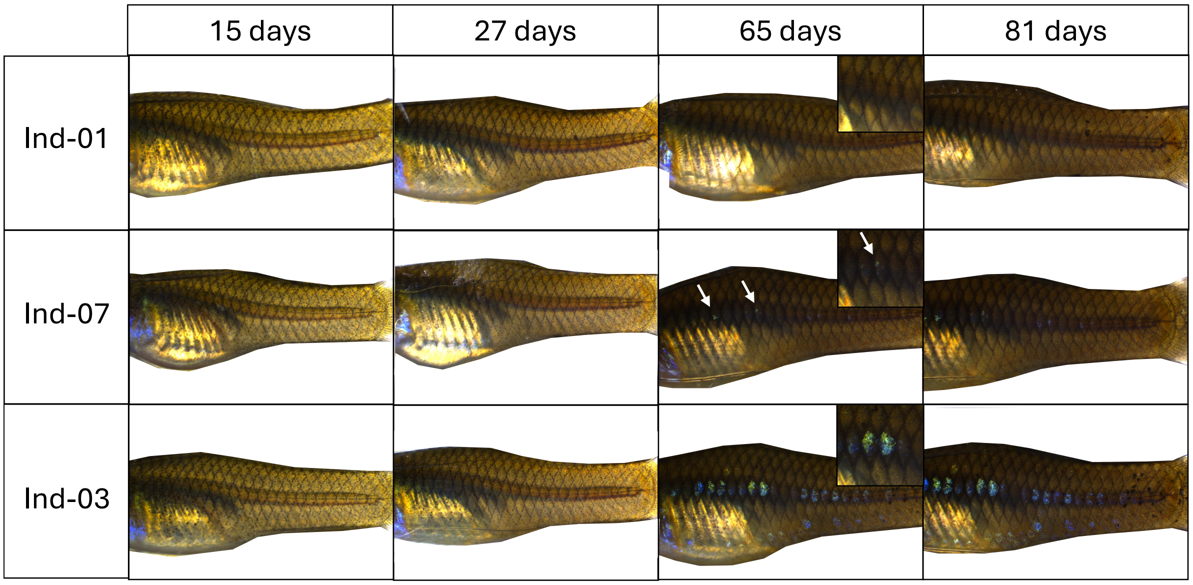


**Fig. S4.** Example of variation in developmental onset of the sparkle phenotype from three representative individuals of age-matched *X. evelynae* derived from a single brood. Images were taken under the K7 microscope before and after individuals develop sparkle. Individual-01 never developed sparkle (non-sparkle), individual-07 developed a small amount of the sparkle trait, and individual-03 demonstrated high expression of the sparkle trait.

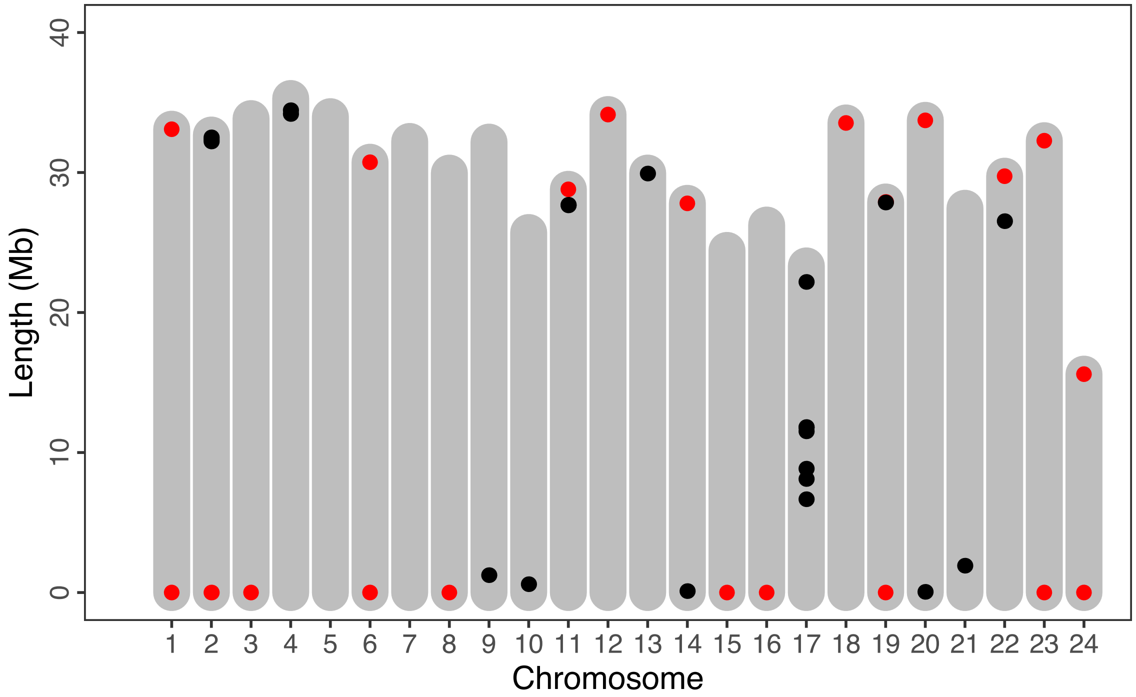


**Fig. S5.** Illustration of all gaps (black) and detected telomeric sequences (red) in the Juntas Chicas sparkle *X. evelynae* reference assembly, plotted by chromosome. The Y-chromosome is excluded from this analysis. Chromosome 21 is the X chromosome in most *Xiphophorus* species.

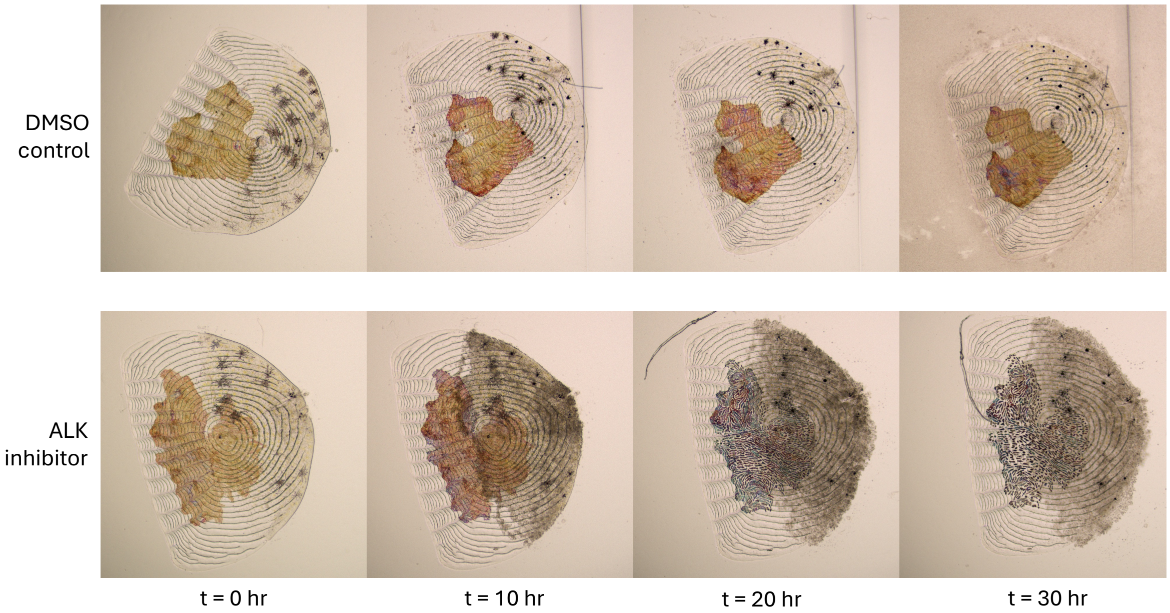


**Fig. S6.** Additional images of scales treated with an ALK inhibitor and control scales. Individual scales were plucked from the midline of sparkle fish and either submerged in a 2% DMSO or 2% ALK inhibitor solution (dissolved in DMSO) in cell culture media. Images were taken on the Agilent Cytation 5 at 0, 10, 20, and 30 hours post treatment. Iridophore cell populations are visible as orange/red cell clusters on the scale. Only the ALK inhibitor treatment group display loss of iridophore cell morphology, potentially indicating cell death.

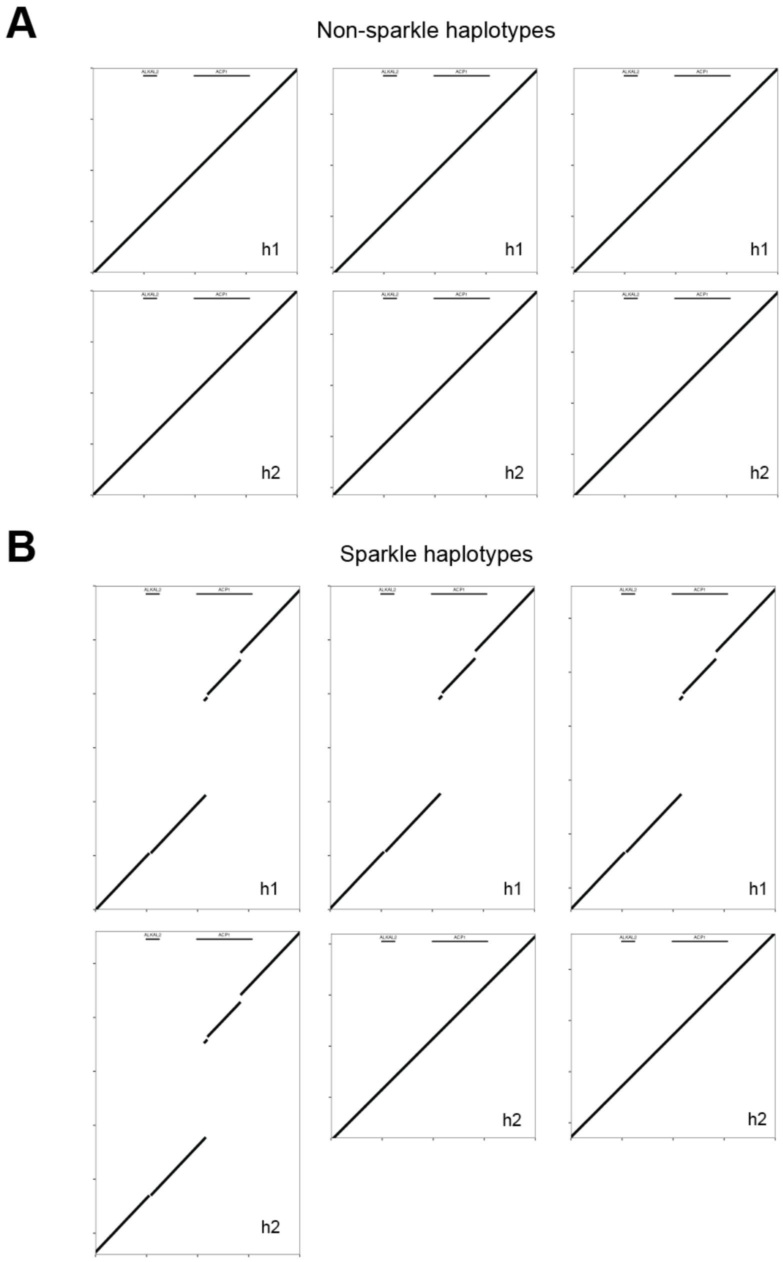


**Fig. S7.** Sequence alignments generated using MUMmer4 in which each haplotype assembled from long-read data (Y-axis) was aligned to the non-sparkle *X. evelynae* reference genome (X-axis). **A**) Non-sparkle haplotypes are syntenic across the 11,070,000–11,110,000 bp region, when aligned to the non-sparkle reference assembly. **B**) Haplotypes from sparkle individuals aligned to the non-sparkle reference assembly. At least one of two haplotypes derived from each sparkle individual contain a large, ~17 kb insertion relative to the non-sparkle reference genome. h1 – haplotype 1, h2 – haplotype 2. The inset lines on the top of each plot indicate the locations of *alkal2a* and *acp1* in the non-sparkle reference assembly.

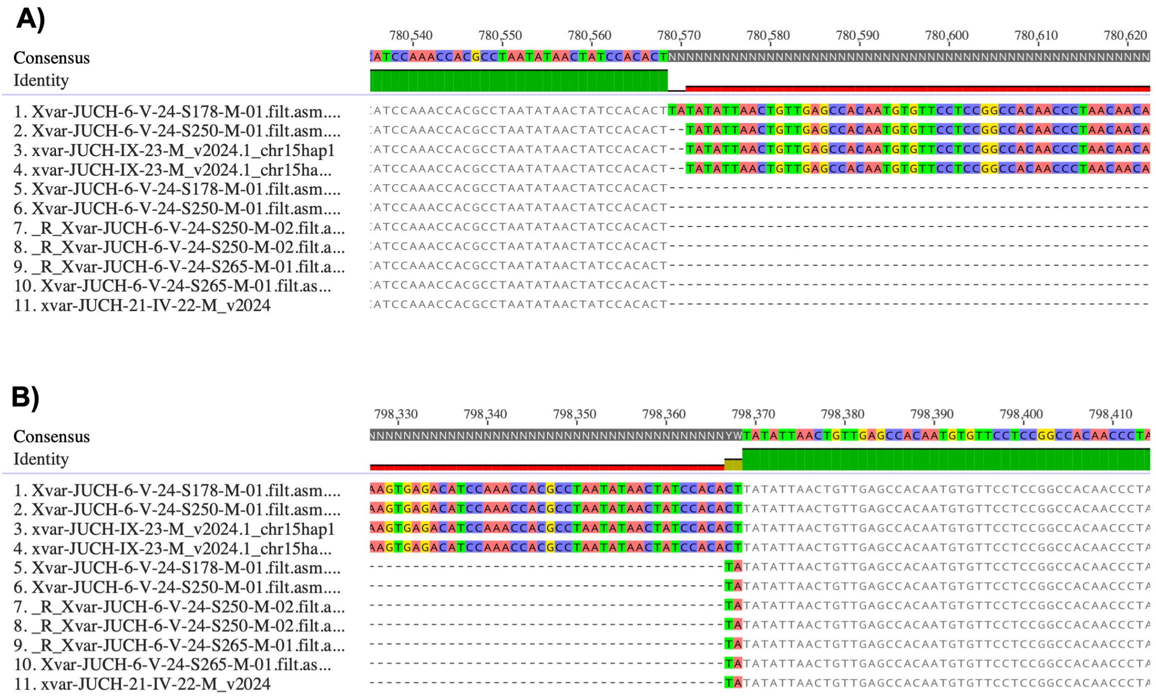


**Fig. S8.** Local haplotype alignment generated using MAFFT, illustrating the boundaries of the ERV-F.1-Xeve-chr-15 insertion. This alignment of the 1.5 Mb region surrounding ERV-F.1-Xeve-chr-15 was used to analyze insertion sites and for downstream population genetic analysis. **A)** Alignment highlighting the 5' edge of the ERV insertion on chromosome 15. Individuals are sorted such that the four haplotypes with ERV-F.1-Xeve-chr-15 insertions are first, followed by haplotypes that lack this insertion. **B)** Alignment highlighting the 3' edge of ERV-F.1-Xeve-chr-15. Alignments were visualized in Geneious.

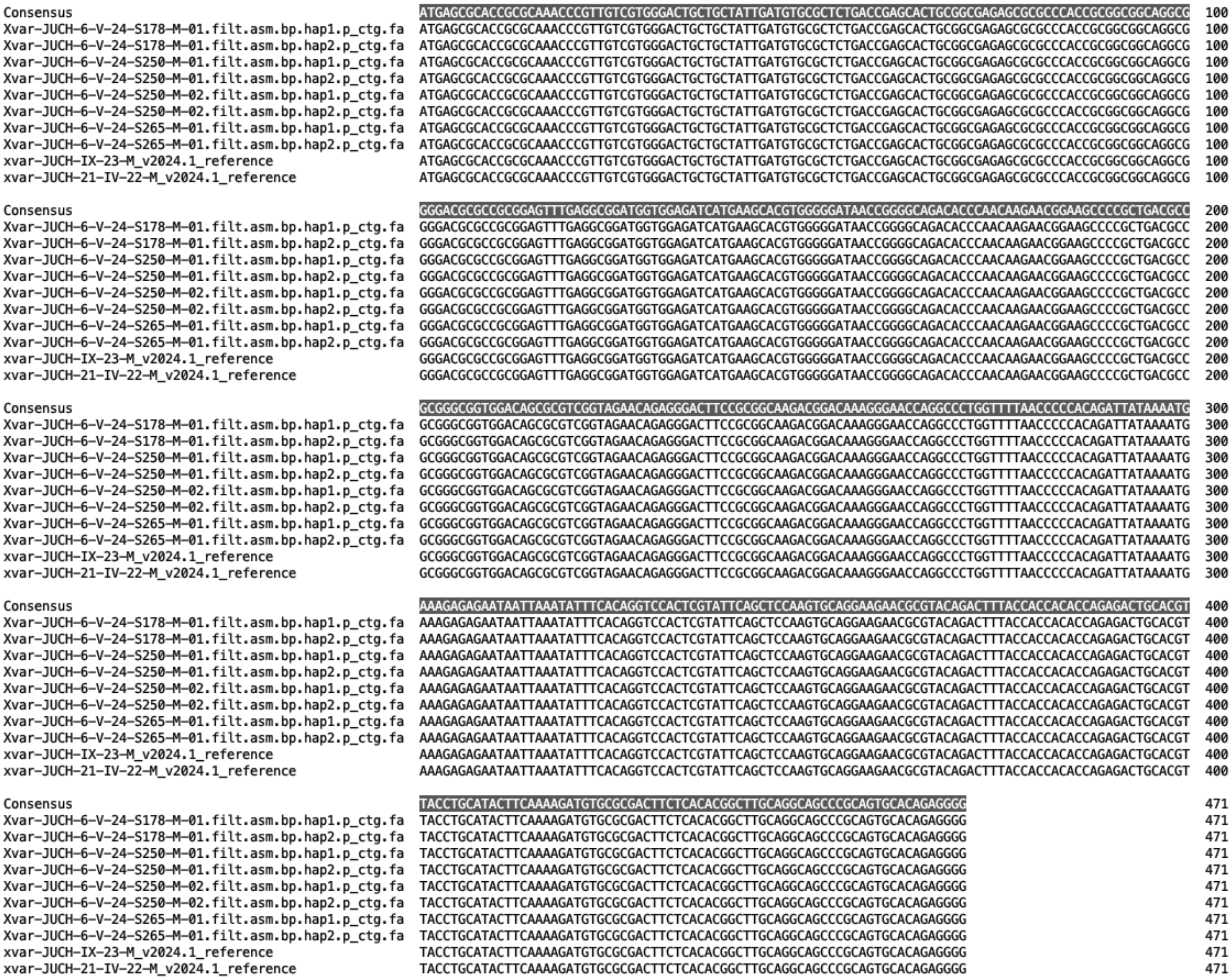


**Fig. S9.** Alignment of predicted *alkal2a* cDNA sequence across assembled *X. evelynae* haplotypes. We observe no variation at synonymous or nonsynonymous sites across *X. evelynae* haplotypes, consistent with high conservation of the *alkal2a* protein coding sequence across species (see Fig. S28).

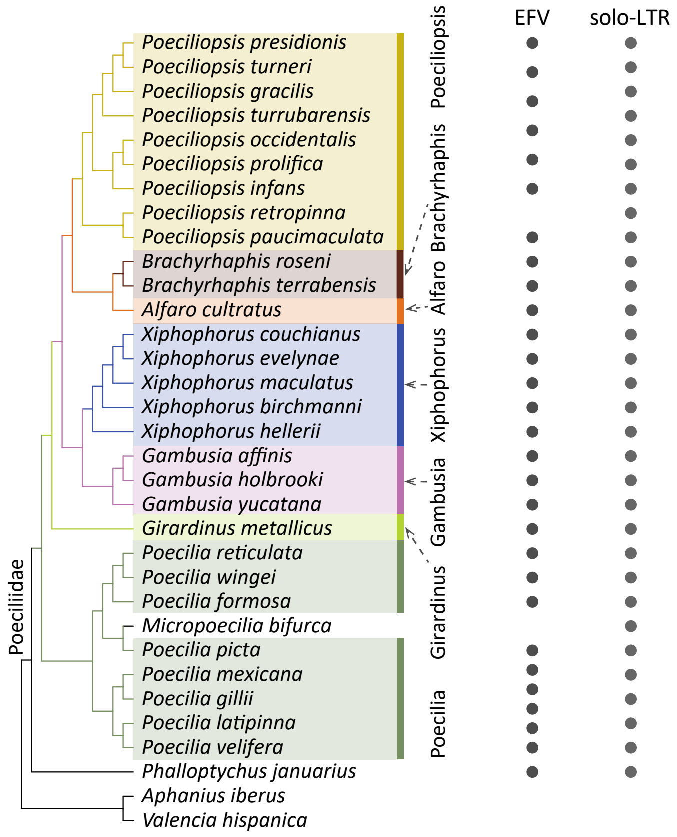


**Fig. S10.** The distribution of endogenous foamy viruses (EFVs), a genus of ERVs, and their solo-LTR derivatives in the genomes of Poeciliidae species. The presence of EFVs and solo-LTRs is indicated by filled circles. The colors for each species match the EFVs derived from these species, shown in Fig. S14.

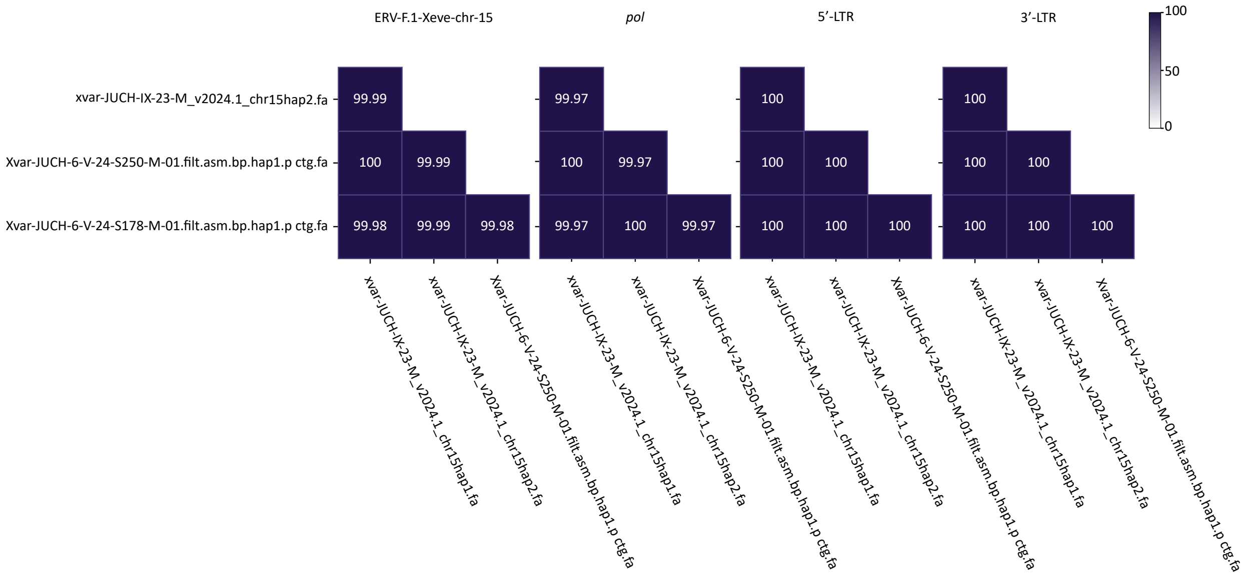


**Fig. S11.** Comparison of sequence divergence across sampled individuals at ERV-F.1-Xeve-chr-15. Analysis shows pairwise sequence identity for each haplotype with the ERV-F.1-Xeve-chr-15 insertion including the entire element, the *pol* gene, and the 5' and 3' LTRs. Notably, the 5' and 3' LTRs of ERV-F.1-Xeve-chr-15 are completely identical across individuals in the sampled haplotypes.

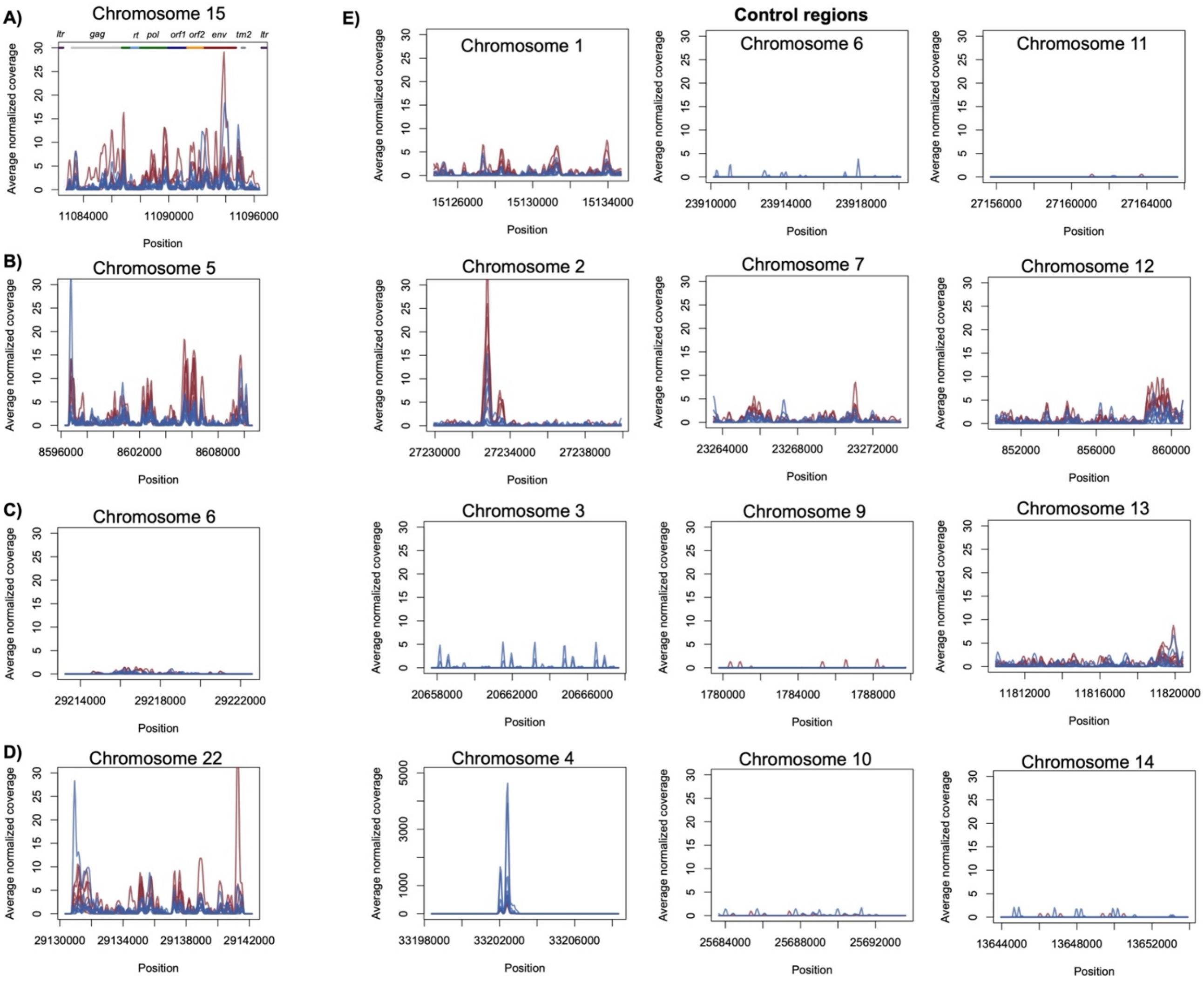


**Fig. S12.** Results of expression analysis for ERV-foamy-Xeve transcripts in brain tissue (red) and testis tissue (blue) compared to control regions. All samples are normalized such that read counts per 10 million mapped reads are plotted. **A-D)** Subset of ERV-foamy-Xeve regions analyzed for evidence of expression in the brain and testis across eight *X. evelynae* individuals. **A**) RNA-seq reads mapped to ERV-F.1-Xeve-chr-15 indicate some level of ERV-foamy-Xeve element expression in both the somatic and germline tissues, despite evidence of methylation of ERV-foamy-Xeve elements in somatic tissue. Line annotations above expression results plotted in **A** show different domains of the ERV-F.1-Xeve-chr-15 region. See Supplementary Materials 12 for details. **B-D)** Other full-length ERV-foamy-Xeve sequences annotated in the sparkle *X. evelynae* reference genome show varying evidence of expression in our analyses. A subset of these regions are shown here. See Supplementary Materials 12 for details. **E)** RNA-seq coverage was also analyzed in control regions that were selected as being size matched noncoding regions that did not contain annotated transposable elements. While some reads map in these regions, it is generally at low levels. Shown here is a representative subset of control regions from the first 14 chromosomes.

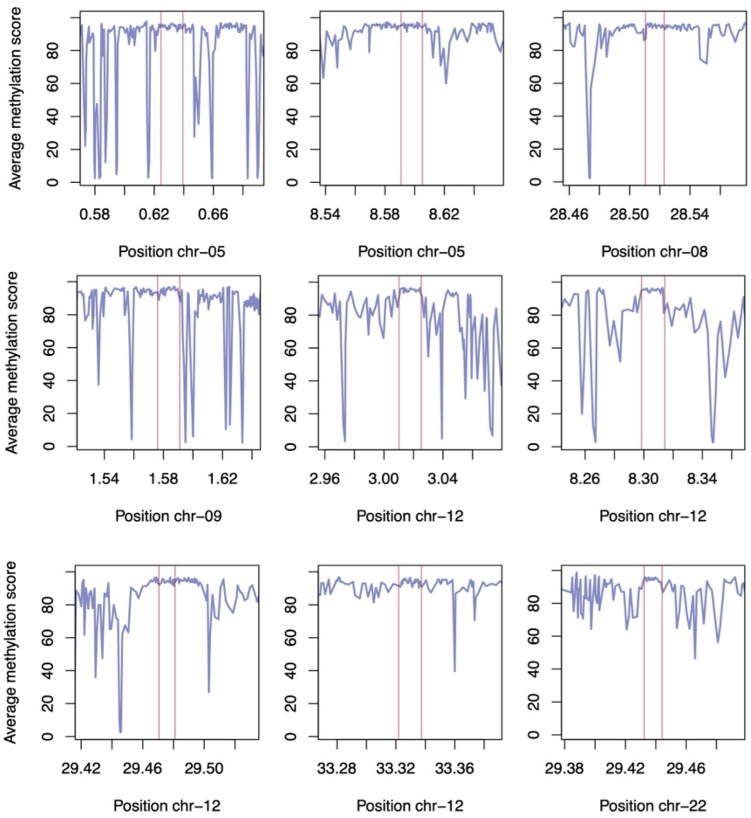


**Fig. S13.** Average discretized methylation score in windows of 10 CpG sites. Plotted data indicate methylation patterns nearby all full-length ERV-foamy-Xeve insertions in long-read data from a single individual, xeve-JUCH-21-IV-22-M, mapped to its own assembly. The red lines show the boundaries of each of the ERV-foamy-Xeve insertions in this individual. Similar patterns are observed for other complete ERV-foamy-Xeve insertions annotated across sequenced individuals.

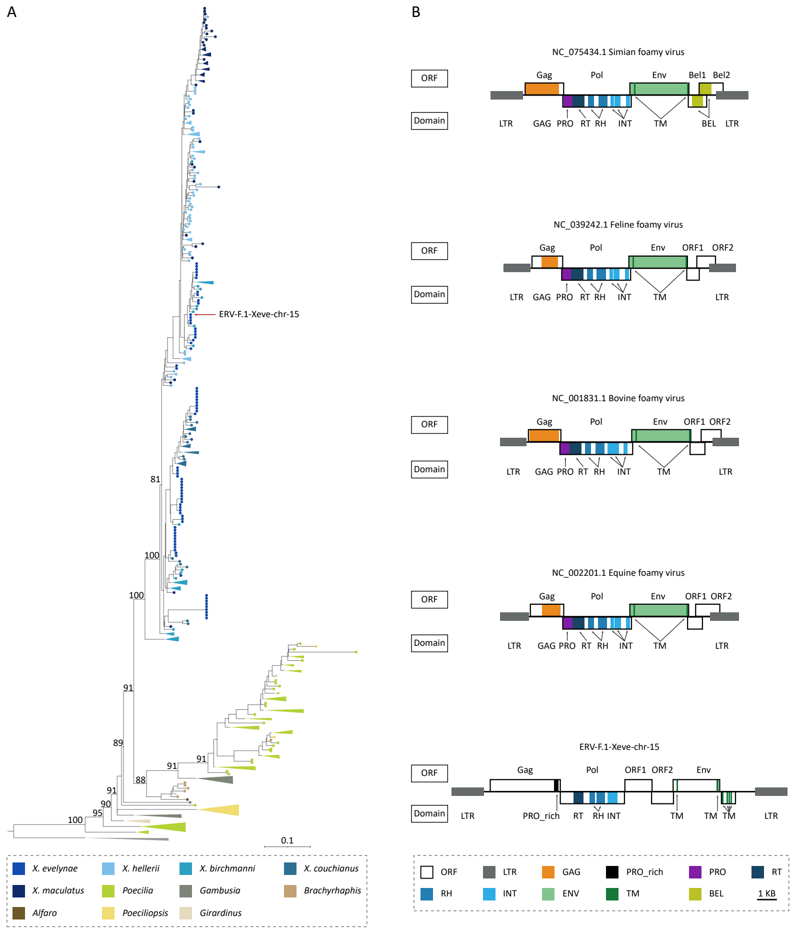


**Fig. S14.** Evidence of phylogenetic mixing in endogenous foamy viruses (EFVs), a genus of ERVs, detected within *Xiphophorus* genomes. **A)** The phylogenetic tree was reconstructed based on the reverse transcripase (RT) nucleotide sequences present within EFVs identified in *Xiphophorus*. ERV-F.1-Xeve-chr-15 is indicated with a red line. Colors indicate species from which EFVs were identified and match Fig. S10. Clades belonging to the same species or genus were collapsed, and bootstrap values at key nodes are indicated. **B)** The genomic organization of diverse representative foamy viruses is shown in comparison to ERV-F.1-Xeve-chr-15 (bottom).

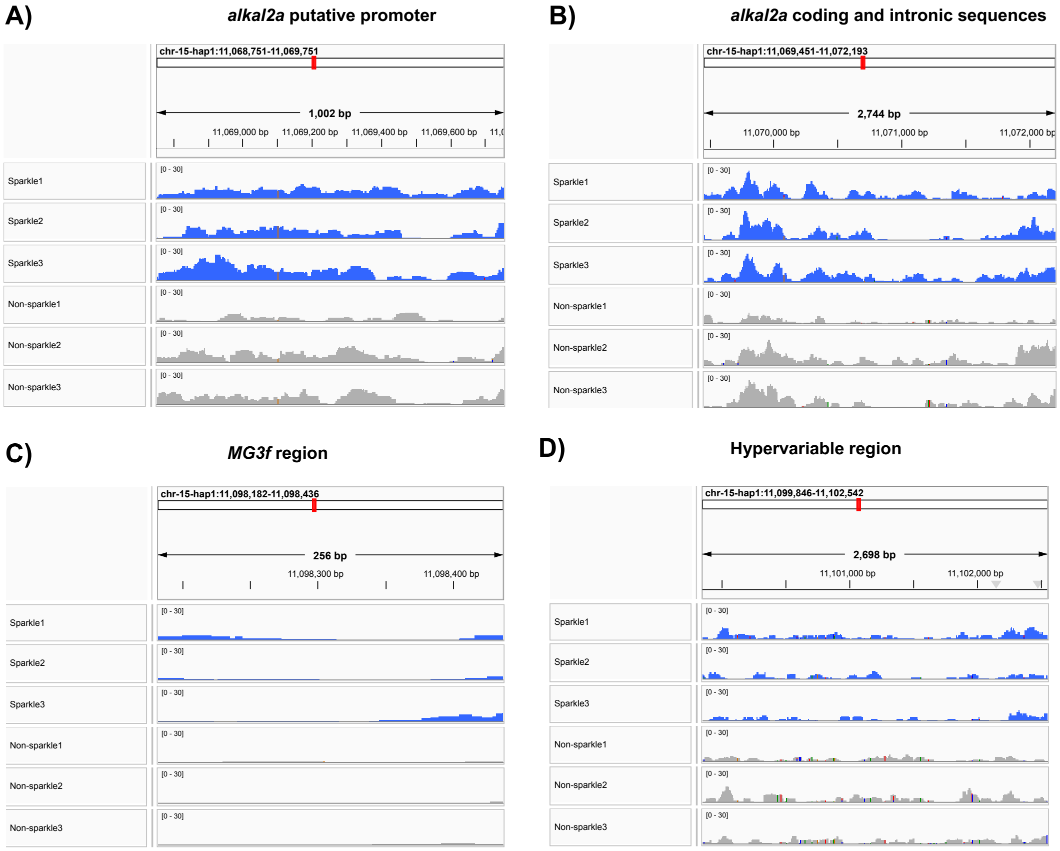


**Fig. S15.** Comparison of chromatin accessibility between sparkle (blue) and non-sparkle individuals (gray) in other regions of interest beyond the LTRs of ERV-F.1-Xeve-chr-15. Data was downsampled using samtools view for visualization such that all individuals had the same number of mapped reads genome-wide. **A)** Chromatin accessibility in all samples in the 1 kb region upstream of *alkal2a*, the putative promoter region of this gene. **B)** Chromatin accessibility across the coding and intronic regions of *alkal2a* in all samples. **C)** Chromatin accessibility within the *MG3f* insertion at the 3' edge of the ERV-F.1-Xeve-chr-15 insertion. **D)** Chromatin accessibility in the hypervariable region at the 3' edge of the GWAS peak.

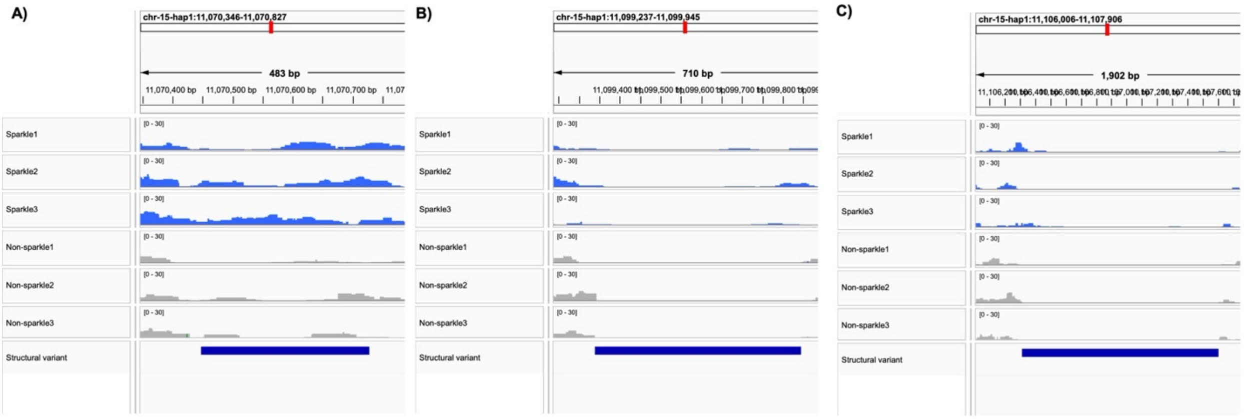


**Fig. S16.** Comparison of chromatin accessibility between sparkle (blue) and non-sparkle individuals (gray) at other structural variants that differ between sparkle and non-sparkle individuals in the GWAS interval besides ERV-F.1-Xeve-chr-15. Data were downsampled using samtools view for visualization such that all individuals had the same number of mapped reads genome-wide. **A-C)** Chromatin accessibility within other structurally variable regions shown in Fig. S27 that differ between sparkle and non-sparkle individuals and fall within or nearby the GWAS peak. Dark blue bar shows the location of the structural variants relative to the ATAC-seq data. **A)** Intronic 296 basepair deletion in an *alkal2a* intron, **B)** 509 basepair insertion nearby the ERV-F.1-Xeve-chr-15 insertion, and **C)** 1301 basepair insertion downstream from the ERV-F.1-Xeve-chr-15 insertion.

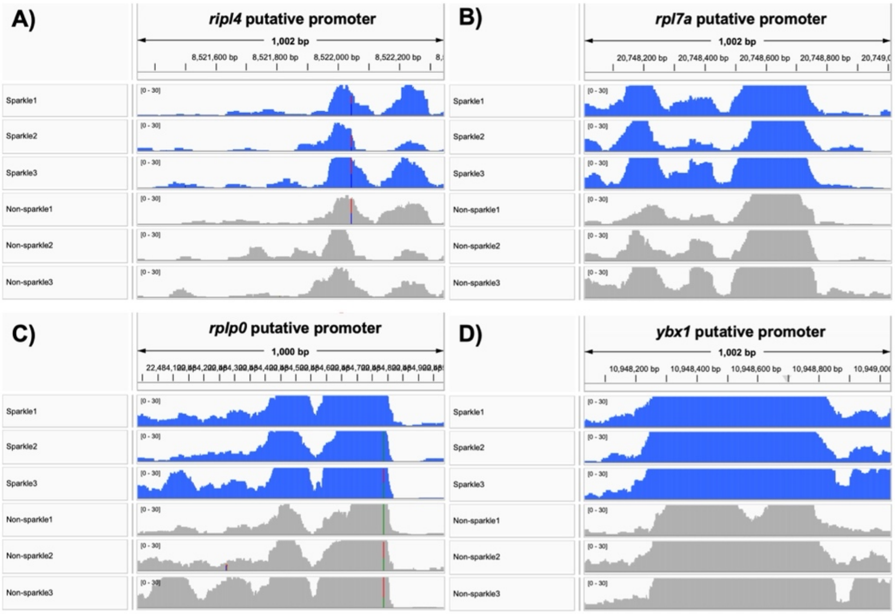


**Fig. S17.** Positive controls in ATAC-seq analysis including putative promoter regions of genes that were previously identified as ubiquitously highly expressed across tissues in *Xiphophorus* and with previous evidence of high chromatin accessibility (*72*). For each gene, accessibility is plotted for sparkle (blue) and non-sparkle (grey) individuals for the 1 kb region upstream of the transcriptional start site. Gene names are noted above each plot. Data were downsampled using samtools view for visualization such that all individuals had the same number of mapped reads genome-wide.

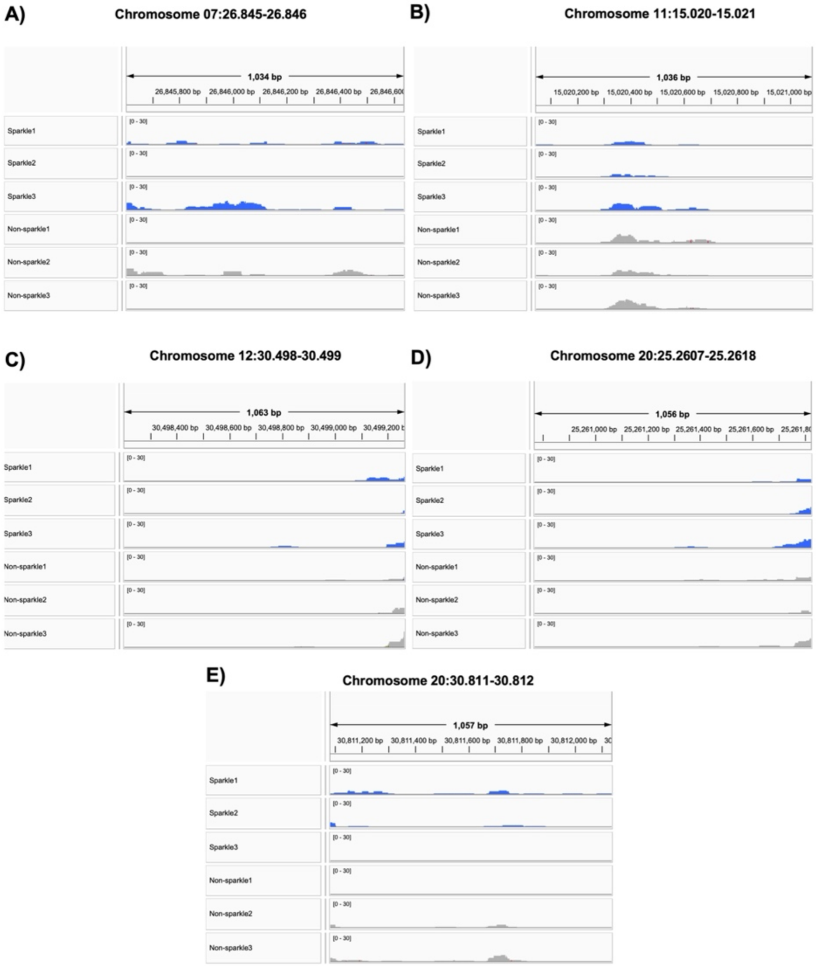


**Fig. S18.** Chromatin accessibility results for data from three sparkle (blue) and three non-sparkle (gray) individuals at a subset of "control" LTR regions with high sequence similarity to ERV-F.1-Xeve-chr-15. We identified 80 LTRs with high sequence identity (≥95% identical) to the LTRs of ERV-F.1-Xeve-chr-15 and that reached at least 60% of the length of the ERV-F.1-Xeve-chr-15 LTRs. The regions plotted here show all regions that exceeded a minimum coverage threshold (see Supplementary Materials 13). Data were downsampled using samtools view for visualization such that all individuals had the same number of mapped reads genome-wide.

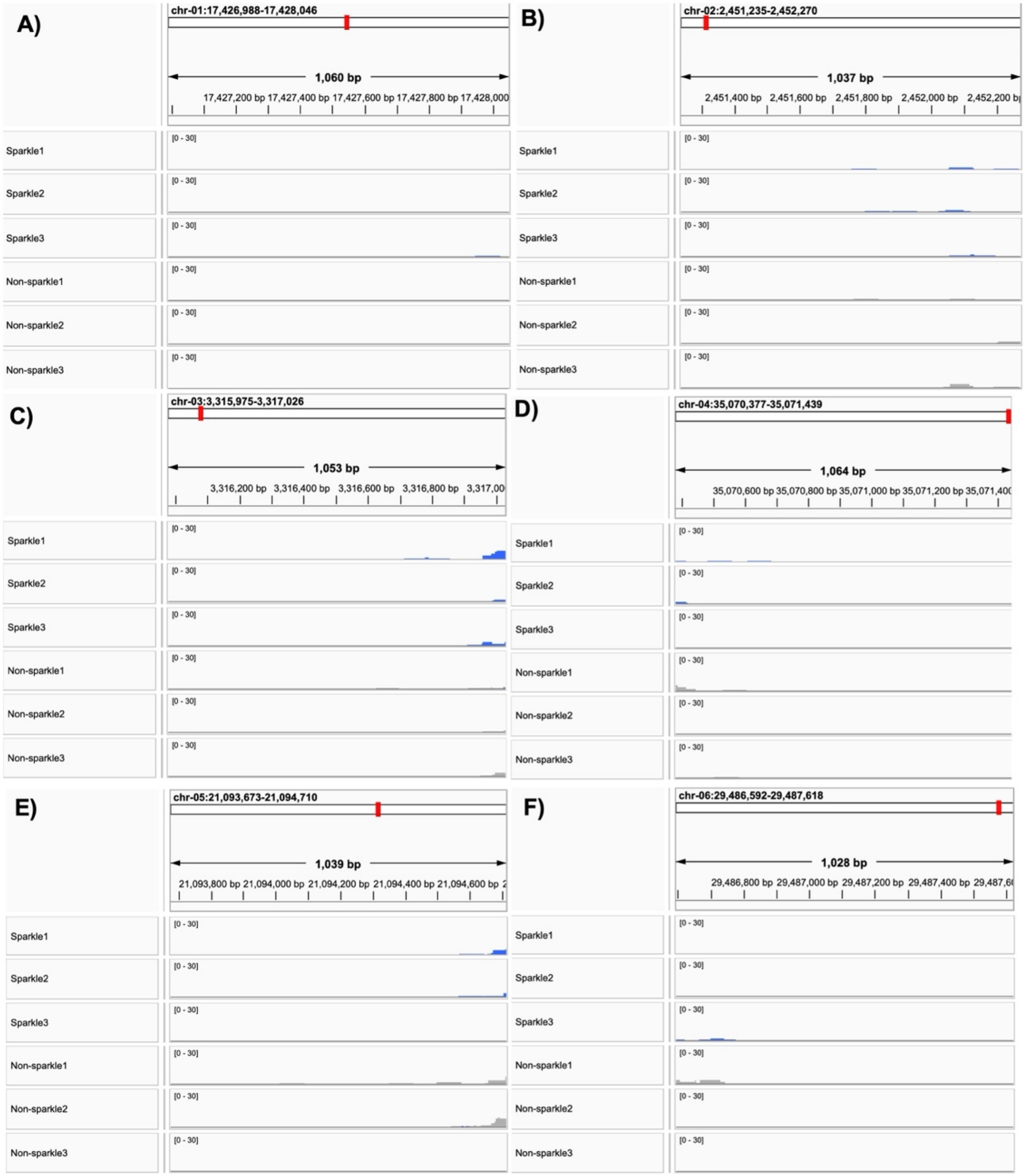


**Fig. S19.** Chromatin accessibility results for data from three sparkle (blue) and three non-sparkle (gray) individuals at a subset of "control" LTR regions with high sequence similarity to ERV-F.1-Xeve-chr-15. We identified 80 LTRs with high sequence identity (≥95% identical) to the LTRs of ERV-F.1-Xeve-chr-15 and that reached at least 60% of the length of the ERV-F.1-Xeve-chr-15 LTRs. The regions plotted here show a sample of the regions with low average coverage across the LTR length (90% of regions; see Supplementary Materials 13). Data were downsampled using samtools view for visualization such that all individuals had the same number of mapped reads genome-wide.

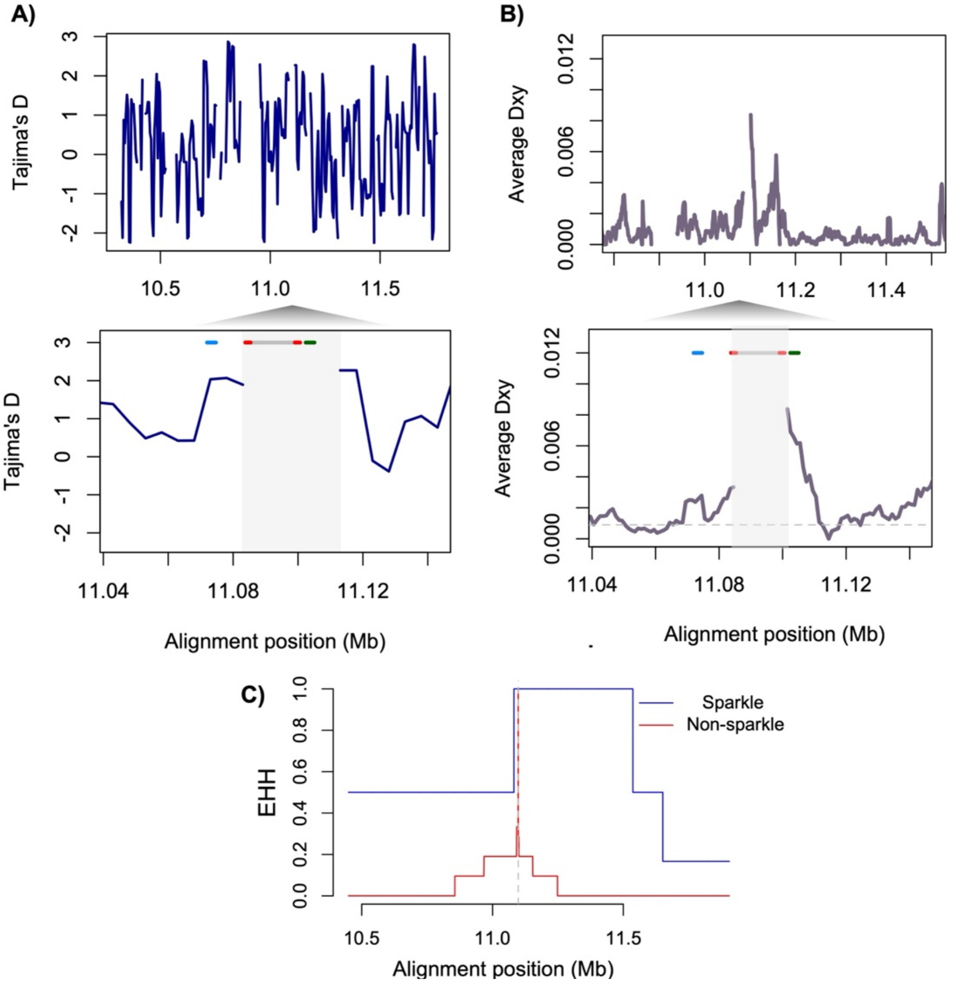


**Fig. S20.** Results of population genetic analyses of aligned haplotypes from assemblies derived from sparkle and non-sparkle individuals. **A)** Tajima's D is plotted in sliding 10 kb windows. Top plot shows variation in Tajima's D across the entire aligned region and bottom shows the region around the ERV-F.1-Xeve-chr-15 insertion, which is absent in all non-sparkle haplotypes. Windows with greater than 20% alignment gaps are excluded from the plot. The light blue box highlights the location of the ERV-F.1-Xeve-chr-15 insertion, where data is not present for non-sparkle individuals. Line segments at the top of the plot indicates the locations of *alkal2a* (blue), ERV-F.1-Xeve-chr-15 (gray and red—flanking LTRs), and the hypervariable region (green) in this alignment. **B)** Average pairwise sequence divergence (Dxy) between sparkle and non-sparkle haplotypes. Top plot shows Dxy across the entire aligned region and bottom shows the region around ERV-F.1-Xeve-chr-15, which is absent in all non-sparkle haplotypes. Line segments at the top of the plot indicates the locations of *alkal2a* (blue), ERV-F.1-Xeve-chr-15 (gray and red—flanking LTRs), and the hypervariable region (green) in this alignment. **C)** Results of extended haplotype homozygosity of an informative SNP within the sparkle haplotype relative to the non-sparkle associated SNP at that locus. The SNP was chosen from the hypervariable region since the insertion itself is absent in non-sparkle individuals. See Supplementary Materials 17 for details on analysis approach for results shown in **A-C**.

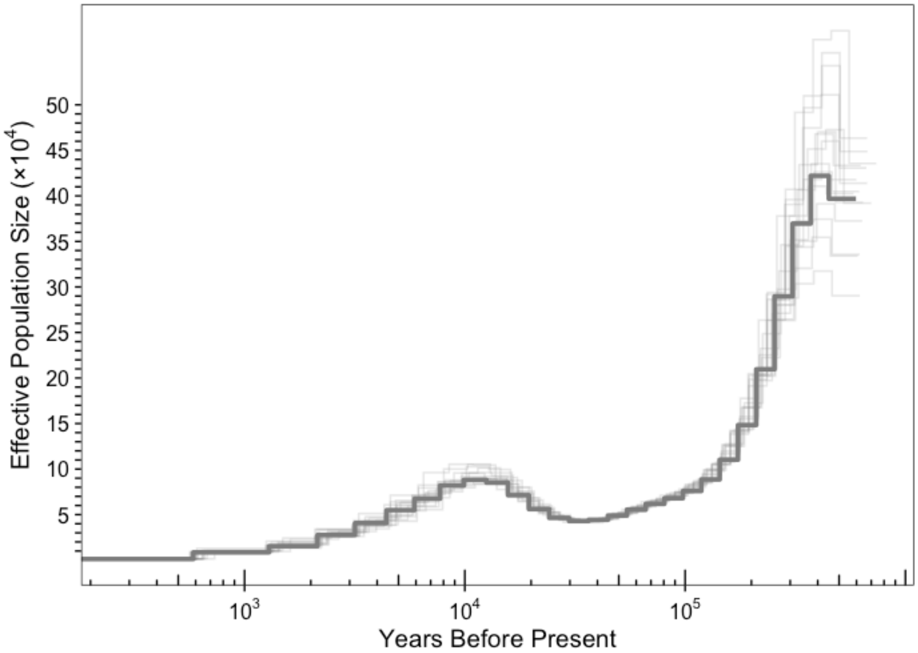


**Fig. S21.** Demographic history of the *X. evelynae* population inferred by PSMC analysis based on the *X. evelynae* reference genome generated for this project. The plotted results assume two generations per year and a mutation rate of $3.5 \times 10^{-9}$ per basepair per generation. Light gray lines indicate results from bootstrap replicates resampling the data and re-running PSMC analysis on resampled data.

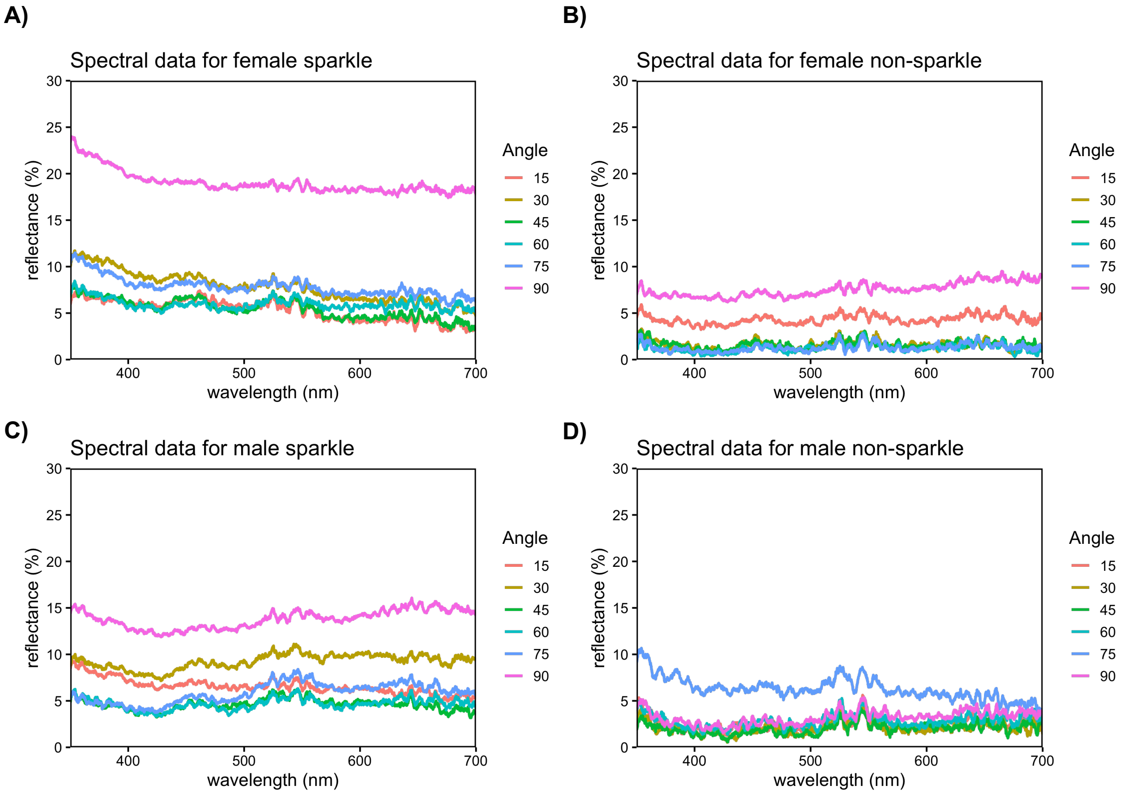


**Fig. S22.** Average of measured light reflectance at an incidence angle of 15°, 30°, 45°, 60°, 75°, and 90° in sparkle females (**A**), non-sparkle females (**B**), sparkle males (**C**), and non-sparkle males (**D**). See Supplementary Materials 19 for more details.

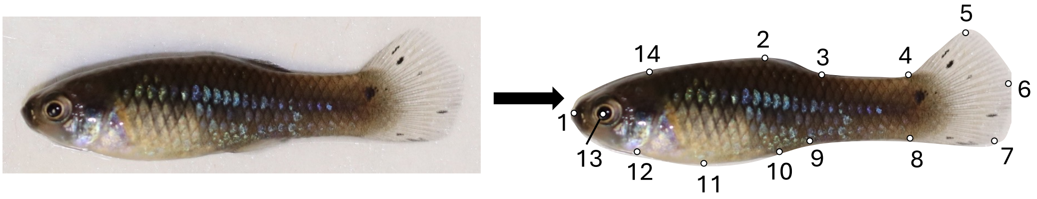


**Fig. S23.** Landmarks used in phenotyping analysis. These landmarks were used to align all adult fish photos to a reference fish for pattern analysis.

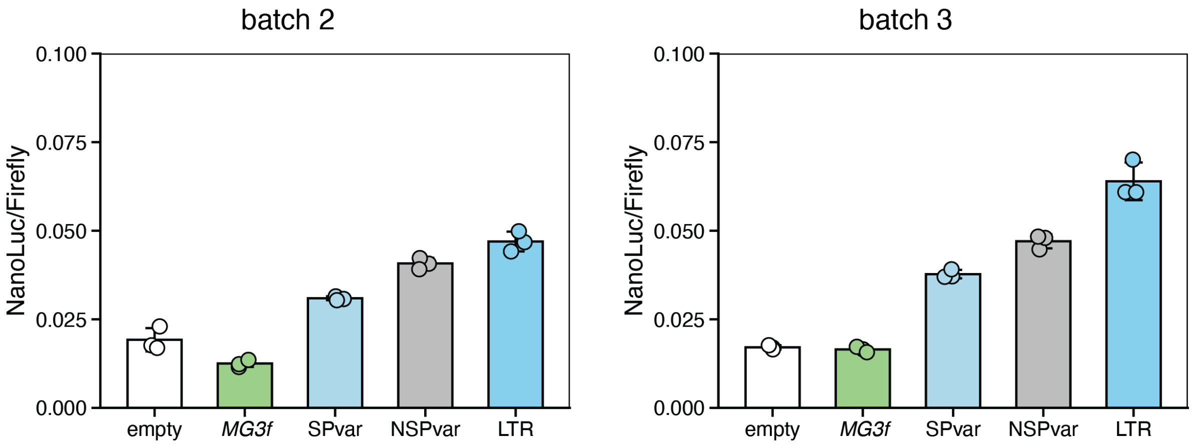


**Fig. S24.** Replication of results of the luciferase assay to investigate *cis*-regulatory potential. As reported in the main text, the LTR sequence acts as the strongest driver of reporter gene expression in HEK 293-T cells compared to an empty construct and other sequences of interest. Shown here are results from two replicate experiments performed on different days from the results shown in Fig. 3G to ensure that the difference in luciferase activity (NanoLuc/Firefly) driven by each construct was consistent. Each point shows the results of a different transfection and represents the average of three technical replicates. The two plots show the results from each replicate experiment. *MG3f* – *MG3f* region, SPvar – sparkle variable region, NSPvar – non-sparkle variable region, LTR – LTR derived from ERV-F.1-Xeve-chr-15 insertion. See Supplementary Materials 15 for more details.

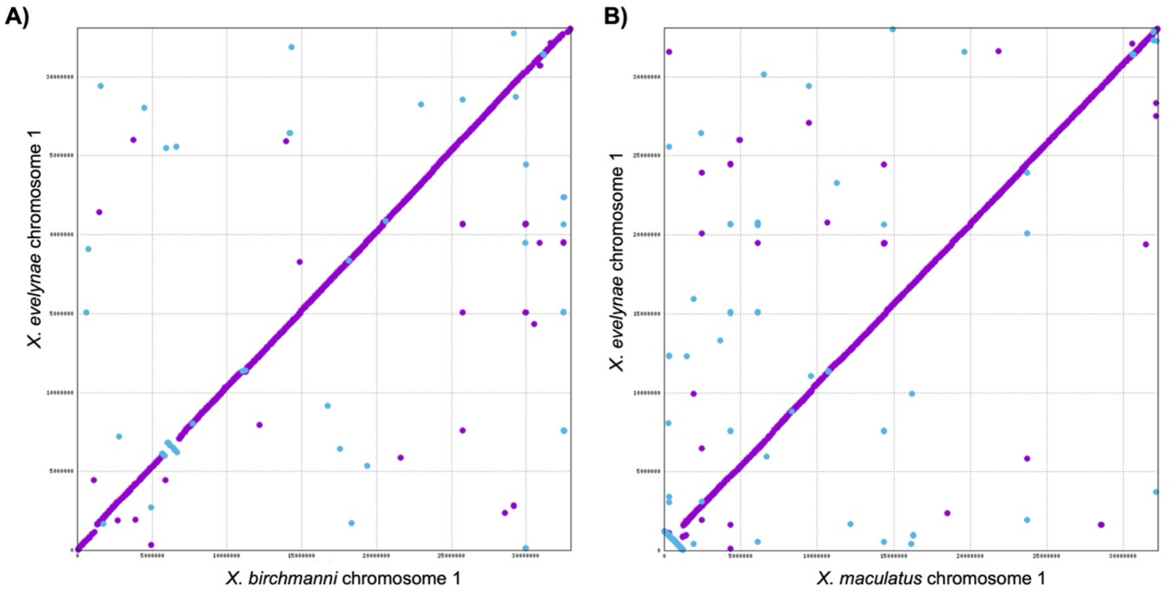


**Fig. S25.** Comparison of the *X. evelynae* genome assembly to *X. birchmanni* and *X. maculatus* using alignments with MUMmer4 (*23*). Shown here is an alignment of chromosome 1 between *X. birchmanni* and *X. evelynae* (**A**) and between *X. maculatus* and *X. evelynae* (**B**). While small rearrangements (or misassemblies) are evident in the off-diagonal blue lines, there is broad synteny along the chromosome. This pattern is mirrored in the majority of the 24 *Xiphophorus* chromosomes (see Fig. S26).

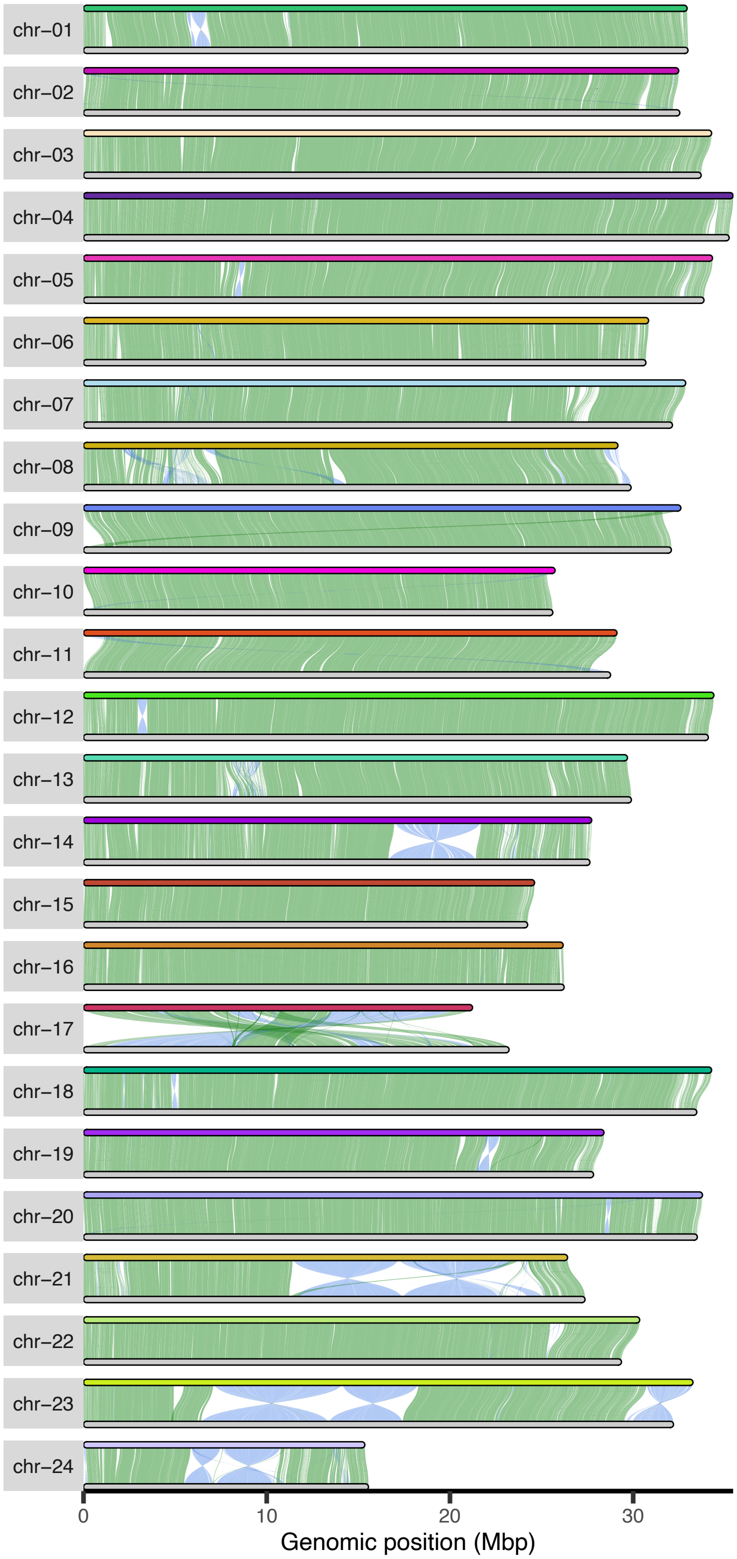


**Fig. S26.** Whole genome alignment of *X. evelynae* reference genome (top) relative to the *X. birchmanni* reference genome (bottom). Green lines show co-linear alignments and blue lines show inverted alignments.

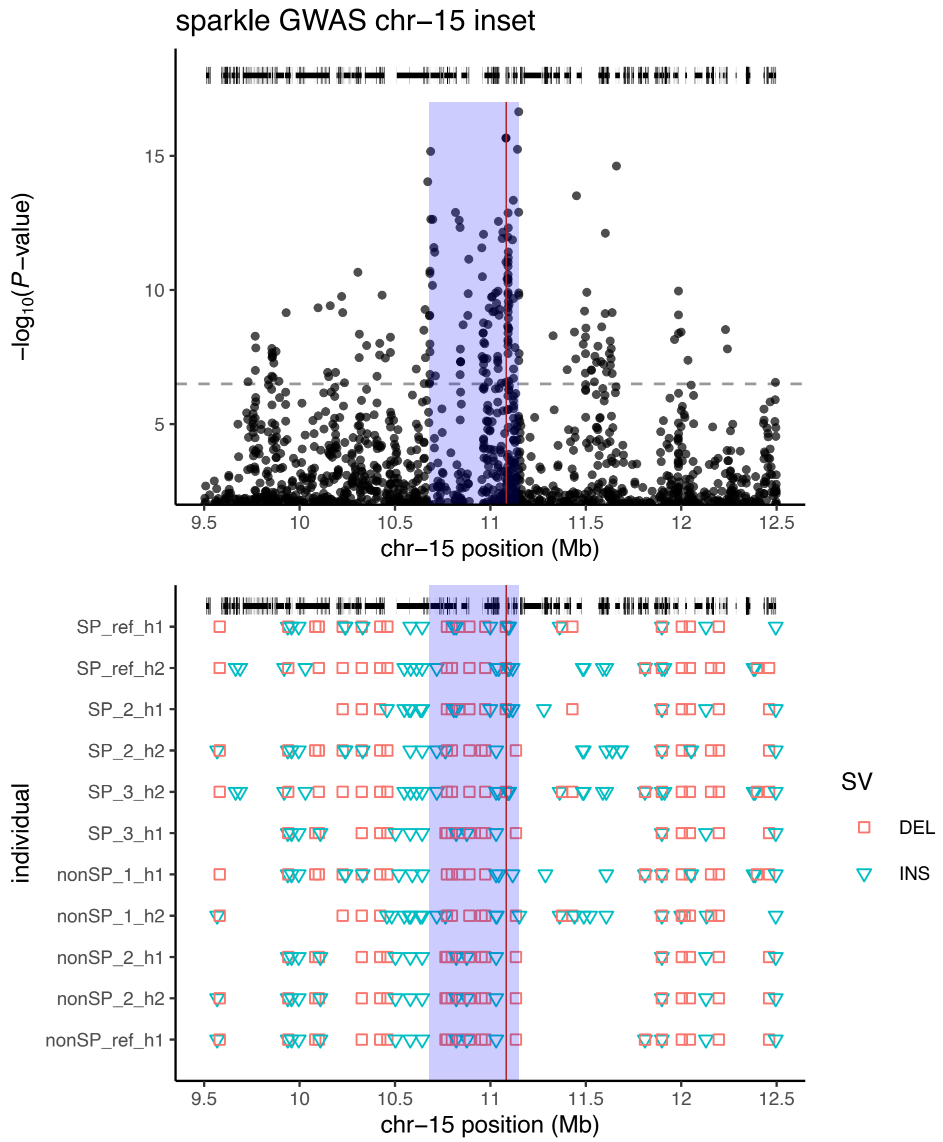


**Fig. S27.** Results of structural variant analysis using the PAV software (*24*) in long-read assemblies, focusing on regions within GWAS peak. The top plot shows the GWAS results for the focal region analyzed. The bottom plot shows structural variant calls for each individual haplotype. The labels on the y-axis of this plot indicate the phenotype and haplotype id for each analyzed haplotype (SP – sparkle individual, nonSP – non-sparkle individual; h1 – haplotype 1, h2 – haplotype 2). In the lower plot, blue triangles indicate detected insertion variants and red squares indicates detected deletion variants relative to the reference. In both plots, black lines and ticks show annotated gene models in the region. The red line indicates the location of *alkal2a*. The blue polygon spans the most strongly associated region in the GWAS (see Supplementary Materials 6).

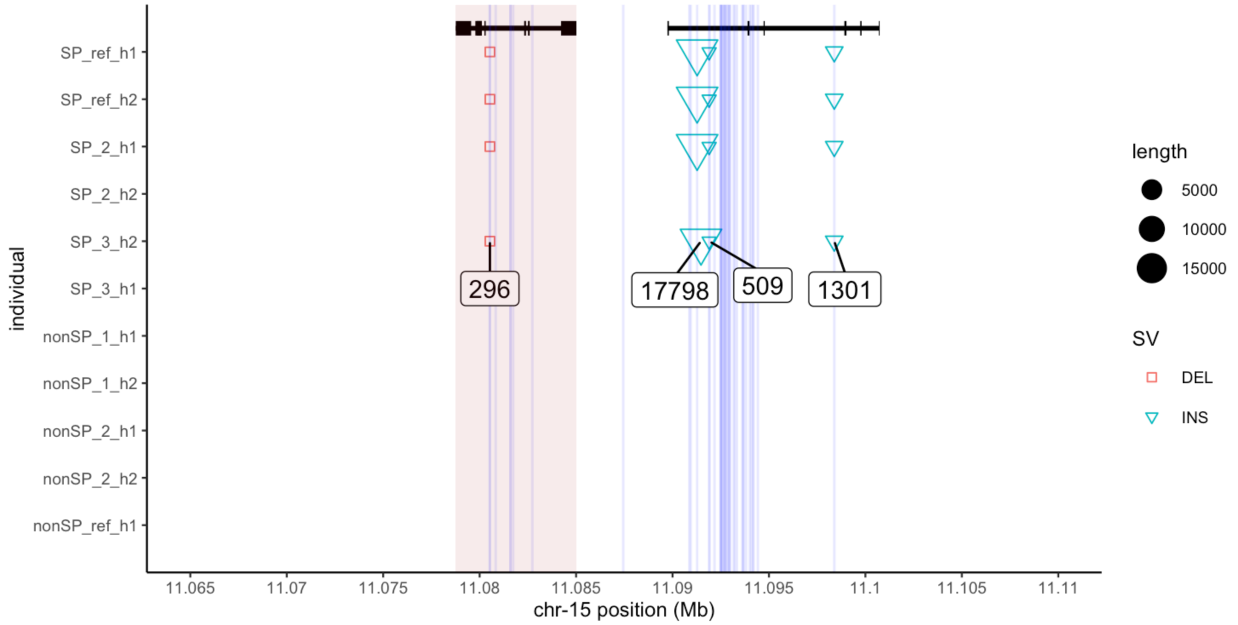


**Fig. S28.** Structural variants that co-segregate with phenotype in our analysis of haplotype phased assemblies with known phenotypes. The labels on the y-axis of this plot indicate the phenotype and haplotype number for each analyzed haplotype (SP – sparkle individual, nonSP – non-sparkle individual; h1 – haplotype 1, h2 – haplotype 2). Black lines show the gene models (*alkal2a* – red polygon, followed by *acp1*), red squares indicate deletions, blue triangles indicate insertions, and numbers represent indel lengths, including the ~17 kb ERV insertion. Blue lines highlight informative SNPs.

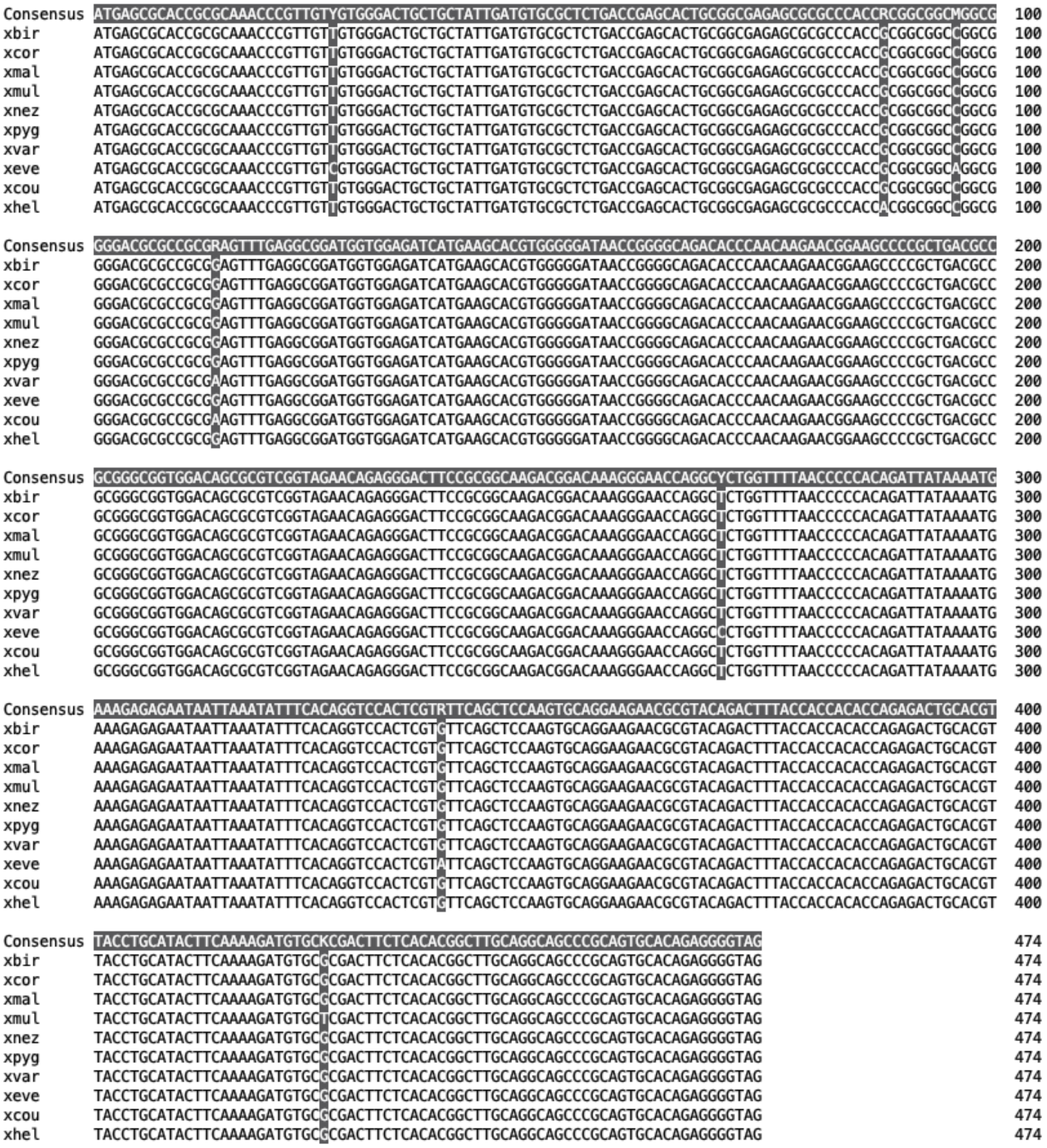

**Fig. S29.** Alignments of *alkal2a* cDNA sequence across available *Xiphophorus* species with annotated genome assemblies. Only one nonsynonymous substitution is observed in the gene across *Xiphophorus* (an A→T substitution at amino acid 30 in the *X. hellerii* reference).

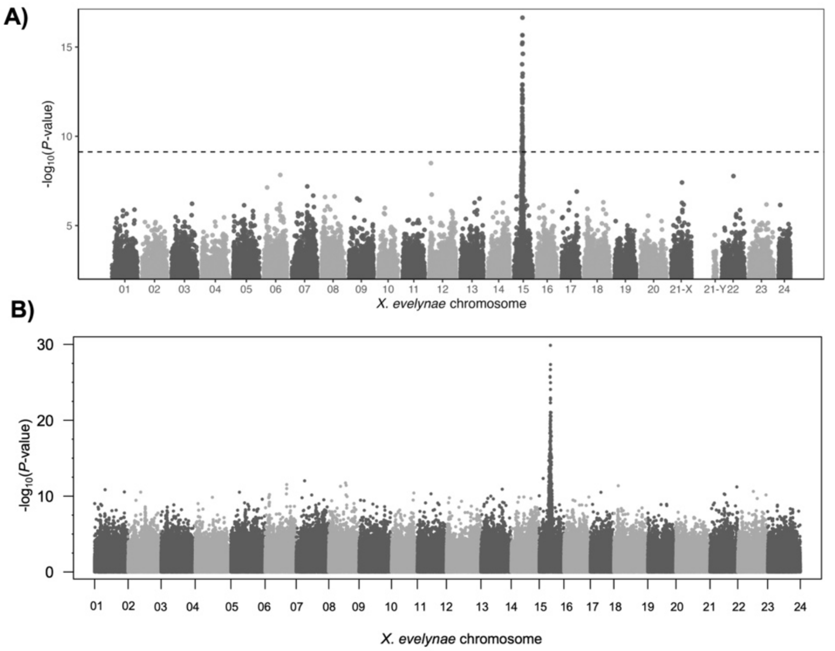


**Fig. S30.** GWAS results using a different reference genome or accounting for population structure in the analysis. **A)** GWAS analysis performed as described in Supplementary Materials 6 but using the non-sparkle reference genome instead of the sparkle reference genome. Dashed line shows the genome-wide significance threshold determined by permutations (see Supplementary Materials 6). **B)** GWAS analysis performed using pseudohaploid data and covariates to account for population structure. In order to include covariates in this low coverage dataset, we converted our low-coverage data to pseudohaploid calls, performed PCA in PLINK, and then included PCs 1-5 as covariates in a GWAS implemented in PLINK. Despite reduced power expected from a pseudohaploid analysis, we recover the chromosome 15 peak in this analysis. See Supplementary Materials 6 for more information.

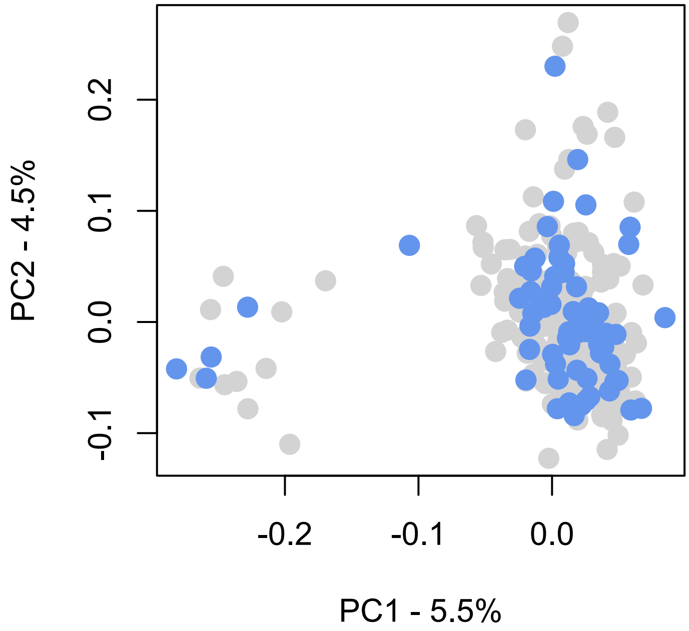


**Fig. S31.** No evidence of population structure as a function of phenotype in our GWAS dataset. PCA was performed on pseudohaploid data using PLINK; see Supplementary Materials 6 for more information. Blue points indicate sparkle individuals and gray points indicate non-sparkle individuals. While the analysis shown here plots PC1 versus PC2, we did not observe correlations between the trait and PCs 1-20 in our analyses.

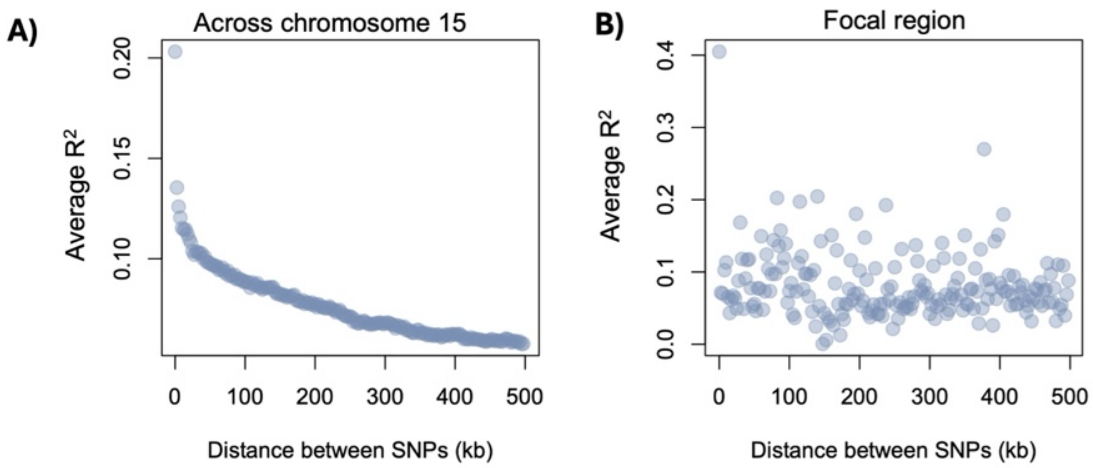


**Fig. S32.** Decay in linkage disequilibrium over physical distance based on the pseudohaploid data generated from GWAS dataset on average across chromosome 15 (**A**) and near the ERV insertion (**B**). Linkage disequilibrium was quantified using the $R^2$ metric and quantified in PLINK.

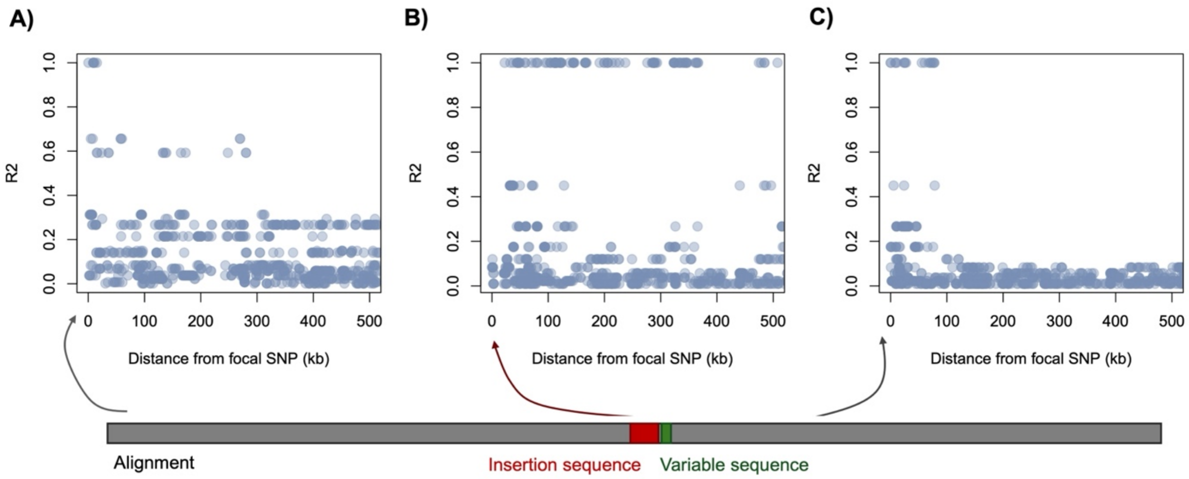


**Fig. S33.** Decay in linkage disequilibrium over physical distance based on alignment of 11 phased haplotypes from PacBio HiFi sequencing, using focal SNPs drawn from different regions of the alignment. **A**) Decay of linkage disequilibrium over physical distance from a SNP at 10.3 Mb on chromosome 15, **B**) Decay of linkage disequilibrium over physical distance from a SNP at 11.08 on chromosome 15, spanning the inserted region, and **C)** Decay of linkage disequilibrium over physical distance from a SNP at 11.13 Mb, after the inserted region. Linkage disequilibrium was calculated using the $R^2$ metric and implemented in PLINK.

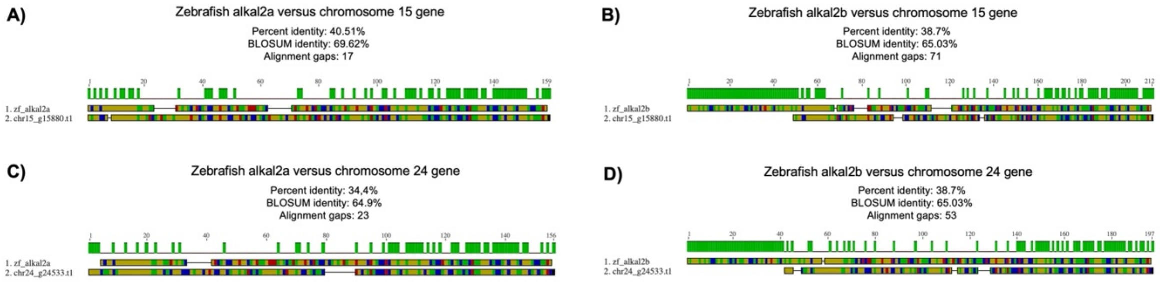


**Fig. S34.** Analysis of protein similarity of *Xiphophorus alkal* paralogs on chromosome 15 and chromosome 24 to zebrafish *alkal2a* and *alkal2b*. Alignments were visualized in Geneious. Green bar above the alignment visualization indicates residues that do not match in state between the two proteins and numbers correspond to alignment position. Text above the alignment indicates the comparison being made, the percent identity of amino acids across the two proteins, the BLOSUM percent similarity, and the number of gaps in the alignment. **A)** Alignment of zebrafish *alkal2a* to chromosome 15 ortholog. **B)** Alignment of zebrafish *alkal2b* to chromosome 15 ortholog. **C)** Alignment of zebrafish *alkal2a* to chromosome 24 ortholog. **D)** Alignment of zebrafish *alkal2b* to chromosome 24 ortholog.

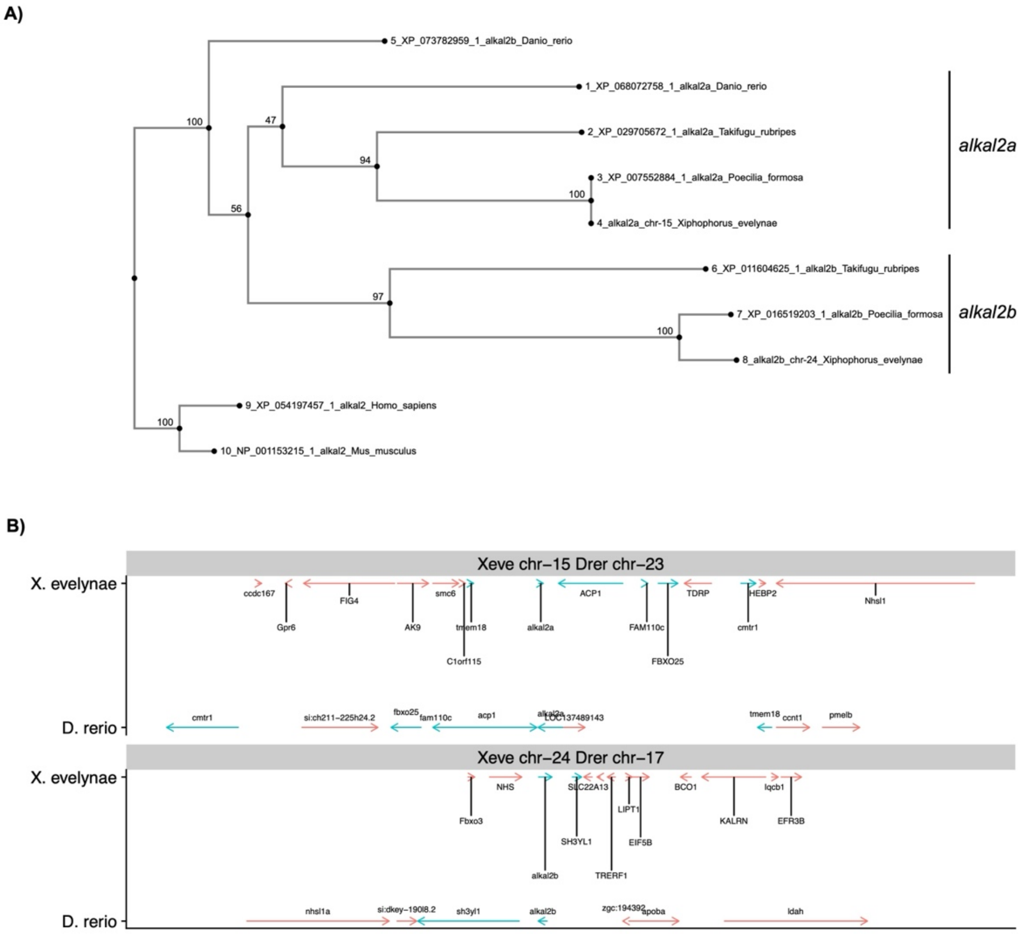


**Fig. S35.** Gene tree and microsynteny pattern analysis of *alkal2* paralogs. **A**) Gene tree of *alkal2a* and *alkal2b* in several species. The copy of *alkal2* on *X. evelynae* chromosome 15 groups with *D. rerio alkal2a*. **B**) Microsynteny pattern analysis near *alkal2* paralogs in *D. rerio* and *X. evelynae*. *D. rerio alkal2a* is flanked by the same up- and down-stream genes as the *alkal2* gene on chromosome 15 in *X. evelynae*, while *D. rerio alkal2b* and the *alkal2* gene in *X. evelynae* on chromosome 24 have lower evidence of microsynteny but share the closest gene (*sh3yl1*).

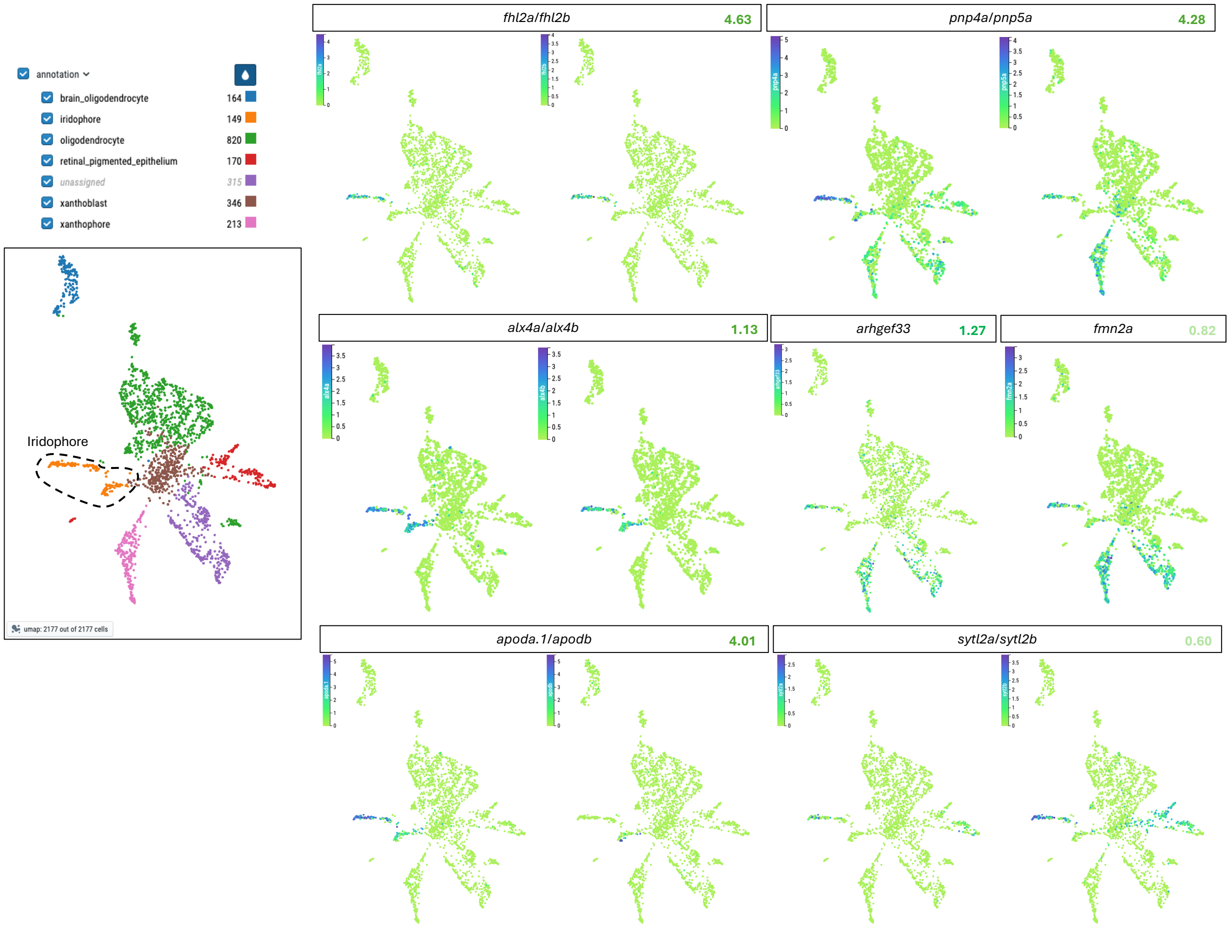


**Fig. S36.** Zebrafish single-cell database shows that several genes identified as upregulated in sparkle scales from our RNA-seq analysis are likely iridophore- and xanthophore-specific. Database analyzed was the Zebrahub database developed by the Chan-Zuckerberg biohub (https://zebrahub.sf.czbiohub.org/). Text above the plot lists the zebrafish gene name. Green numbers at the right of each zebrafish gene name denote the fold change value of the corresponding *X. evelynae* gene that is differentially expressed in our scale RNA-seq analysis. Inset to the left of the plot shows the Neural Crest lineage type, with iridophores in orange and circled with a dashed line.

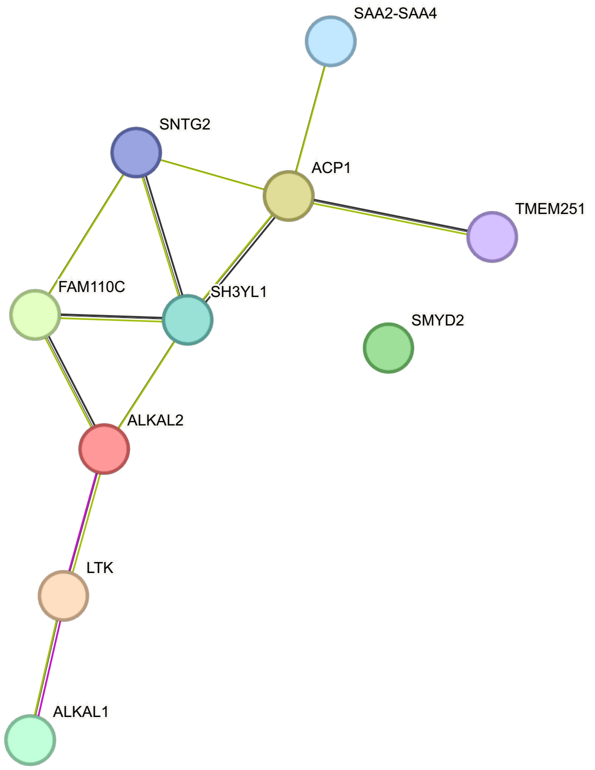


**Fig. S37.** STRING interaction network for ALKAL2 generated from the *Homo sapiens* database. This network highlights likely interactions between ALKAL2, FAM110C, and ACP1. Pink connections show experimentally verified interactions between proteins, purple shows interactions inferred from protein homology, green indicates interactions inferred from the 'gene neighborhood,' and black lines show interactions inferred by coexpression data. Notably, SMYD2, the only differentially expressed gene between sparkle and non-sparkle scales that localizes to the GWAS region apart from ALKAL2, is not inferred to be a part of this network.

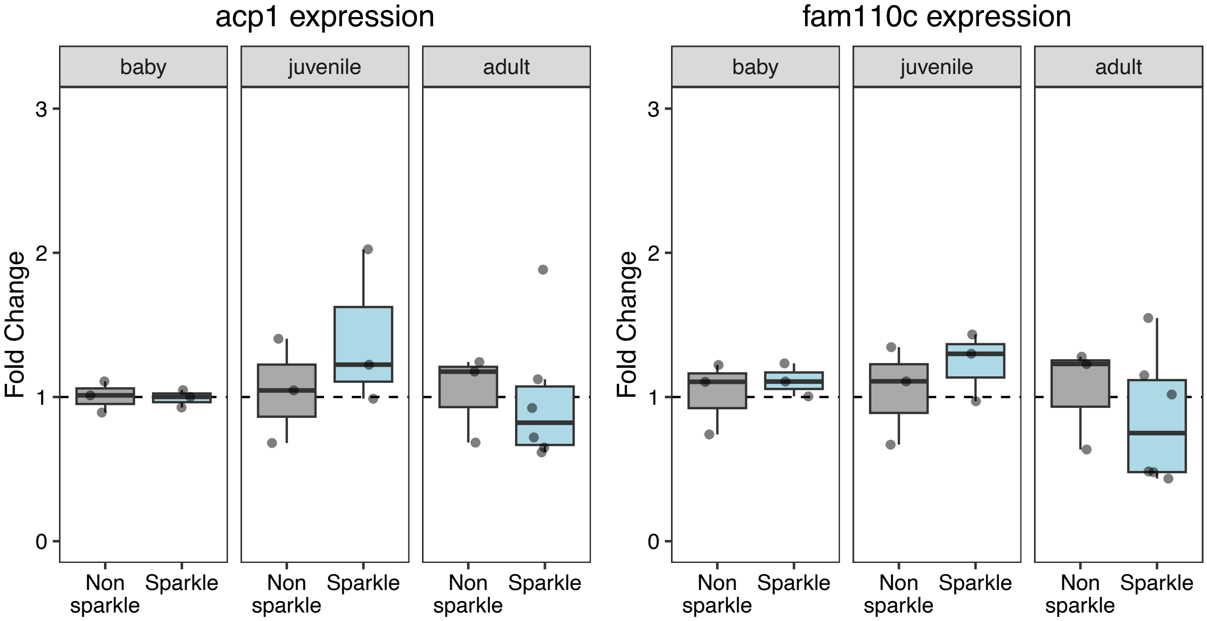


**Fig. S38.** qPCR results for *acp1* and *fam110c* for non-sparkle (grey) and sparkle (blue) individuals at different development stages. Fold change was calculated by the $2^{-\Delta\Delta Ct}$ method (with the focal gene normalized to *efa1-alpha* expression). Fry and juveniles were genotyped prior to qPCR to distinguish between individuals that had not yet developed the trait versus individuals that harbored non-sparkle haplotypes. We ran a Kruskall-Wallis test on the fold change between non-sparkle and sparkle individuals at each life stage for *acp1* (fry: p-value = 0.83; juvenile: p-value = 0.51; adult: p-value = 0.44) and for *fam110c* (fry: p-value = 0.51; juvenile: p-value = 0.51; adult: p-value = 0.30), finding no significant differences between phenotypes across all tested life stages.

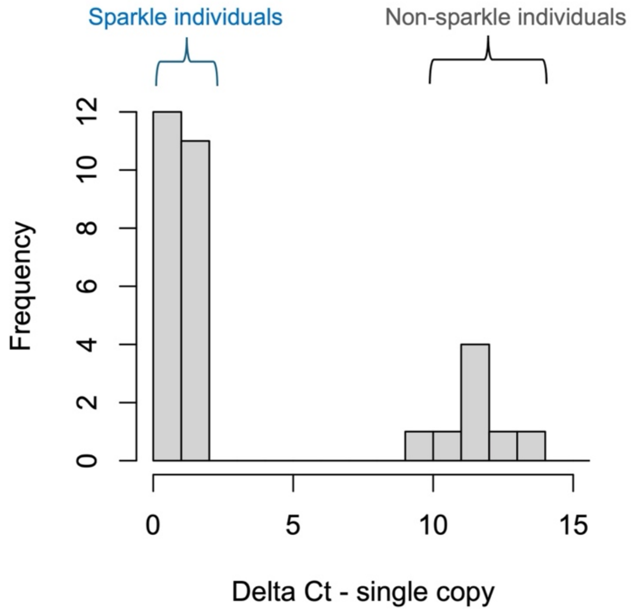


**Fig. S39.** Proof of principle for genomic DNA qPCR approach to quantify the presence or absence of the sparkle haplotype. Individuals with no sparkle haplotype have a large Delta Ct when measured relative to the single copy housekeeping gene *Nup43.* Individuals with one or two copies of the sparkle haplotype have a Delta Ct close to zero or one.

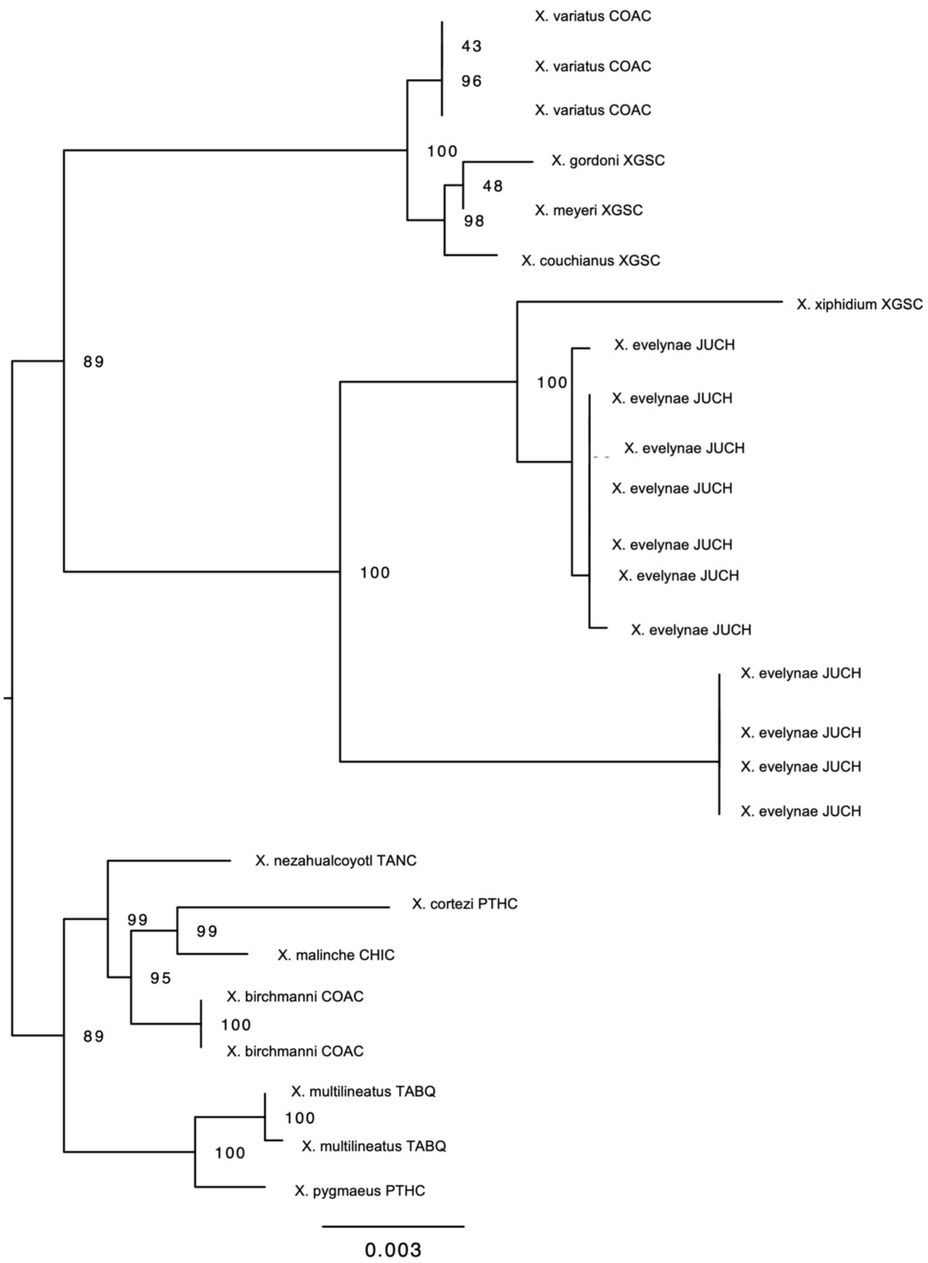


**Fig. S40.** Inferred phylogenetic relationships based on alignments of available long read assemblies in the hypervariable region identified between sparkle and non-sparkle individuals at the 3' end of the ERV insertion on chromosome 15. Relationships were inferred using RAxML, nodes list support from 100 rapid bootstraps. Nodes with bootstrap support less than 40 are not shown. Note that all non-sparkle haplotypes from the Juntas Chicas population cluster with the haplotype from *X. xiphidium.*

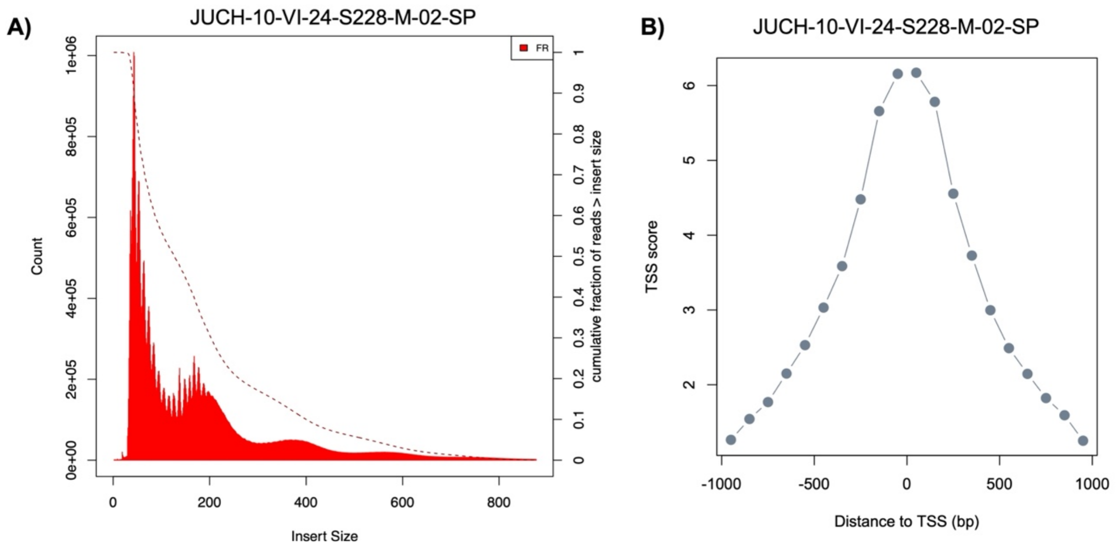


**Fig. S41.** Quality assessment of an example ATAC-seq sample with a sparkle phenotype (JUCH-10-VI-24-S228-M-02). **A**) Mapped read profile follows expectations for a successful ATAC-seq library. **B**) Enrichment of mapped fragments at the transcriptional start site (TSS) follows expectations for a successful ATAC-seq library.

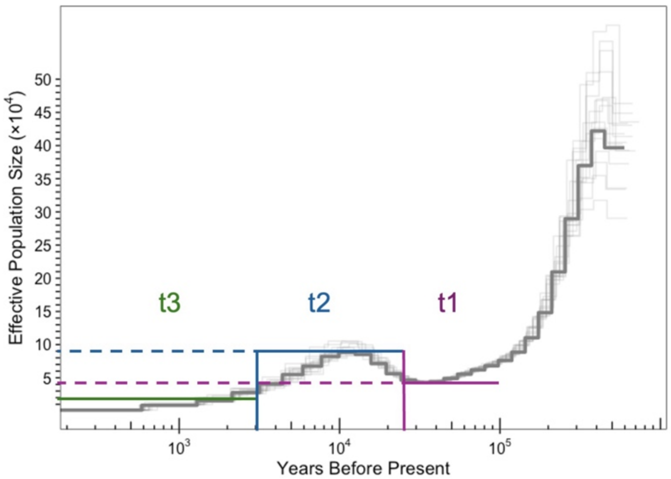


**Fig. S42.** Annotated PSMC history showing the step functions used to model change in population size in a subset of SLiM simulations. See Supplementary Materials 18 for more information.

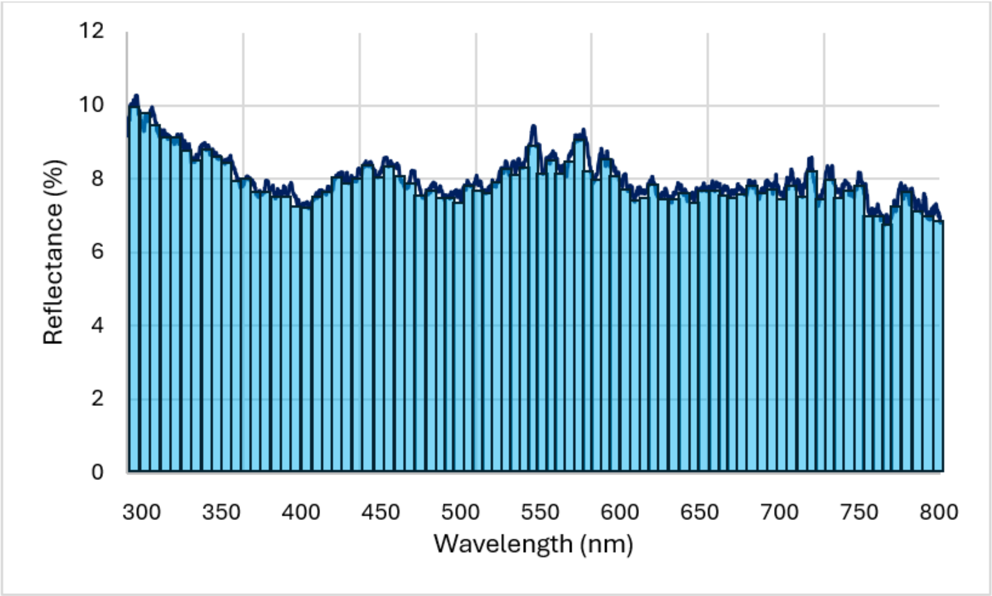


**Fig. S43.** Representation of the method used for measuring the brightness of the scales of *X. evelynae* fish. The brightness is measured as the area under the curve of the reflectance spectrum of the scales from the minimum to the maximum reflectance for each wavelength. See Supplementary Materials 19 for more information.

**Supplementary Tables**

**Table S1.** Genome assembly statistics for the primary *X. evelynae* sparkle assembly, excluding the Y chromosome.

| **Metric** | ***X. evelynae* sparkle genome (no Y)** |
|---|---|
| Assembly size | 715.8 Mb |
| Contig N50 | 30.9 Mb |
| Scaffold N50 | 32.2 Mb |
| Number of contigs | 72 |
| Number of scaffolds | 50 |
| Percent of sequence in 24 canonical chromosomes | 99.8% |
| Number of gaps | 22 |
| Number of chromosomes assembled telomere to telomere | 4 |
| Read N50 | 19,370 |
| Bases | 50,048,861,992 |
| Coverage | 70.0 |

**Table S2.** List of all genes in the GWAS associated region on chromosome 15. *Provided as an attached Excel file*.

**Table S3.** Structure of haplotype resolved assemblies at focal region on chromosome 15.
*Provided as an attached Excel file*.

**Table S4.** Genome assembly statistics for additional *X. evelynae* individuals with PacBio data collected for this project. Reported statistics exclude the Y chromosome for each genome.

| **Metric** | ***X. evelynae* non-sparkle reference genome** | ***X. evelynae* S178-M-01 h1** | ***X. evelynae* S178-M-01 h2** | ***X. evelynae* S250-M-01 h1** | ***X. evelynae* S250-M-01 h2** | ***X. evelynae* S250-M-02 h1** | ***X. evelynae* S250-M-02 h2** | ***X. evelynae* S265-M-01 h1** | ***X. evelynae* S265-M-01 h2** |
|---|---|---|---|---|---|---|---|---|---|
| Assembly size (Mb) | 716.5 | 676.4 | 675.9 | 694.6 | 688.3 | 700.6 | 706.5 | 695.7 | 676.1 |
| Contig N50 (Mb) | 30.6 Mb | 24.3 | 18.4 | 26.7 | 25.8 | 29.4 | 27.8 | 23.1 | 14.9 |
| Scaffold N50 (Mb) | 32.3 Mb | 29.7 | 29.1 | 30.1 | 29.7 | 32.0 | 31.1 | 30.7 | 29.9 |
| Number of contigs | 81 | 171 | 195 | 104 | 93 | 116 | 107 | 176 | 212 |
| Number of scaffolds | 65 | 37 | 31 | 44 | 38 | 69 | 58 | 47 | 41 |
| Percent of sequence in 24 chromosomes | 99.7% | 99.9 % | 99.9 % | 99.9 % | 99.9 % | 99.7 % | 99.7 % | 99.8 % | 99.8 % |
| Number of gaps | 16 | 134 | 164 | 60 | 55 | 47 | 49 | 129 | 171 |
| Number of chromosomes assembled telomere to telomere | 3 | 0 | 0 | 2 | 1 | 2 | 0 | 0 | 0 |
| Read N50 | 15,876 | 22,823 | | 18,942 | | 14,312 | | 15,904 | |
| Bases | 94,013,468,758 | 20,778,227,175 | | 34,725,471,458 | | 26,961,595,670 | | 19,565,662,492 | |
| Coverage | 131.5 | 29.0 | | 48.6 | | 37.71 | | 27.4 | |

**Table S5.** Results of bulk RNA-seq analysis of expression profiles in sparkle and non-sparkle scales using DESeq2. *Provided as an attached excel file*.

**Table S6.** Zebrafish knockout database phenotypes associated with each of the differentially regulated genes detected in our RNA-seq analysis. *Provided as an attached excel file.*

**Table S7.** Primer sequences used for qPCR and PCR. Primers used for qPCR include an efficiency estimate. Lower case sequences indicate overhang sequence for cloning for luciferase assays.

| **Primer target** | **Forward primer (5' to 3')** | **Reverse primer (5' to 3')** | **Efficiency** |
|---|---|---|---|
| *Alkal2a* cDNA | CACAGGTCCACTCGTGTTCA | GCAGGTAACGTGCAGTCTCT | 99.8% |
| *Efa1-alpha* cDNA | CAGGCCCGTTTTGAGGAGAT | AGAGTGGTTCCGTTGGCATT | 101.1% |
| *Acp1* cDNA | TCGACAGGTGACCAAAGATGACT | TTCAAAGTCTGCATCGCTTCC | 109.7% |
| *Fam110c* cDNA | CAACAGAGGACGACAGCGAA | TCCCTCCCCAAGGTCTCAAT | 101.7% |
| *MG3f* gDNA | ggcttaGGTACCCAAAGCAACCCACAGCAGAG | ggtcagCTCGAGACCTACGATCCAACAACCCC | NA |
| Variable region primers gDNA | ggcttaGGTACCCAGGCCTGCAGCTAAAGTGA | ggtcagCTCGAGAGCTATAACTCACTCCCCGGT | NA |
| Unique sparkle locus gDNA | GAAGTAGTGAGGAGTGGGCG | GGGGGTTTATGGGACTGCTT | NA |

**Table S8.** Genomes searched for analysis of endogenous retrovirus insertions. *Provided as an attached excel file*.

**Table S9.** Coordinates of complete ERV-foamy-Xeve elements beyond chromosome 15 detected in phased haplotypes generated for this project. *Provided as an attached excel file*.

**Table S10**. Results of scan of 5' LTR region on Xeve-F.1-Xeve-chr-15 for JASPAR 2022
